# Validation of human telomere length trans-ancestry meta-analysis association signals identifies *POP5* and *KBTBD6* as novel human telomere length regulation genes

**DOI:** 10.1101/2023.07.12.548702

**Authors:** Rebecca Keener, Surya Chhetri, Carla J. Connelly, Margaret A. Taub, Matthew P. Conomos, Joshua Weinstock, Bohan Ni, Benjamin Strober, Stella Aslibekyan, Paul L. Auer, Lucas Barwick, Lewis C. Becker, John Blangero, Eugene R. Bleecker, Jennifer A. Brody, Brian E. Cade, Juan C. Celedon, Yi-Cheng Chang, L. Adrienne Cupples, Brian Custer, Barry I. Freedman, Mark T. Gladwin, Susan R. Heckbert, Lifang Hou, Marguerite R. Irvin, Carmen R. Isasi, Jill M. Johnsen, Eimear E. Kenny, Charles Kooperberg, Ryan L. Minster, Sergei Nekhai, Nathan Pankratz, Patricia A. Peyser, Kent D. Taylor, Marilyn J. Telen, Baojun Wu, Lisa R. Yanek, Ivana V. Yang, Christine Albert, Donna K. Arnett, Allison E. Ashley-Koch, Kathleen C. Barnes, Joshua C. Bis, Thomas W. Blackwell, Eric Boerwinkle, Esteban G. Burchard, April P. Carson, Zhanghua Chen, Yii-Der Ida Chen, Dawood Darbar, Mariza de Andrade, Patrick T. Ellinor, Myriam Fornage, Bruce D. Gelb, Frank D. Gilliland, Jiang He, Talat Islam, Stefan Kaab, Sharon L.R. Kardia, Shannon Kelly, Barbara A. Konkle, Rajesh Kumar, Ruth J.F. Loos, Fernando D. Martinez, Stephen T. McGarvey, Deborah A. Meyers, Braxton D. Mitchell, Courtney G. Montgomery, Kari E. North, Nicholette D. Palmer, Juan M. Peralta, Benjamin A. Raby, Susan Redline, Stephen S. Rich, Daniel Roden, Jerome I. Rotter, Ingo Ruczinski, David Schwartz, Rank Sciurba, M. Benjamin Shoemaker, Edwin K. Silverman, Moritz F. Sinner, Nicholas L. Smith, Albert V. Smith, Hemant K. Tiwari, Ramachandran S. Vasan, Scott T. Weiss, L. Keoki Williams, Yingze Zhang, Elad Ziv, Laura M. Raffield, Alexander P. Reiner, NHLBI Trans-Omics for Precision Medicine (TOPMed) Consortium, TOPMed Hematology and Hemostasis Working Group, TOPMed Structural Variation Working Group, Marios Arvanitis, Carol W. Greider, Rasika A. Mathias, Alexis Battle

## Abstract

Telomere length genome-wide association studies (GWAS) have become well-powered to detect novel genes in telomere length regulation. However, no prior work has validated these putative novel genes to confirm the contribution of GWAS loci to telomere length regulation. We conducted a trans-ancestry meta-analysis of 211,369 individuals. Through enrichment analyses of chromatin state and cell-type heritability we identified blood and immune cells as the most relevant cell type to examine telomere length association signals. We validated specific GWAS associations by overexpressing *KBTBD6*, a component of an E3 ubiquitin ligase complex, and *POP5*, a component of the Ribonuclease P/MRP complex, and demonstrating that both lengthened telomeres as predicted by our statistical analyses. CRISPR/Cas9 deletion of the predicted causal regions of these association peaks in K562 immortalized blood cells reduced expression of these genes, demonstrating that these loci are related to transcriptional regulation of *KBTBD6* and *POP5*, respectively. Together our results demonstrate the utility of telomere length GWAS in the identification of novel telomere length regulation mechanisms and highlight the importance of the proteasome-ubiquitin pathway in telomere length regulation.

## Introduction

Telomeres shorten with age and short telomeres are associated with several age-related diseases including bone marrow failure and immunodeficiency (Stanley and Armanios 2015). Individuals with these Short Telomere Syndromes have rare variants with large effects on telomere length regulation genes. Identification of causal variants in short telomere syndrome patients has led to the discovery of several genes we now appreciate as core telomere length regulation genes including *DKC1*, *NAF1*, *PARN*, and *ZCCHC8* (Alder et al. 2013; Stuart et al. 2015; Gable et al. 2019). Rare and common variants highlight the same set of core genes for many complex traits (Weiner et al. 2023), therefore a genome-wide association study (GWAS) on telomere length could feasibly be used to discover additional critical telomere length regulation genes. Despite the fact that 19 GWAS on leukocyte telomere length have been published (M. Mangino et al. 2009; Codd et al. 2010; Levy et al. 2010; Gu et al. 2011; Prescott et al. 2011; Massimo Mangino et al. 2012; Codd et al. 2013; J. H. Lee et al. 2013; Pooley et al. 2013; Liu et al. 2014; Saxena et al. 2014; Walsh et al. 2014; Massimo Mangino et al. 2015; Delgado et al. 2018; Zeiger et al. 2018; Dorajoo et al. 2019; C. Li et al. 2020; Codd et al. 2021; Taub et al. 2022), identifying 143 loci associated with telomere length, very little has been done to validate these signals representing new facets of telomere length regulation.

A key challenge facing interpretation of telomere length GWAS signals is accurately identifying causal genes driving the association signals. The vast majority of GWAS signals, including telomere length GWAS loci, are in non-coding regions, making it difficult to determine the likely causal gene (Maurano et al. 2012). Some telomere length GWAS have used colocalization analysis, statistically comparing GWAS signal to quantitative trait locus (QTL) data, to support shared causal signal with putative target genes (C. Li et al. 2020; Codd et al. 2021; Taub et al. 2022). Each of these were limited to expression QTLs (eQTLs) highlighting transcriptional regulatory genetic effects, but additional mechanisms may be involved, including alternative splicing revealed by splicing QTLs (sQTLs) (Y. I. Li et al. 2016). Furthermore, colocalization evidence does not confirm causal genes or relevant cell types. Such conclusions require functional validation of genetic regulatory and gene mechanism impacting telomere length, which were not explored in prior telomere length GWAS.

A second barrier to capitalizing on telomere length GWAS associated loci is that many of the associated loci are often in or near genes with no prior known direct effect on telomere length, making it difficult to understand the value in characterizing the underlying molecular mechanisms. Indeed, many of these association signals likely represent peripheral genes with indirect mechanisms on telomere length regulation (Boyle, Li, and Pritchard 2017). This is consistent with observations from screens assaying the effect of knock-out libraries in *Saccharomyces cerevisiae* (*S. cerevisiae*) on telomere length which identified genes involved in diverse pathways either lengthening or shortening telomeres (Askree et al. 2004; Gatbonton et al. 2006). Similarly, immunoprecipitation followed by mass spectrometry of *S. cerevisiae* telomerase components identified interactions with proteins with diverse functions (Askree et al. 2004; Gatbonton et al. 2006; Lin et al. 2015). In both types of experiments, the majority of the results were interpreted to indirect mechanisms on telomere length regulation. However, validation of genes identified in these studies has also identified direct effects on telomerase (Maicher et al. 2017; Laterreur et al. 2018).

Here, we leveraged four telomere length GWAS that used non-overlapping cohorts in a random-effects trans-ancestry meta-analysis on 211,369 individuals to identify 56 loci associated with human telomere length. Using stratified linkage disequilibrium score regression (S-LDSC) (Finucane et al. 2015) and enrichment analysis of Roadmap Epigenomics chromatin data (Roadmap Epigenomics Consortium et al. 2015) we determined that blood and immune cells were the most relevant cell type for telomere length association signals. We validated some of our colocalization analysis results in cultured cells and demonstrated that overexpression of *KBTBD6* and *POP5* increased telomere length as predicted by our statistical analyses. CRISPR/Cas9 deletion of the predicted causal regions for signals attributed to these genes in immortalized blood cells reduced expression of both genes, further supporting the conclusion that *KBTBD6* and *POP5* are the causal genes at these telomere length association signals. Together this work shows the utility of human telomere length GWAS in identifying new aspects of telomere biology.

## Results

### Trans-ancestry meta-analysis of leukocyte telomere length identifies 7 novel signals

We leveraged four GWAS with non-overlapping cohorts in a trans-ancestry meta-analysis of 211,379 individuals. Three studies were homogenous ancestries of European (C. Li et al. 2020), Singaporean Chinese (Dorajoo et al. 2019), or Bangladeshi (Delgado et al. 2018) individuals. The fourth study used HARE (Fang et al. 2019) to broadly categorize individuals as European, African, Asian, or Hispanic/Latino and generated ancestry-specific summary statistics (Taub et al. 2022)(Supplementary Table 1). We meta-analyzed these seven sets of summary statistics and broadly refer to the Asian, Singaporean Chinese, and Bangladeshi individuals as Asian in this manuscript (Figure 1). Across the four studies telomere length was estimated from blood leukocytes computationally from whole genome sequencing data using TelSeq (Taub et al. 2022) or experimentally using qPCR or a Luminex-based platform (Delgado et al. 2018; Dorajoo et al. 2019; C. Li et al. 2020). These studies previously demonstrated that all three assays are well correlated with telomere Southern blots. We used a random-effects model to identify 56 genome-wide significant loci (p-value < 5×10^-8^) including seven novel signals (Figure 1, Supplementary Table 2, Methods). Loci were considered novel if there were no other reported sentinels within 1 Mb of the lead single nucleotide polymorphism (SNP) at the locus.

**Figure 1:**
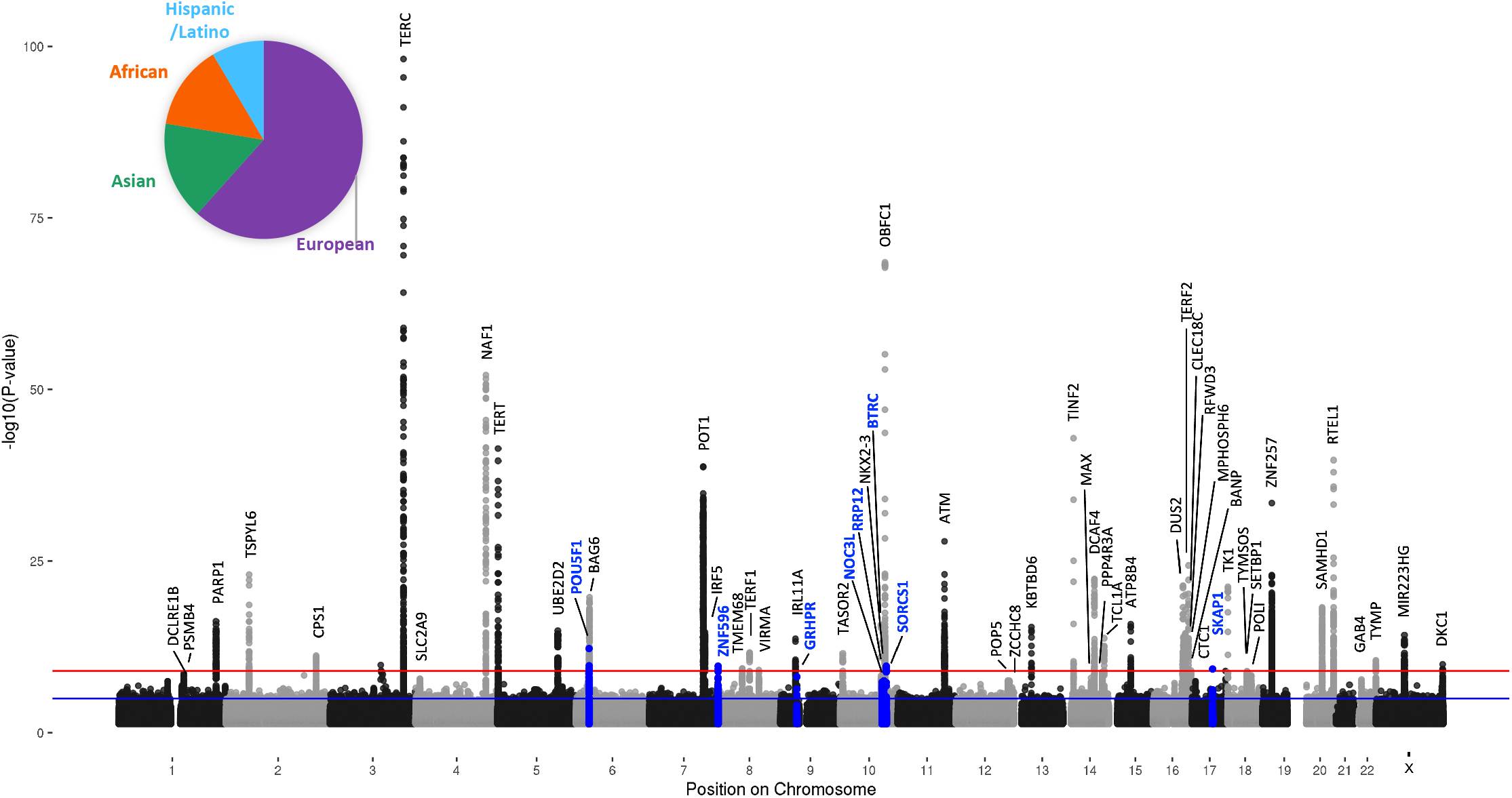
Trans-ancestry meta-analysis of leukocyte telomere length identifies 7 novel signals. Manhattan plot showing the results from the meta-analysis. The novel signals are shown in blue. The inset pie chart displays the proportion of different ancestries used in the meta-analysis.

### Fine-mapping analyses nominate putative causal variants and genes affecting telomere length

#### Colocalization analysis suggests genes underlying association signals

We used colocalization analysis (Giambartolomei et al. 2014) to determine whether each of our GWAS signals overlapped a signal from an independent quantitative trait locus (QTL) dataset (Methods), indicating causal genetic variants shared between telomere length and gene regulation. We began by examining large-scale expression quantitative trait locus (eQTL) and splicing quantitative trait locus (sQTL) datasets from diverse cellular contexts. Each GWAS included in our meta-analysis estimated telomere length from leukocytes extracted from whole blood. However, strong QTLs are often shared across cellular contexts (GTEx Consortium 2020) and telomere length is correlated across GTEx tissues (Demanelis et al. 2020); therefore, we included all 49 GTEx v8 tissues in our colocalization analysis. We found that 32 of 56 meta-analysis signals strongly colocalized (PPH4 > 0.7) with at least one eQTL or sQTL in at least one tissue (Supplementary Figure 1A,B,E). 12 signals colocalized (PPH4 > 0.7) with an eQTL or sQTL across more than five tissues and there was colocalization (PPH4 > 0.7) of at least one meta-analysis signal with at least one eQTL or sQTL in 45 out of 49 GTEx tissues (Supplementary Tables 3-4). We also conducted colocalization analysis using eQTLGen eQTLs (Võsa et al. 2021) and DICE eQTLs (Schmiedel et al. 2018; Võsa et al. 2021) (Supplementary Tables 5-6). eQTLGen increases power, with 31,685 individuals compared to GTEx whole blood with 755 individuals. DICE introduces cell type specificity, with eQTLs called from RNA-seq on 13 sorted blood and immune cell types, in 91 individuals. 11 of our signals colocalized (PPH4 > 0.7) with eQTLGen eQTLs (Supplementary Figure 1C) and 9 signals colocalized with DICE eQTLs in at least one cell type (Supplementary Figure 1D). Together, we found colocalization data to suggest putative target genes for 33 of our 56 signals (Figure 2A). Only 4 signals colocalized in all four QTL datasets and 19 of the signals with supporting colocalization data only colocalized in one dataset (Figure 2B).

**Figure 2:**
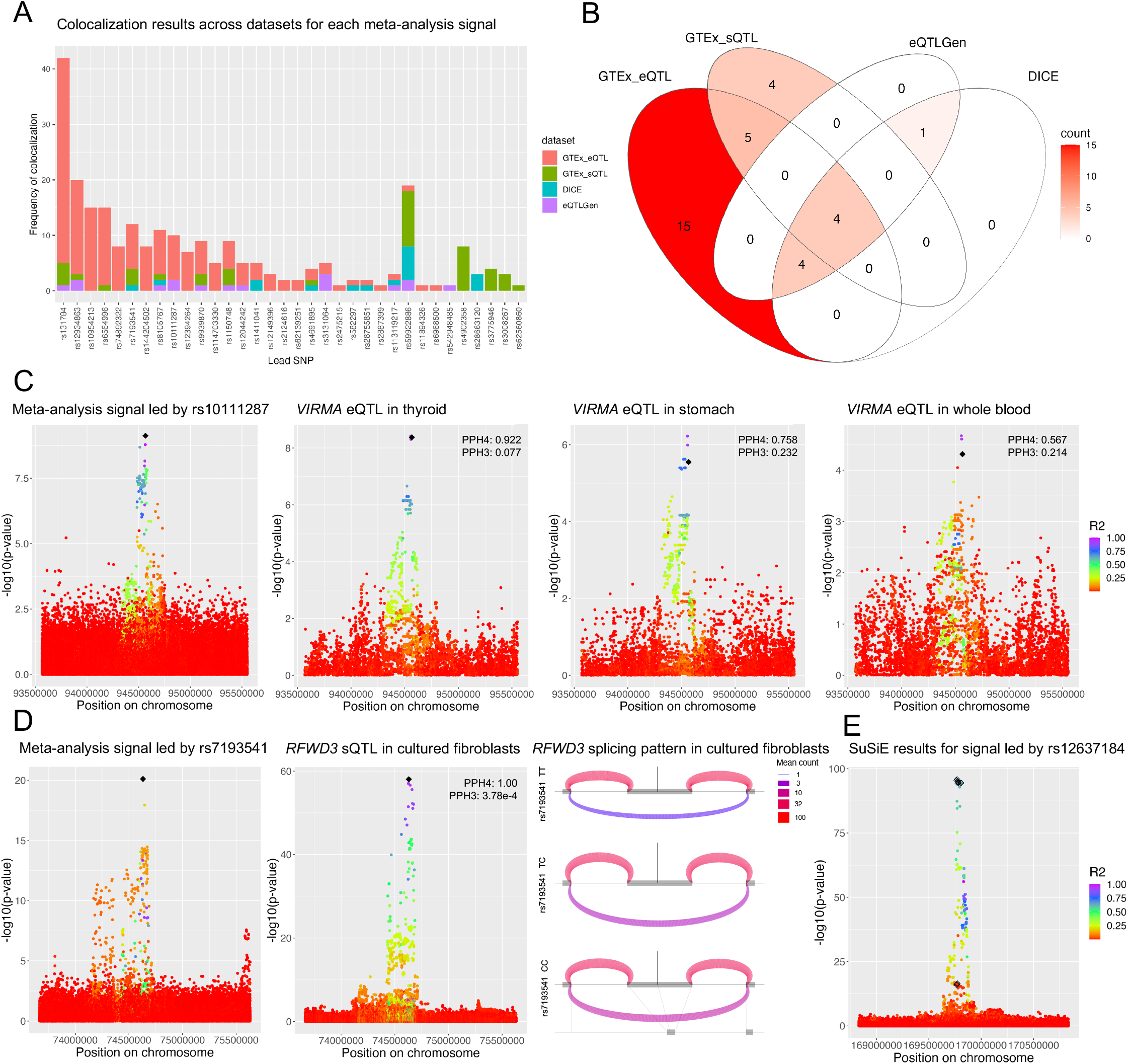
Fine-mapping analyses nominate putative causal variants and genes affecting telomere length. A. A barplot showing the number of colocalization events between a meta-analysis signal (labelled by the lead SNP) and a QTL for any gene in any cellular context across QTL datasets. All colocalization results for each signal are included in Supplementary Tables 3-6. B. Venn diagram showing which meta-analysis signals colocalized with any gene quantitative trait locus (QTL) in any cell type across datasets. We considered PPH4 > 0.7 to be colocalized for GTEx and eQTLGen. We considered PPH4 > 0.5 to be colocalized for DICE. C. Meta-analysis signal near rs10111287 colored by r^2^ with the sentinel SNP (black diamond) and *VIRMA* eQTLs in three GTEx tissues: thyroid, stomach, and whole blood. Colocalization results for each eQTL with the meta-analysis signal are indicated in the top right corner. PPH3 = posterior probability that the signals do not colocalize, PPH4 = posterior probability that the signals colocalize. Colocalization analysis between the eQTLs suggests there are shared causal SNPs: thyroid eQTL with stomach eQTL PPH3=0.090 PPH4=0.906, thyroid eQTL with whole blood eQTL PPH3=0.144 PPH4=0.745, stomach eQTL with whole blood eQTL PPH3=0.190 PPH4=0.655. D. Meta-analysis signal near rs7193541 colored by r^2^ with the sentinel SNP (black diamond) and *RFWD3* splicing QTL. Colocalization results for the QTL with the meta-analysis signal are in the top right corner. In the LeafCutter splicing cluster diagram grey boxes represent the *RFWD3* exons involved in the splicing cluster, the central exon is exon 14 and is located at chr16:74630780-74630957 (hg38). The curved lines represent the average number of reads spanning each exon-exon junction across individuals. Thinner, purple curves represent lower expressed exon-exon junctions and thicker, pink/red curves represent higher expressed exon-exon junctions. The plot is stratified by genotype of the lead SNP at the meta-analysis locus. The location of the lead SNP is depicted by the vertical grey line. The line at the bottom shows the linear base pair position of each exon and intron depicted in the plots. There were 167 TT individuals, 236 TC individuals, and 80 CC individuals included in this analysis. E. SuSiE 95% credible set results for the signal led by rs12637184. Black diamonds indicate SNPs predicted to be part of the 95% credible set. This signal had two credible sets, one comprised of SNPs at the top of the association peak and the second at approximately -log10(p-value) = 12. r^2^ is calculated with respect to the lead SNP at the signal.

To identify putative molecular mechanisms underlying each signal, we synthesized the available data to converge on a high likelihood candidate gene, where possible (Methods, Supplemental Note). 28 meta-analysis signals colocalized with QTLs for one gene but in multiple cellular contexts (Supplementary Tables 3-4). For example, the signal led by rs10111287 colocalized best with a *VIRMA* eQTL in thyroid (Figure 2C), but also significantly colocalized with *VIRMA* eQTLs in stomach and whole blood. Across genes, this signal only significantly colocalized with *VIRMA* eQTLs which made it straightforward to conclude this signal is likely linked to regulating *VIRMA* gene expression. Importantly, these results are not sufficient to make conclusions about the relevance of specific cellular contexts. Observed colocalization tends to correlate with the strength of the QTL, exemplified by the trend across the *VIRMA* eQTLs in thyroid (eQTL min p=3.79×10^-9^, PPH4=0.922), stomach (eQTL min p=5.94×10^-7^, PPH4=0.758), and whole blood (eQTL min p=2.13×10^-5^, PPH4=0.567). Variable power in eQTL data across tissues or cohorts is one reason that colocalization analysis is limited to suggesting candidate causal genes but not relevant cellular contexts (Urbut et al. 2019; Arvanitis et al. 2022).

#### Interpreting sQTL colocalization results

13 meta-analysis signals colocalized (PPH4 > 0.7) with a GTEx sQTL (Figure 2A-B), of which 4 also colocalized with an eQTL for the same gene (Supplementary Figure 1E). sQTLs are called based on exon read depth relative to other exons in the splicing cluster; a reduction in the expression levels of just one exon can result in the locus also being reported as an eQTL due to fewer total reads mapping to the gene. Therefore, it is possible for a signal regulating splicing to have colocalization results with an sQTL and an eQTL. This was the case for the signal led by rs7193541 (Figure 2D) which colocalized with an *RFWD3* sQTL in cultured fibroblasts (PPH4=1.000) and an *RFWD3* eQTL in skeletal muscle (Supplemental Note, PPH4=0.993).

This meta-analysis signal also colocalized (PPH4 > 0.7) with an *RFWD3* sQTL in two other GTEx tissues (EBV-transformed lymphocytes and brain cerebellar hemisphere) and an *RFWD3* eQTL in seven other GTEx tissues (adipose visceral omentum, adrenal gland, breast mammary tissue, liver, prostate, minor salivary gland, and transverse colon). We can be confident that splicing is the likely molecular mechanism if the splicing cluster is clear and supported by effects on expression over affected exons. A LeafCutter (Y. I. Li et al. 2018) plot of this splicing cluster demonstrated that individuals with more copies of the lead SNP at this locus increasingly excluded the fourteenth exon in *RFWD3* (Figure 2D). This was further supported by examining the RNA expression alignment which showed decreased expression of only the fourteenth exon in individuals with one or two copies of rs7193541 (Supplementary Figure 1F). This exon is excluded in observed RFWD3 protein isoforms (NP_001357465.1). These results lend strong support to the conclusion that this meta-analysis signal is driven by the association of telomere length with the regulation of *RFWD3* splicing and is it possible that this isoform may have distinct molecular effects on telomere length.

#### Interpreting conflicting colocalization analysis results

While colocalization analysis is an excellent tool for identifying potential causal genes for a meta-analysis signal, comparison across diverse cellular contexts and between datasets at times led to multiple putative causal genes. There were 6 meta-analysis signal-gene QTL colocalization pairs that were replicated between datasets (Supplementary Figure 1E). In 19 cases there was only colocalization evidence from one QTL dataset (Figure 2B) and in 14 cases there was conflicting colocalization results for a meta-analysis signal (Supplemental Note). For example, the signal led by rs59922886 colocalized strongly with a *CTC1* eQTL in GTEx sun exposed skin (PPH4 = 0.861). But in eQTLGen the same meta-analysis signal best colocalized with an *AURKB* eQTL (PPH4=0.919). Colocalization analysis from DICE further supported attribution to *CTC1* where the signal colocalized with a *CTC1* eQTL in M2 cells (PPH4=0.641). In this case, known biology allowed us to confidently attribute the signal to *CTC1* because CTC1 functions as part of the CST complex to regulate telomere length (Miyake et al. 2009; Surovtseva et al. 2009).

Recently there has been discussion about whether assigning genes to GWAS or meta-analysis signals should rely upon colocalization analysis as opposed to the proximal gene (Mostafavi et al. 2022). 20 of our 56 meta-analysis signals best colocalized with the proximal gene. We assigned a gene to each meta-analysis signal based on known biology of proximal genes (proximity-plus-knowledge) (Okamoto et al. 2023), colocalization analysis results, or the proximal gene where no other information was available. We discuss these situations and our rationale for putative causal gene assignment in the Supplemental Note.

#### Credible set analysis suggests that some loci consist of multiple independent causal variants which regulate the same gene in different contexts

To identify putative causal SNPs at each locus we applied fine-mapping using SuSiE (Zou et al. 2022) to estimate 95% credible sets. This analysis results in a set of SNPs estimated to contain a casual SNP with 95% confidence based on GWAS summary statistics and accounting for linkage disequilibrium estimates. We were able to identify 95% credible sets at 38 of 56 loci (Supplemental Table 7, Methods).

SuSiE identified two credible sets for the signal led by rs35510081 (Figure 2E). We did not observe any significant colocalization results for this locus. It is not unusual for a considerable proportion of GWAS signals to not colocalize with QTLs (Chun et al. 2017; Umans, Battle, and Gilad 2021; Connally et al. 2022; Mostafavi et al. 2022) and in such cases, prior knowledge and proximity to nearby genes is considered. In this case *TERC*, the RNA component of telomerase, is not the immediate proximal gene but is nearby (4.5 kb). Given the *a priori* information we have about *TERC* as a component of telomerase (Feng et al. 1995), we can be confident attributing this signal to *TERC*. In this and similar cases known biological information superseded the proximal gene or colocalization analysis results in assigning the peak (Supplemental Note).

16 of the 38 loci where credible set estimation was possible are predicted to have multiple causal SNPs. The number of predicted causal SNPs at each locus is consistent with conditional analysis on the pooled ancestry GWAS (Taub et al. 2022) (Supplementary Figure 1G). Many of these signals also have stronger association with telomere length and the detection of multiple causal SNPs is likely due to increased power. The exceptions to this trend are the *TERF1* locus, which is a telomere binding protein (Zhong et al. 1992), and the *DCLRE1B* (aka *APOLLO*) locus, which is important for telomere end processing (Lenain et al. 2006; van Overbeek and de Lange 2006; Wu et al. 2010). The association signals at these loci were not as strong (p=2.04×10^-12^ and p=3.26×10^-8^, respectively) yet are estimated to have 6 and 3 causal SNPs at the signals, respectively. We previously demonstrated that the multiple signals at the *OBFC1* (aka *STN1*) locus colocalize strongly with *OBFC1* eQTLs in distinct tissues (Taub et al. 2022). This is also true for *NAF1* (Supplementary Figure 1H). Both *NAF1* and *OBFC1* could be considered core telomere length regulation genes as they have direct mechanisms on biosynthesis and regulation of telomerase (Stanley et al. 2016; Miyake et al. 2009; Surovtseva et al. 2009) and their independent signals could reflect distinct regulatory mechanisms across cellular contexts. However, as discussed above, QTL detection can be influenced by technical factors, and from this work alone we are unable to eliminate the possibility that there may be undetected QTLs in these cellular contexts that would colocalize with one another. But the prevalence of multiple causal SNPs at many association signals reiterates the importance of these core genes in telomere length regulation across cellular contexts.

### Genes suggested by colocalization analysis highlight nucleotide synthesis and ubiquitination

We looked for GO biological process pathway enrichment using PANTHER (Mi et al. 2019; Thomas et al. 2022) and observed very strong enrichment of telomere regulation and DNA damage response pathways, as expected (Supplementary Table 8). We observed similar GO process enrichment using proximal genes and colocalization analysis-supported genes (Supplementary Figure 2). We also observed significant enrichment of nucleotide synthesis processes (e.g. cellular aromatic compound metabolic process, nucleic acid metabolic process). The importance of dNTP pools in regulating telomerase has been well documented (Hammond and Cech 1997; Gupta et al. 2013; Maicher et al. 2017; van Mourik et al. 2018) and one of the GWAS included in our meta-analysis also highlighted the importance of nucleotide metabolism in telomere length regulation (C. Li et al. 2020). Though we did not observe enrichment of any protein degradation biological processes, we attributed several of our meta-analysis signals to genes involved in proteasomal degradation including *UBE2D2*, *KBTBD6*, *PSMB4*, and *RFWD3*. *UBE2D2* is proximal to the rs56099285 signal and is an E2 ubiquitin conjugating enzyme (Saville et al. 2004). The signal near rs1411041 colocalized strongly with both *KBTBD6* and *KBTBD7*; these neighboring genes function as part of an E3-ubiquitin ligase complex (Genau et al. 2015). Additionally, we observed a signal near rs12044242 which we attributed to *PSMB4*, a non-catalytic component of the 20S proteasome (Nothwang et al. 1994), and a signal near rs7193541 which we and others attributed to *RFWD3*, an E3 ubiquitin ligase (Fu et al. 2010). Together this collection of genes highlights an unappreciated role of ubiqutination regulation in telomere length regulation dynamics.

### Meta-analysis signals are enriched for transcription factor binding sites of transcription factors with roles in telomere length regulation

Several transcription factors are known to regulate core telomere genes and disruption or creation of their transcription factor binding sites can result in dysregulation of telomerase and telomere length regulation (Huang et al. 2013). We examined whether the 95% credible set SNPs for our meta-analysis signals were enriched for transcription factor binding sites of any transcription factors with known consensus sequence using ENCODE ChIP-seq data (Figure 3A)(ENCODE Project Consortium 2012; Luo et al. 2020) or ReMap consensus sequences (Supplementary Figure 3A, Methods)(Hammal et al. 2022). We also analyzed the enrichment of the lead SNP alone at each meta-analysis signal (Supplementary Figure 3B-C). Many transcription factors involved in telomere length regulation had binding sites that were enriched in our meta-analysis using both analyses (Figure 3A, Supplementary Figure 3A, Supplementary Table 9). The transcription factor binding site enrichment calculated using ENCODE data was correlated with that of ReMap (95% credible set analysis R2 =0.336, lead SNP analysis R2 = 0.589)(Supplementary Figure 3D-E).

**Figure 3:**
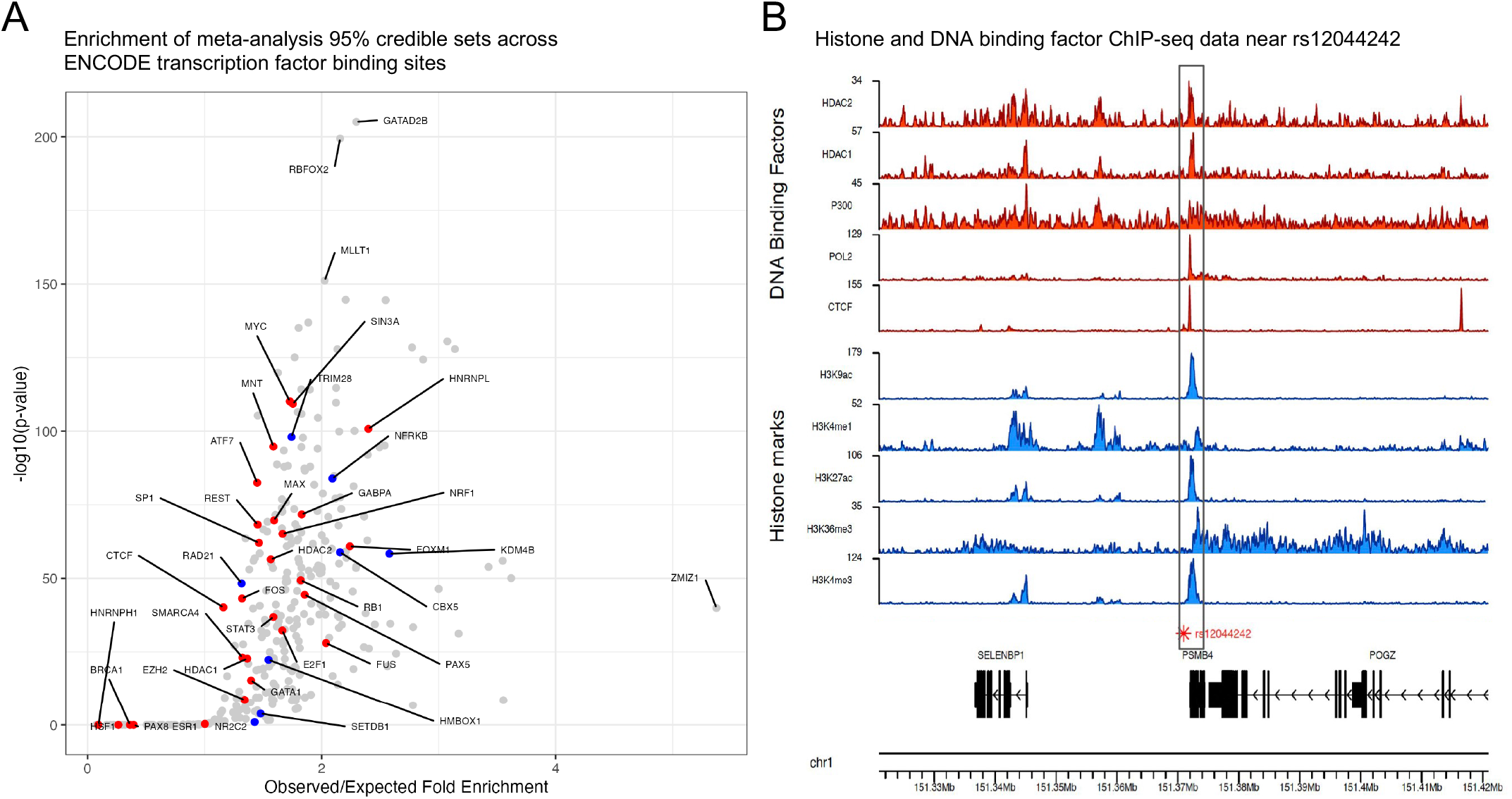
Meta-analysis signals are enriched for transcription factor binding sites of transcription factors with roles in telomere length regulation. A. The enrichment of 95% credible set SNPs across all transcription factors with ChIP-seq data available from ENCODE ChIP-seq data (Methods). Red points represent transcription factors with known roles in regulating telomere length regulation genes and blue points represent transcription factors with known roles in the alternative telomere lengthening (ALT) pathway. There were 18 transcription factors that fall at the (0,0) coordinate that are not plotted for the sake of clarity; one (XRCC3) had known roles in ALT. A complete list of transcription factors is provided in Supplementary Table 9. B. ChIP-seq data for the indicated DNA binding factor (red) or histone mark (blue) was generated by ENCODE and downloaded as bigwig files from the UCSC genome browser. The gene structure and genomic coordinates are depicted below the ChIP-seq data.

Previous work demonstrated that PAX5 increases *TERT* expression in B cells and fibroblasts (Bougel et al. 2010; Qin et al. 2021). We observed that there is a PAX5 transcription factor binding site overlapping the signal led by rs12044242, which we assigned to *PSMB4* (Supplemental Note). This SNP ablates a highly weighted cytosine in the consensus sequence and overlaps ChIP-seq peaks for activating histone marks (H3K4me3, H3K1me1, H3K27ac) and binding sites for transcriptional regulators (POL2, CTCF, HDAC1, HDAC2) (Figure 3B). Lead SNPs at signals we attributed to *OBFC1* and *TINF2,* both of which produce key telomere binding proteins, overlap binding sites for SOX2 and KLF4, respectively. In addition, one of our novel signals, which we attributed to the proximal gene *RRP12*, overlaps a MYC binding site. Furthermore, MYC is a well established regulator of *TERT* expression (Greider 1999). SOX2, KLF4, and MYC are pluripotency factors (Takahashi and Yamanaka 2006) and the presence of their binding sites at these telomere length association signals suggests regulatory roles for these genes in pluripotent cells. Our meta-analysis lead SNPs also overlapped transcription factor binding sites for FOXE1, GABPA, and HMBOX1 (Supplementary Table 10) which have all been reported to regulate expression of *TERT*, the protein component of telomerase (Bullock et al. 2016; Helbig et al. 2017; S. Zhou et al. 2017). Present literature on this topic has been focused on transcription factors regulating telomerase; these results demonstrate that these transcription factors may regulate other key telomere length regulation genes.

### *TCL1A* 95% credible set SNPs are more strongly associated with telomere length in older individuals

Because age accounts for a significant amount of telomere length variation (Demanelis et al. 2020), we ran a GWAS with an interaction term between age and genotype. Five signals had a genotype x age p-value that was below genome-wide significance (p-value < 5.39×10^-9^) and another 48 signals had genotype x age p-values that cleared suggestive thresholds (p-value < 5×10^-5^) (Supplementary Table 11). None of the genome-wide significant interaction signals were within 2 Mb of a meta-analysis signal, therefore we ran a GWAS stratified by age as an orthogonal approach (Supplementary Table 12). This analysis required individual-level data, therefore it was limited to the 109,122 individuals from TOPMed. We divided these individuals into three age groups ([0, 43], (43, 61], and (61, 98]) such that there were a similar number of individuals in all three groups. Expanding the analysis to more granular age groups was not possible with this sample size without singularity issues in the GWAS analysis. Although the ratio of males to females was similar between groups (Supplementary Figure 4A), the distribution of ancestries varied such that the proportion of European individuals increased over age (Supplementary Figure 4B). We filtered candidate regions to identify loci with similar minor allele counts between groups, but with non-overlapping effect size estimate confidence intervals. We also required that the locus have a minimum SNP x age interaction p-value < 5×10^-5^ and that the locus have a genome-wide significant association signal (p < 5×10^-8^) in the meta-analysis (Methods). The rs2296312 locus was the single locus that met the filtering pipeline criteria with a SNP x age interaction p-value = 2.599×10^-6^ (Figure 4A). The effect size estimate increased over age (Figure 4B) and this trend was independent of ancestry as the effect estimate for rs2296312 was similar between all examined ancestries (Figure 4C). The association signal increased in significance over age, mirroring the effect size estimate trend (Figure 4D-F). In the meta-analysis, rs2296312 was part of a peak that colocalized best with a *TCL1A* eQTL from GTEx whole blood (PPH4 = 0.714). SuSiE credible set analysis identified 14 SNPs in the credible set for this peak all of which have a similar trend in their effect estimates over age. Together these data demonstrate that putative causal SNPs regulating *TCL1A* expression are associated with age and telomere length. TCL1A activates the AKT signaling pathway increasing cellular proliferation (Pekarsky et al. 2000) and *TCL1A* expression was previously reported to decrease in whole blood as age increases (Demanelis et al. 2020). Furthermore, rs2296312 has been reported to act through *TCL1A* to be protective against loss of the Y chromosome and clonal hemoatopoesis (W. Zhou et al. 2016; Weinstock et al. 2023). Our data are concordant with previous findings and suggest that these protective phenomena reduce proliferation, leading to longer telomere length.

**Figure 4:**
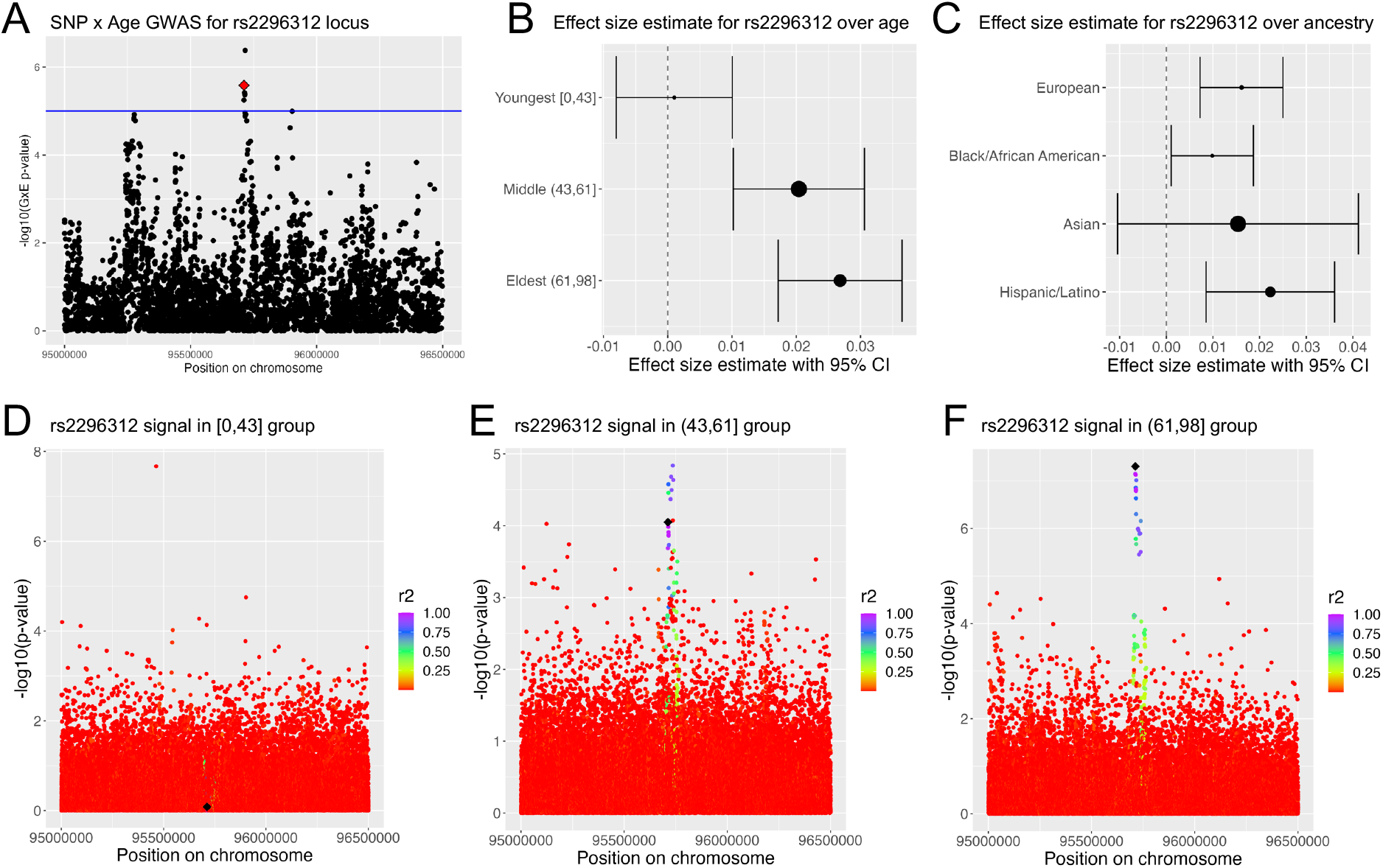
***TCL1A* 95% credible set SNPs are more strongly associated with telomere length in older individuals.** A. Manhattan plot for the region around rs2296312 (red star) using summary statistics from a GWAS that included a covariate for age and genotype interaction. The log10(p-value) for the interaction covariate is plotted on the y-axis. B. Forest plot indicating the effect size estimate for rs2296312 across age groups from the age-stratified GWAS. The tested allele, C, was the minor allele. [0,43] minor allele count = 15,922; (43,61] minor allele count = 16,315; (61,98] minor allele count = 13,547. C. Forest plot indicating the effect size estimate for rs2296312 across ancestry groups from ancestry-stratified GWAS (Taub et al. 2022). European minor allele count = 16,443; Black/ African American minor allele count = 19,963; Asian minor allele count = 5,683; Hispanic/Latino minor allele count = 18,019. D-F. Manhattan plots for the rs2296312 (black diamond) locus in age-stratified GWAS. Color indicates linkage disequilibrium (r2) calculated with respect to rs2296312.

### Blood and immune cells are a key cell type for telomere length

To understand the biology of our associated loci and to support validation of our findings, we first had to determine the most relevant cellular context to examine telomere length associated signals. Telomere length was estimated from blood leukocytes in all samples, however, telomere length regulation is relevant in many different cell types, to differing extents (Armanios 2013). In relevant cellular contexts, causal SNPs are expected to be in genomic regions with active chromatin states. We tested for enrichment of the meta-analysis lead SNPs across Roadmap Epigenomics samples (Supplementary Table 13) and the 25 state chromHMM model (Figure 5A) (Roadmap Epigenomics Consortium et al. 2015). The strongest enrichment of several active chromatin states was observed in blood and T-cell samples. Because the chromHMM model is a predicted state, we also examined whether there was enrichment when looking at the primary data for specific chromatin marks. Consistent with the chromHMM model results, we saw that the strongest enrichment of lead SNPs in H3K4me1 and H3K27ac peaks was in blood and T-cell samples (Supplementary Figure 5).

**Figure 5:**
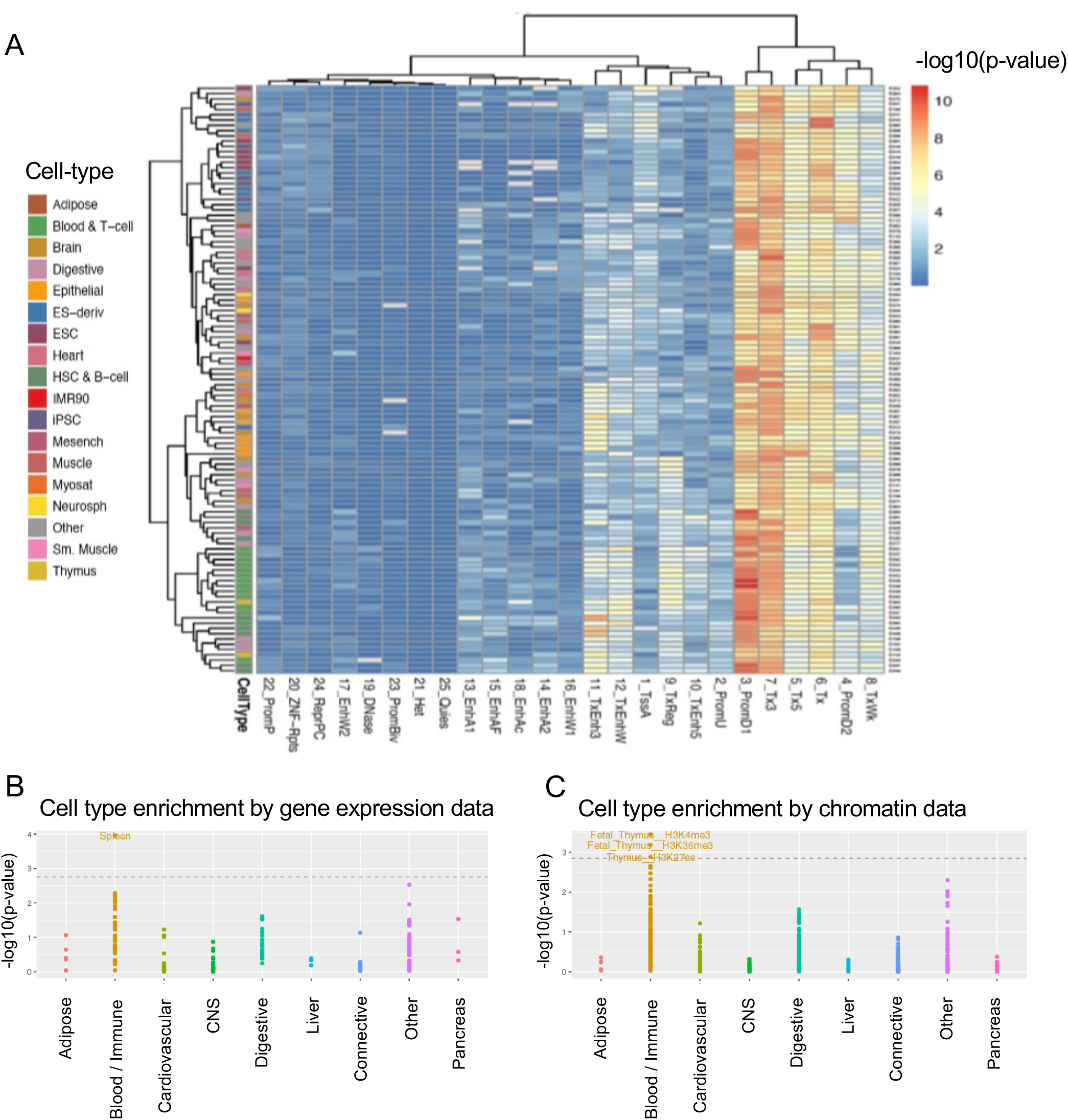
Blood and immune cells are a key cell type for telomere length. A. Hierarchical clustering of the enrichment of meta-analysis lead SNPs in predicted active states using the Roadmap Epigenomics 25 state chromHMM model. B-C. Stratified LDSC was conducted on 130,246 meta-analyzed European individuals in our dataset (Li et al. 2020; Taub et al. 2022) using the 1000 Genomes European linkage disequilibrium reference panel.

As an orthogonal approach we ran stratified linkage disequilibrium score regression (S-LDSC) on the meta-analyzed European individuals in our study (Methods). S-LDSC uses the meta-analysis summary statistics to examine whether, given linkage disequilibrium, a category of SNPs has increased association with telomere length compared to SNPs not in that category. In this case, we used categories based on previously reported cell type specific annotations based on gene expression or chromatin marks (Finucane et al. 2015). Using both gene expression and chromatin marks we observed that the blood/immune cell category was the only category that was significantly enriched (Figure 5B-C). Together with the Roadmap Epigenomics enrichment analysis, these data suggest that blood and immune cells are the most relevant cell type for genetic regulation of leukocyte telomere length.

### Overexpression of *POP5* and *KBTBD6* increases telomere length in HeLa-FRT cells

We began our validation experiments by screening candidate genes for an effect on telomere length. It has been well documented that shRNAs with loss of function effects often become epigenetically silenced over time in cell culture (Goff 2021). Therefore, we identified candidate genes where the lead SNP was predicted to increase gene expression. Of those we chose five genes that had one known protein coding sequence isoform, had strong colocalization analysis results, and had some known biology: *OBFC1*, *PSMB4*, *CBX1*, *KBTBD6*, and *POP5* (Methods). To generate constitutive overexpression cell lines we used the Flp-in system (Thermo Fisher Scientific) to incorporate the FLAG-tagged gene of interest under the control of a CMV promoter into HeLa-FRT cells (Methods). HeLa cells are not derived from blood or immune cells but are highly tractable for this screening stage of the validation experiments. Three independent transfection clones were passaged and the effect of gene overexpression on telomere length was observed by Southern blot.

The lead SNPs for each meta-analysis signal that we attributed to these genes was estimated to have a positive effect on telomere length in our meta-analysis (Supplementary Table 2), therefore we predicted that overexpression of these genes should increase telomere length. As a control we also overexpressed *GFP,* which had no effect on telomere length, as expected (Figure 6). Overexpression of *OBFC1* or *PSMB4* also had no effect on telomere length (Supplementary Figure 6A). Overexpression of *CBX1* slightly increased telomere length (Supplementary Figure 6A) while overexpression of *KBTBD6* or *POP5* showed a clear telomere length increase over increased cell division, concordant with the expectation from our meta-analysis (Figure 6). The median, minimum, and maximum telomere lengths were estimated for each lane in the Southern blots using ImageQuant TL (Methods, Supplementary Figure 7). Protein expression was assayed by western blot analysis. Western blot comparison of early population doubling timepoints to late population doubling timepoints showed that *POP5* overexpression was maintained through the duration of the experiment while *KBTBD6* overexpression was suppressed in clones 6 and 7 (Supplementary Figure 6B). This likely accounts for the plateau in telomere lengthening in *KBTBD6* overexpression clone 7 (Figure 6A- B).

**Figure 6:**
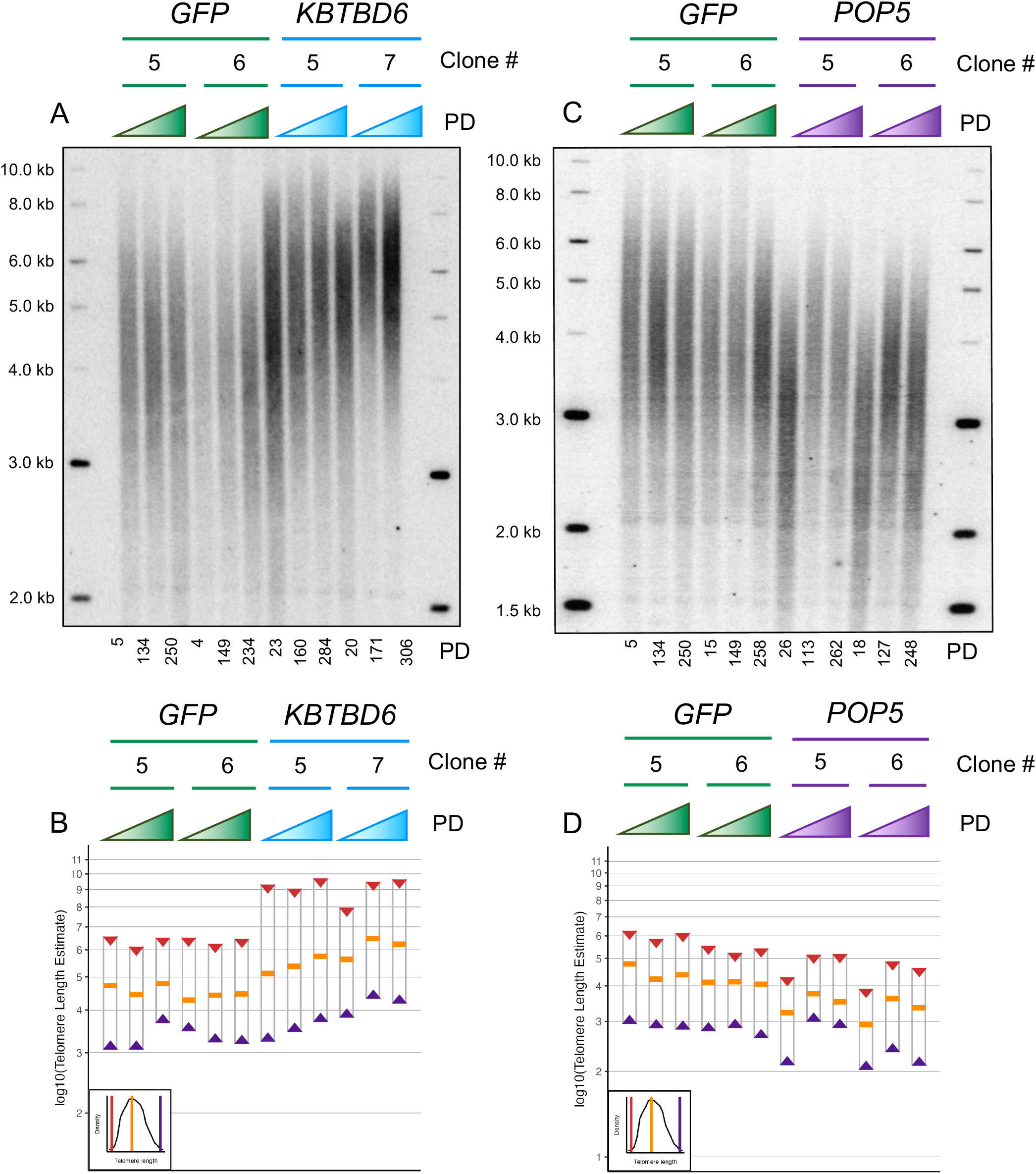
Overexpression of *POP5* or *KBTBD6* increases telomere length in HeLa-FRT cells. *KBTBD6*, *POP5*, or *GFP* was constitutively overexpressed from the CMV promoter in HeLa-FRT cells using the FLP-in system. A,C. Telomere Southern blots showing the bulk telomere length from a population of cells following an approximate normal distribution. Molecular weight standards were run alongside the samples and their size is indicated in kilobases (kb). Three time points are shown for each clone and the estimated number of population doublings (PD) for each timepoint are indicated below the Southern. Each clone has the opportunity to form a distinct starting telomere length distribution which is why the first timepoint for some clones appear to have distinct telomere length distributions, for example the starting timepoint for the *POP5* clones compared to the *GFP* clones. All transfection experiments began from the same population of HeLa-FRT cells. B,D. The Southern blot densitometry was analyzed using ImageQuant TL to generate line plots of the pixel density. The software estimated the median telomere length (orange bar) as the pixels with greatest density and estimated a molecular weight for that position taking into account the molecular weight standards on both sides of the gel. The ImageQuant TL line plots (Supplementary Figure 7) were used to estimate the minimum (purple triangle) and maximum (red triangle) telomere lengths in the bulk telomere band. A simulated diagram in the bottom left of the plot representing the ImageQuant TL plots is provided as a guide for the source of these values. The y-axis is plotted on a log10 scale to better estimate how linear DNA moves through an agarose gel at rate inversely proportional to its length.

### CRISPR removal of *KBTBD6* and *POP5* regulatory regions reduced expression of each gene

We next sought to examine whether high likelihood causal elements in the respective meta-analysis signals affect the expression of these genes. SuSiE was unable to predict a 95% credible set analysis for the *POP5* locus, likely because the association signal is below genome-wide significance in the summary statistics used for fine-mapping (Methods). We utilized a second credible set estimation algorithm, CAVIAR (Hormozdiari et al. 2014), with a single assumed causal SNP, however, the 95% credible set included 3,041 SNPs and did not reduce the position range of the region (Supplementary Figure 8A). In the absence of useful 95% credible set estimation, we considered the genome region spanning the lead SNP and SNPs with r^2^ > 0.9 and p-value < 1×10^-6^ (Supplementary Figure 8B). To prioritize a subset of this 124 kb region, we intersected these top SNPs with ATAC-seq, Hi-C, and chromatin ChIP-seq data from blood samples, but were unable to form a consensus (not shown). We removed the 124 kb region upstream of *POP5* using CRISPR/Cas9 in K562 cells (Supplementary Figure 8C) and identified 24 clones where the region had been successfully deleted at one allele, generating heterozygous deletions (Methods). qPCR analysis (primer sequences in Supplementary Table 14) of these clones showed significantly reduced *POP5* expression compared to controls (p=0.047) demonstrating that this region contains critical SNPs for regulating *POP5* expression in blood cells (Figure 7A).

**Figure 7:**
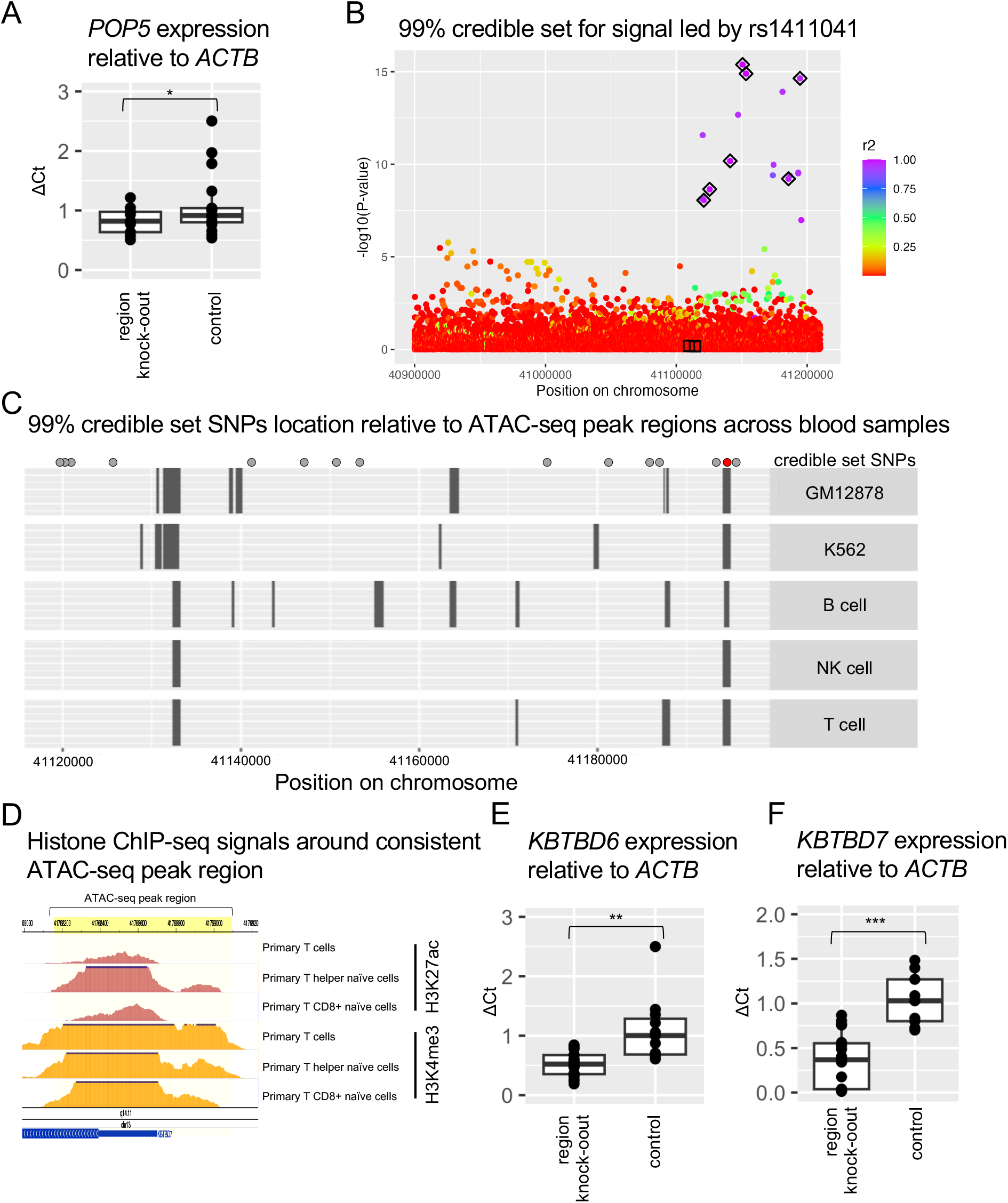
CRISPR removal of *KBTBD6* and *POP5* regulatory regions reduced expression of each gene. A. qPCR estimates of *POP5* expression were normalized to *ACTB* using the Pfaffl method (Methods). A one-sided *t*-test calculated a p-value = 0.047. B. 99% SuSiE credible set colored by r^2^ with the lead SNP. Black diamonds indicate SNPs in the predicted credible set. C. ATAC-seq peak regions are represented as boxes for each blood related sample. Points above the plot area represent SNPs in the 99% credible set predicted by SuSiE or CAVIAR. The 95% credible set from either SuSiE or CAVIAR did not overlap any regions where ATAC-seq peaks were shared across blood cell types and cell lines. The red SNP is rs9525462. NK cell = natural killer cell. Samples were downloaded from ENCODE (ENCODE Project Consortium 2012; Luo et al. 2020) (identifiers: ENCFF058UYY, ENCFF333TAT, ENCFF421XIL, ENCFF470YYO, ENCFF558BLC, ENCFF748UZH, ENCFF751CLW, ENCFF788BUI, ENCFF867TMP) or from ATACdb (Wang et al. 2021)(sample codes: Sample_1195, Sample_1194, Sample_1175, Sample_1171, Sample_1020, Sample_1021, Sample_1209, Sample_1208). D. Roadmap chromatin ChIP-seq for hg19 chr13:41768158-41769095 (yellow highlighted region). Samples included were E044, E039, and E047. E-F. qPCR estimates of gene expression were normalized to *ACTB* using the Pfaffl method (Methods). A one-sided *t*-test calculated a p-value = 0.003037 for *KBTBD6* and p-value = 2.093×10^-5^ for *KBTBD7*. * p-value < 0.05 ** p-value < 0.01 *** p-value < 0.001.

KBTBD6 functions as a component of an E3 ubiquitin ligase complex along with CUL3 and KBTBD7 (Genau et al. 2015). *KBTBD7* is a neighboring gene and we observed colocalization with the signal led by rs1411041 with both *KBTBD6* and *KBTBD7* eQTLs in GTEx (Supplementary Table 3). We were interested in determining whether CRISPR editing of high likelihood SNPs in this meta-analysis signal would affect the expression of *KBTBD6*, *KBTBD7*, or both. We intersected the position of the 99% credible set SNPs (Figure 7B) with ATAC-seq peaks in blood samples (Figure 7C). Only one SNP, rs9525462, was located in a region where the ATAC-seq peaks were shared across blood samples. rs9525462 was predicted to be in the 99% credible set by both SuSiE and a second credible set analysis software, CAVIAR. This region overlaps promoter and enhancer chromatin marks (H3K27ac and H3K4me3, respectively) in Roadmap Epigenomics blood samples (Figure 7D), further supporting that this region is in an active state in blood samples. We used CRISPR/Cas9 to remove the 938 bp ATAC-seq peak region in K562 cells (Supplementary Figure 8D) and identified 31 clones where this region had been successfully removed at least at one allele, generating heterozygous deletions (Methods). Clones with the ATAC-seq peak region knocked-out had significantly decreased *KBTBD6* (p=0.003037) and *KBTBD7* (p=2.093e-05) expression relative to controls, demonstrating that this region is critical in regulating the expression of both genes. Together these data demonstrate that our meta-analysis signals are driven by *POP5* and *KBTBD6/KBTBD7*, and we identify them as novel telomere length regulation genes.

## Discussion

Our results demonstrate the utility of telomere length GWAS in the identification of novel telomere length regulatory mechanisms. Our fine-mapping of telomere length associated loci and discussion of relevant cell types in which to validate these signals is a useful platform for further experimental validation. We determined that blood and immune cells are the most relevant cellular context to examine leukocyte telomere length association signals based on chromatin accessibility and S-LDSC. Telomere length was estimated from blood leukocytes in all samples; it is possible that this boosted the strength of blood and immune cell enrichment in our analyses. However, telomere length regulation is relevant in many different cell types, to differing extents (Armanios 2013). We propose that blood and immune cells are the most relevant cell type for leukocyte telomere length GWAS validation experiments, but that these genes contribute to telomere length regulation across cellular contexts. This idea is further supported by our observation that independent association signals at the *OBFC1* (Taub et al. 2022) and *NAF1* loci colocalize with eQTLs for their respective genes in different cellular contexts.

While prior telomere length GWAS (C. Li et al. 2020; Codd et al. 2021) have used colocalization to support putative causal genes for their association signals, we extended this work to include multiple QTL datasets across tissues and to include splicing in addition to expression QTLs. This made it possible to uncover splicing mechanisms that may be associated with telomere length, as we saw with *RFWD3*, and increased the confidence of our putative causal gene assignment.

Experimental validation of putative causal genes identified novel genes involved in telomere length regulation. POP5 is a subunit of the Ribonuclease P/MRP complex (van Eenennaam et al. 2001). Previous work in *S. cerevisiae* demonstrated a role for specific components of the homologous complex in telomerase holoenzyme complex regulation (Laterreur et al. 2018). In addition, POP1, another subunit of the Ribonuclease P/MRP complex, was recently shown to interact with human telomerase RNA (Zhu et al. 2023). Together, these results suggest that the role of the POP proteins also play a role in human telomerase regulation. KBTBD6 and KBTBD7 are members of an E3 ubiquitin ligase complex (Genau et al. 2015). CRISPR/Cas9 deletion of the high-likelihood causal region affected expression of both genes, but overexpression of *KBTBD6* alone affected telomere length. Our results suggest that increased expression of the KBTBD6-KBTBD7-Cul3 complex or altered complex stoichiometry affect telomere length.

In addition to the *KBTBD6/KBTBD7* signal, we observed association signals that we attribute to *RFWD3*, another E3 ubiquitin ligase, *PSMB4*, a component of the core proteasome, and *UBE2D2*, an E2 ubiquitin conjugating enzyme. ATM and ATR are kinases that contribute to the DNA damage response and telomere length regulation, though phosphorylation targets with strong effects on telomere length regulation have remained elusive (S. S. Lee et al. 2015; Tong et al. 2015; de Lange 2018; Keener, Connelly, and Greider 2019). Prior proteome analysis demonstrated that ATM/ATR regulate the ubiquitin-proteasome pathway in response to DNA damage and validated RFWD3 as an ATM/ATR substrate (Mu et al. 2007; Fu et al. 2010). Our results underscore the importance of ubiquitination in telomere length regulation; future work examining whether ATM/ATR substrates regulating the ubiquitination-proteasome pathway affect telomere length may identify ATM/ATR substrates with important roles in telomere length regulation. Furthermore, identification of the ubiquitination targets by these E3 ubiquitin ligases may reveal novel telomere length regulation mechanisms. Together, this work demonstrates the potential contribution of telomere length GWAS to understanding mechanisms underlying telomere length regulation. Future work extending the findings reported here and validating additional loci will increase our understanding of both the genetics and molecular mechanisms underlying telomere length regulation.

## Acknowledgements

We thank Chen Li, Claudia Langenberg, Veryan Codd, Dayana Delgado, Brandon Pierce, and Rajkumar Dorajoo, in addition to all the individuals who were sampled, for providing summary statistics from their telomere length GWAS that were included in this meta-analysis. The whole-genome sequencing for the Trans-Omics in Precision Medicine (TOPMed) program was supported by the National Heart, Lung, and Blood Institute (NHLBI). Specific funding sources for each study and genomic center are given in the Supplemental Acknowledgements. px458 was a gift from Andrew Holland’s lab. We acknowledge the ENCODE Consortium and the following ENCODE production laboratories: Michael Snyder and J. Michael Cherry. We would also like to thank Margaret Strong, Emily DeBoy, the JHU Synthesis & Sequencing Facility, and the JHU Ross Flow Cytometry Core for their technical assistance, Andrew Holland and Emmanouil Tampakakis for helpful discussion about CRISPR/Cas9 editing experiments, and the Mathias, Greider, and Battle labs for helpful discussion throughout the course of this work.

## Author Contributions

R.Keener, C.W.G, R.A.M, and A.B. conceived of and led the study. R.Keener, S.B.C., C.J.C., M.A.T., J.S.W., L.R.Y., L.M.R., A.P.R., C.W.G., R.A.M., and A.B. drafted the manuscript. R.Keener, S.B.C., C.J.C, M.A.T, M.P.C., J.S.W., B.N., B.J.S., M.A., C.W.G, R.A.M., and A.B. contributed substantive analytical guidance. R.Keener, S.B.C., C.J.C, M.A.T, M.P.C., J.S.W., B.N., B.J.S., M.A., C.W.G, R.A.M., and A.B. performed and led the analysis. R.Keener, M.A.T., M.P.C., J.S.W., S.A., P.L.A., L.B., L.C.B., J.B., E.R.B., J.A.B., B.E.C., J.C.C., Y.C., L.A.C., B.C., B.I.F., M.T.G., S.R.H., L.H., M.R.I., C.R.I., J.M.J., E.E.K., C.K., R.L.M., S.N., N.P., P.A.P., J.I.R., K.D.T., M.J.T., B.W., L.R.Y., I.V.Y., C.A., D.K.A., A.E.A.K., K.C.B., J.C.B., T.W.B., E.B., E.G.B., A.P.C., Z.C., Y.I.C., D.D., M.dA., P.T.E., M.F., B.D.G., F.D.G., J.H., T.I., S.Kaab, S.L.R.K., S.Kelly, B.A.K., R.Kumar, R.J.F.L., F.D.M., S.T.M., D.A.M., B.D.M., C.G.M., K.E.N., N.D.P., J.M.P., B.A.R., S.R., S.S.R., D.R., I.R., D.S., F.S., M.B.S., E.K.S., M.F.S., N.L.S., A.V.S., H.K.T., R.S.V., S.T.W., L.K.W., Y.Z., E.Z., L.M.R., A.P.R., M.A., R.A.M., and A.B. were involved in the guidance, collection, and analysis of one or more of the studies that contributed data to this article. All of the authors read and approved the final draft.

## Methods

### Studies and telomere length estimation

We incorporated four telomere length GWAS with non-overlapping cohorts. (Delgado et al. 2018) had 5,075 samples from Bangladeshi individuals and telomere length was estimated using qPCR or Luminex-based assay. (Dorajoo et al. 2019) had 23,096 samples from Singaporean Chinese individuals and telomere length was estimated using qPCR. (C. Li et al. 2020) had 78,592 samples from European individuals and telomere length was estimated using qPCR. (Taub et al. 2022) had 51,654 individuals of European ancestry, 5,683 individuals of Asian ancestry, 29,260 individuals of African ancestry, and 18,019 individuals of Hispanic/Latino ethnicity. In this study telomere length was estimated bioinformatically from whole genome sequencing data (Taliun et al. 2021) using TelSeq (Ding et al. 2014).

### Meta-analysis

One concern with a meta-analysis approach was whether it is reasonable to compare summary statistics from GWAS where telomere length was estimated using different methods. Previous work determined that each method produces telomere length estimates that are highly correlated with Southern blot analysis (Aviv et al. 2011; Pierce et al. 2016; Taub et al. 2022) and in each study telomere length estimates were standardized prior to running the GWAS. We used GWAMA (Mägi and Morris 2010) to conduct a random effect meta-analysis that represents a total of 211,379 individuals. Taub et al. stratified individuals from the Trans-Omics for Precision Medicine (TOPMed) program cohorts by ancestry group where individuals were broadly categorized as European, African, Asian, or Hispanic/Latino using HARE and we maintain language used from that study here for clarity. That study also defined an “Other” group which was not included in our analysis. We provide a list of TOPMed cohorts whose data are represented in the meta-analysis and the broad ancestral groups individuals were categorized as (Supplementary Table 1). A detailed enumeration of individuals over ancestry by TOPMed cohort was previously published in Supplementary Table 1 of Taub et al. SNP positions were converted to hg38 using LiftOver (Hinrichs et al. 2006) prior to meta-analysis. The Delgado et al. summary statistics were harmonized to the forward strand and palindromic SNPs were removed from this dataset. Loci were considered novel if there were no other reported sentinels within 1 Mb of the lead SNP in the signal.

Lead SNPs were identified by minimum p-value within a 2 Mb window. We examined all loci with at least one variant that was genome-wide significant (p-value < 5×10^-8^) and had a minor allele frequency > 0.0001. This excluded loci where the lead SNPs were rs903494390, rs976923370, rs990671169, rs982808930, rs992178597, rs961617801, and rs1324702094. The signal led by rs3131064 is near the *HLA* locus and due to the extensive linkage disequilibrium in this region, we expanded the width of this signal to 4.2 Mb.

### Colocalization analysis

All colocalization analysis was conducted using the coloc package (Giambartolomei et al. 2014) using the coloc.abf() command with the prior probability that the SNP is shared between the two traits (p12) set to 1e-6 and that there was at least 1,000 shared variants between the two datasets. For GTEx_v8 (GTEx Consortium 2020) colocalization we evaluated all genes for which the lead SNP was a significant QTL in any of the 49 GTEx_v8 tissues. For colocalization with eQTLGen cis-eQTLs (version available 2019-12-11)(Võsa et al. 2021) and DICE cis-eQTLs (version available 2019-06-07)(Schmiedel et al. 2018) we evaluated all genes within a 2 Mb window centered on the lead SNP and the meta-analysis summary statistics were lifted down to hg19 using LiftOver (Hinrichs et al. 2006) to compare SNPs based on chromosome and position. The X-chromsome signals could not be evaluated for colocalization with eQTLGen data as that dataset is limited to autosomes. Colocalization was conducted using minor allele frequency, p-value, and the number of samples for eQTLGen. Minor allele frequency was estimated from TOPMed pooled across ancestries. For all other colocalization analyses effect size estimates and their standard errors were used. We report the posterior probability that there are two signals but they do not share a causal signal (PPH3) and the posterior probability that there are two signals and they do share a causal signal (PPH4) within the text, figures, and figure legends. Posterior probabilities for the cases that there is no signal in one or either of the datasets (PPH0, PPH1, and PPH2) are reported in the appropriate Supplementary Tables (3-6). We considered cases where PPH4 > 0.7 to be colocalized except for colocalization analysis with DICE cis-eQTLs where we reduced this threshold to PPH4 > 0.5 to account for the reduced power in the dataset. For Manhattan plots colored by linkage disequilibrium, r^2^ was calculated using a trans-ancestry group of all TOPMed individuals included in the meta-analysis.

### Visualizing sQTLs

RNA alignment information for each individual was extracted using SAMtools (version 1.16) (Danecek et al. 2021) in the GTEx_v8 cultured fibroblast samples on AnVIL (Schatz et al. 2022). We extracted genotype information from GTEx_v8 for the corresponding individuals and plotted the average alignment depth at each base position (hg38) stratified by genotype using Matplotlib (Hunter 2007). Visualization of LeafCutter (Y. I. Li et al. 2018) splicing clusters was produced using LeafCutter exon-exon junction quantifications generated by GTEx_v8 (GTEx Consortium 2020).

### Variant fine-mapping

Due to the trans-ancestry nature of our meta-analysis we used individual-level data from TOPMed individuals spanning all four ancestries represented in our meta-analysis (European, Asian, African, and Hispanic/Latino) as our linkage disequilibrium reference. Despite the fact that TOPMed individuals represent the largest group in the meta-analysis, the mismatch between the linkage disequilibrium reference and meta-analysis summary statistics was problematic for SuSiE (susieR_0.12.16) (G. Wang et al. 2020; Zou et al. 2022). Therefore, we used summary statistics from the pooled TOPMed GWAS ((Taub et al. 2022) to estimate credible sets for all meta-analysis signals (Supplementary Table 7) and generated a genotype correlation matrix using a random subset, preserving the proportion of ancestries, of 15,000 TOPMed individuals to manage SNP density. We did not use a minor allele frequency threshold for SNP inclusion. At 2 loci the signal was over 1 Mb wide and calculating the genetic correlation matrix exceeded the ability of computational resources on the premises. At 16 loci there was not sufficient signal in the TOPMed GWAS to predict a credible set. CAVIAR (Hormozdiari et al. 2014) requires specification of the assumed number of causal signals whereas SuSiE jointly models the likelihood of varying numbers of causal signals and converges on the highest likelihood case. Due to this assumption and the computational burden of running CAVIAR, we only ran CAVIAR on the *POP5* and *KBTBD6*/*KBTBD7* loci.

For the signal led by rs1411041, which we attributed to *KBTBD6* and targeted for CRISPR/Cas9 editing, we further fine-mapped the locus by intersecting the credible set SNPs with ATAC-seq peaks and with ChIP-seq data from Roadmap Epigenomics. ATAC-seq data were downloaded from ENCODE (ENCODE Project Consortium 2012; Luo et al. 2020)(identifiers: ENCFF058UYY, ENCFF333TAT, ENCFF421XIL, ENCFF470YYO, ENCFF558BLC, ENCFF748UZH, ENCFF751CLW, ENCFF788BUI, and ENCFF867TMP) or from ATACdb (F. Wang et al. 2021) (Sample_1195, Sample_1194, Sample_1175, Sample_1171, Sample_1020, Sample_1021, Sample_1209, and Sample_1208). BEDTools (Quinlan and Hall 2010) was used to identify intersecting regions. Roadmap Epigenomic ChIP-seq data was visualized using the WashU Epigenome browser (D. Li et al. 2019).

### GO enrichment analysis

All gene ontology (GO) enrichment analysis was conducted using PANTHER (Thomas et al. 2022; Mi et al. 2019) overrepresentation test with the GO Ontology database (released on 2022- 07-01) with the all *Homo sapiens* gene set list as the reference list. PANTHER GO biological process complete terms were tested for enrichment using a Fisher’s exact test with false discovery rate correction. Proximal genes were assigned as the gene with minimal distance to the gene body in the UCSC genome browser (Kent et al. 2002).

### Transcription factor binding site analyses

To assess the enrichment of 95% credible set SNPs with transcription factor and chromatin regulator DNA binding sites, we downloaded the ENCODE regulation track transcription factor binding site cluster ChIP-seq index file to report data for 330 DNA binding proteins spanning 129 cell types (ENCODE Project Consortium et al. 2020). The intersection of variants with transcription factor binding sites was performed by BEDTools v2.29.2 (Quinlan and Hall 2010). We computed the enrichment of 95% credible set SNPs in transcription factor binding sites using a GREGOR Perl based pipeline (Schmidt et al. 2015). Briefly, this pipeline sums independent binomial random variables for the number of index SNPs falling in a single feature and calculates the enrichment p-value using a saddlepoint approximation method. The SNPs are considered to have a positional overlap if the input SNP, or variants in high linkage disequilibrium with the input SNP (r^2^ > 0.7, linkage disequilibrium window size = 1 Mb), fall within the regulatory features or overlap by ≥ 1 bp. The pairwise linkage disequilibrium (r^2^) was computed using the 1000 Genomes European reference panel (1000 Genomes Project Consortium et al. 2015). Transcription factor binding site fold enrichment is measured as the fraction of index SNPs (or SNPs in linkage disequilibrium) overlapping the feature (as observed) over the mean number of overlaps with the control set of SNPs (as expected). Control SNPs are matched based on the number of variants in linkage disequilibrium, minor allele frequency, and distance to the nearest gene of the index SNPs. We also performed the enrichment analysis of 95% credible set SNPs with 1,210 DNA-associated factors spanning across 737 cell-tissue types using the peak bed files downloaded from the ReMap 2022 database (Hammal et al. 2022) using the same pipeline. In addition, we performed both the ENCODE and ReMap enrichment analyses using only the lead SNP at each signal (Supplementary Figure 3B-C). In addition to the enrichment analysis, we identified transcription factor binding sites overlapping the lead SNP for each meta-analysis association signal by searching the rsID on the UCSC genome browser (Kent et al. 2002; Hinrichs et al. 2006) and identified overlapping binding sites using the JASPAR 2022 track with default settings (Castro-Mondragon et al. 2022). We identified transcription factors with known roles in telomere length regulation by searching PubMed. Publication references supporting known roles for these transcription factors are indicated in Supplementary Table 9.

### Telomere length GWAS with an age x genotype interaction term

We repeated the pooled analysis from Taub et al. 2022 using all 109,122 TOPMed individuals with telomere length estimates. We ran the GWAS including an interaction term for genotype and age in addition to cohort, sequencing center, sex, age at sample collection, and 11 genotype PCs as covariates on Analysis Commons (Brody et al. 2017).

### Age-stratified GWAS

We divided the 109,122 TOPMed individuals with telomere length estimates into three age bins: ages 0 - 43 years old, ages 43.1 - 61 years old, and 61.1 - 98 years old. We ran the GWAS including cohort, sequencing center, sex, age at sample collection, and 11 genotype PCs as covariates on Analysis Commons (Brody et al. 2017). TOPMed cohorts included in this analysis are indicated in Supplementary Table 1. There were 36,980 individuals in the [0,43] group, 37,470 individuals in the (43,61] group, and 34,671 individuals in the (61,98] group. Any peak that cleared genome-wide significance (p<5×10^-8^) in at least one age group was considered. We then required that the lead SNP in the signal was evaluated in all three age groups. To ensure a reasonable comparison between groups, we required that the minor allele count for the SNP was at least half of the maximum group minor allele count in each group. Then we identified loci where the effect size estimate confidence interval was non-overlapping in at least one age group. Finally, we examined loci that had a genotype x age interaction p-value < 5×10^-5^ and had a meta-analysis association p-value < 5×10^-8^.

### Enrichment of meta-analysis signals in chromatin states

We estimated the enrichment of lead meta-analysis signal SNPs across each state of the 25- state chromatin state model from Roadmap Epigenomics (Roadmap Epigenomics Consortium et al. 2015) across all 127 Roadmap Epigenomics samples (Supplementary Table 13). Similarly, Roadmap Epigenomics consolidated narrowPeak files for H3K4me1 and H3K27ac from 98 and 127 samples, respectively (Supplementary Table 13), were used to compute the enrichment of lead SNPs in ChIP-seq peak regions for these histone modifications. Control SNPs were randomly selected from the genome and matched for the number of linkage disequilibrium proxy SNPs, the minor allele frequency, and the distance to the nearest gene. The same GREGOR Perl script pipeline (Schmidt et al. 2015) used to evaluate transcription factor binding site enrichment (above) was used for these analyses.

### Partitioned heritability across cell types (S-LDSC)

We limited our analysis to European individuals because the accuracy of this method depends upon an accurate match with the linkage disequilibrium reference panel. Therefore, we meta-analyzed the European individuals from two studies included in our meta-analysis (Li et al. 2020; Taub et al. 2022) using GWAMA as described above and ran stratified linkage disequilibrium score regression (S-LDSC, 1.0.1) using the cell-type specific analyses pipeline. We directly used the 1000 Genomes European baseline files, multi-tissue gene expression counts, and multi-tissue chromatin marker data generated as part of the S-LDSC pipeline (Finucane et al. 2015, 2018).

### Molecular Cloning

Gibson assembly primers were designed using Snapgene software (GSL Biotech) and sequencing primers were identified using the GenScript sequencing primer tool. All primers were synthesized by IDT. Primer sequence and a brief description of their use are provided in Supplementary Table 14. Polymerase chain reaction products were amplified using Phusion HS II DNA polymerase (F549; Thermo Fisher). Gibson Assembly was conducted using Gibson Assembly Master Mix (E2611; NEB) according to the recommended protocol. Plasmids were transformed into NEB5α cells (C2987; NEB), prepared using the QIAprep Miniprep Kit (27104; Qiagen) or the Qiagen Plasmid Midiprep Kit (12143; Qiagen), and sequence verified using the Sanger method at the Johns Hopkins School of Medicine Synthesis & Sequencing Facility.

### Overexpression constructs

Putative causal genes of interest for this experiment were required to fit three conditions: colocalization between the candidate gene GTEx eQTL and a meta-analysis signal, the lead variant at the meta-analysis signal was required to be associated with increased gene expression in the GTEx tissue where colocalization was strongest for that gene, and the gene was required to have one transcriptional isoform reported in NCBI or a coding sequence less than 15 kB, allowing it to be expressed from a plasmid. We note that *POP5* and *CBX1* had multiple transcriptional isoforms, but their transcripts result in a single, shared coding sequence. All cDNA sequences were ordered through GenScript (OHu26641, OHu13170, OHu31184, OHu26125, OHu108607) with the coding sequence subcloned into a pcDNA3.1/C-DYK vector. We added the FLAG tag to the N-or C-terminus in accordance with precedent in the literature: CBX1 C-terminus (Rosnoblet et al. 2011), PSMB4 C-terminus (Brehm et al. 2015), POP5 N-terminus (van Eenennaam et al. 2001), OBFC1 N-terminus (Bhattacharjee et al. 2016), and KBTBD6 N-terminus (Mena et al. 2018). We used Gibson Assembly to add a 3x FLAG tag to the appropriate end and insert the tagged coding sequence into a pcDNA5/FRT vector (Thermo Fisher). We note that we overexpressed the propeptide of PSMB4 (removing amino acids 2-45).

### Cell Culture

HeLa-FLP cells were generated from HeLa cells using the FLP-in system and were cultured in 1x Dulbecco’s modified Eagle’s medium (11965118; Thermo Fisher). K562 cells were purchased from ATCC (CCL-243) and were cultured in 1x RPMI medium (11875119; Thermo). Cells were cultured in the indicated media supplemented with 10% heat-inactivated fetal bovine serum (16140071; Thermo Fisher) and 1% Penicillin-Streptomycin-Glutamine (10378016; Thermo Fisher).

### Overexpression experiments and passaging

For overexpression experiments 100 ng of the indicated overexpression construct and 900 ng of the pOG44 flippase plasmid were co-transfected into HeLa-FLP cells by the use of the FLP-in system using Lipofectamine 3000 (L3000008; Invitrogen) with the recommended protocol and hygromycin resistant (550 μg/mL; 30-240-CR; Corning) cells were examined. The GFP overexpression plasmid (pAMP0605) was previously generated (Pike et al. 2019). For each construct we used one pool of HeLa-FLP cells to conduct multiple independent transfections, which we refer to as independent clones. Twice a week cells were treated with 0.05% trypsin-EDTA (25300054; Invitrogen), washed in 1x PBS (10010049; LifeTech), and counted using a Luna II Automated Cell Counter (Logos Biosystems). The number of population doublings for each passage was estimated as the number of cells counted divided by the number of cells seeded for that passage.

### Telomere Southern blot analysis

For each time point, 2-4×10^6^ cells were collected, washed in 1x PBS (10010049; LifeTech), and pellets stored at -80°C. Genomic DNA was isolated using the Promega Wizard gDNA kit (A1120; Promega) as directed. Genomic DNA was quantified using the broad range double-stranded DNA kit (Q32853; Thermo Fisher) for QuBit 3.0 (Thermo Fisher). Approximately 1 μg of genomic DNA was restricted with *Hinf*I (R0155M; NEB) and *Rsa*I (R0167L; NEB) and resolved by 0.8% Tris-acetate-EDTA (TAE) agarose gel electrophoresis. 10 ng of a 1kB Plus DNA ladder (N3200; NEB) was included on either side of the Southern as a size reference. Following denaturation (0.5 M NaOH, 1.5M NaCl) and neutralization (1.5 M NaCl, 0.5 M Tris-HCL, pH 7.4), the DNA was transferred in 10x SSC (3M NaCl, 0.35 M NaCitrate) to a Nylon membrane (RPN303B; GE Healthcare) by vacuum blotting (Boekel Scientific). The membrane was UV crosslinked (Stratagene), prehybridized in Church buffer (0.5M Na2HP04, pH7.2, 7% SDS, 1mM EDTA, 1% BSA), and hybridized overnight at 65°C using a radiolabelled telomere fragment and ladder, as previously described (Morrish and Greider 2009; S. Wang et al. 2017). The membrane was washed twice with a high salt buffer (2x SSC, 0.1% SDS) and twice with a low salt buffer (0.5X SSC, 0.1% SDS) at 65°C, exposed to a Storage Phosphor Screen (GE Healthcare), and scanned on a Storm 825 imager (GE Healthcare). The images were copied from ImageQuant TL (GE Life Sciences) to Adobe PhotoShop CS6, signal was adjusted across the image using the curves filter, and the image was saved as a .tif file. Minimum, maximum and median telomere length was estimated in ImageQuant TL using the original, unedited scan from the Phosphor Screen and accounted for differences in DNA migration across the gel by including the 1 kB Plus ladder on either side of the Southern blot.

### Western blot analysis

2×10^6^ cells were collected, washed in 1x PBS (10010049; LifeTech), resuspended in 1x sample buffer (1x NuPAGE loading buffer (NP0008; Thermo Fisher), 50 μM DTT) and stored at -80°C. Samples were thawed on wet ice, lysed by sonication, and boiled at 65°C for 10 min. Proteins were resolved using recommended parameters on 4-12% Bis-Tris NuPAGE pre-cast gels (NP0321BOX; Invitrogen) and Precision Plus Dual Color protein ladder (161-0374; BioRad) was run for comparison. Proteins were transferred to a PVDF membrane (170-4273; BioRad) using a Trans-Blot Turbo Transfer System (BioRad). The membrane was blocked in 5% milk-TBST (w/v powdered milk (170-6404; BioRad) resuspended in 1x Tris Buffered Saline, pH 7.4 (351- 086-101CS; Quality Biological), 0.01% Tween-10 (P1379-100ML; Sigma) for one hour at room temperature. Primary antibodies were diluted in blocking buffer and incubated at room temperature for one hour with mild agitation (M2 FLAG 1:2,000 (F1804-5MG; Sigma), tubulin 1:5,000 (ab6046; Abcam)). Blots were washed in 1x TBST with mild agitation before incubation with horseradish peroxidase-conjugated secondary antibodies diluted in blocking buffer (α- mouse 1:10,000 (170-6516; BioRad), α-rabbit 1:10,000 (170-6515; BioRad)). Blots were washed in 1x TBST with mild agitation, incubated with Forte horseradish peroxidase substrate (WBLUF0100; Millipore) for five minutes with agitation, and imaged on an ImageQuant LAS 4000 mini biomolecular imager (GE Healthcare). Image files were copied from ImageQuant TL software to Adobe PhosShop CS6, the curves filter was applied across the image, and then saved as a .tif file. To reprobe a membrane with the loading control, the membrane was incubated with Restore Western Blot Stripping Buffer (21059; Thermo Fisher) for 30 minutes, washed in 1x TBST, and processed as described above.

### CRISPR editing constructs

We sequence verified the CRISPR target regions in our K562 cells and selected gRNA sequences with a high likelihood of on-target editing (and a low likelihood of off-target editing) using CRISPOR.org (Concordet and Haeussler 2018). We subcloned the guides into px458 as previously described (Moyer and Holland 2015). To edit both the *POP5* and *KBTBD6/KBTBD7* regions we chose one guide to each side of the target region (Supplementary Figure 8C-D). For guide sequence and genome coordinates (hg38), see Supplementary Table 14.

### CRISPR editing experiments

Low-passage K562 cells were cultured to a density of 3×10^5^ cells/mL in media without antibiotics, but otherwise as described above, two days prior to nucleofection. Cells were electroporated using the SF Cell Line 4D-Nucleofector X Kit (V4XC-2012; Lonza) with 8 μg of each guide plasmid and the K562 cell line recommended protocol (FF-120). Cells were cultured in antibiotic-free media for 24 hours to allow for GFP expression before being single-cell sorted in a 96 well plate at the Johns Hopkins Ross Flow Cytometry Core. Each sample had 1-10% GFP positive cells. Plates were expanded clonally using media described above. After approximately two weeks cell concentration was estimated using the Luna II Automated Cell Counter (Logos Biosystems), 4×10^4^ cells were collected, and genomic DNA was extracted using QuickExtract DNA Extraction Solution (QE09050; Epicentre) following the protocol recommended in the Alt-R genomic editing detection kit (1075931; IDT). Target editing regions were amplified (primers described in Supplementary Table 14, diagrams in Supplementary Figure 8) and confirmed by Sanger sequencing. Sequencing reads were aligned in Snapgene (GSL Biotech) and we considered a clone to at least be heterozygous for editing if the alignment began on one side of the deletion, failed across the intended deletion, but resumed across the deletion. Because the *POP5* locus deletion was so extensive, we did two separate PCRs on each sample: one that would amplify if the deletion was present (RK236+RK231) and one that would amplify if a wildtype allele was present (RK236+RK234) (Supplementary Figure 8C). All *POP5* edited clones were confirmed to be heterozygous.

### RNA extraction and qPCR

2×10^6^ cells were collected, washed in 1x PBS (10010049; LifeTech), and RNA was purified using a QIAshreddar column (79656; Qiagen) and RNeasy kit (74104; Qiagen) following the recommended protocols, including DNase digestion of RNA prior to RNA cleanup (79254; Qiagen). RNA concentration was estimated using a high sensitivity RNA kit (Q32852; Thermo Fisher) for QuBit 3.0 (Thermo Fisher). cDNA was generated with random hexamers using a SuperScript IV First Strand Synthesis kit (18091050; Thermo Fisher). qPCR primers were designed using the GenScript RT-PCR primer design tool and a standard reference plasmid was generated by amplifying genomic DNA from K562 cells with each primer pair followed by TA cloning the amplicon into a pCR2.1 vector (Supplementary Table 14) using a TA cloning kit (451641; Thermo Fisher). TA cloning was conducted using the recommended protocol and plasmids were transformed into TOP10 cells (C404003; Invitrogen). Each qPCR reaction included approximately 10 ng of cDNA, 1x iQ SYBER Green Super Mix (1708882; BioRad), and 0.25 μM of each primer; qPCR was conducted on a CFX96 real-time qPCR system (BioRad). *KBTBD6* and *KBTBD7* expression was measured in the *POP5*-edited clones as CRISPR/Cas9-edited controls and *POP5* expression was measured in the *KBTBD6*/*KBTBD7*-edited clones as CRISPR/Cas9-edited controls. Samples were analyzed in triplicate and instances where the Cq range was greater than 1 were excluded from further analysis. Standard plasmids were analyzed in duplicate on each plate at a range of 0.001 ng - 100 ng as a quality control measure and plates where the standards Cq had an R^2^ < 0.98 were excluded from further analysis. Plates that passed this threshold were used to estimate the efficiency of the qPCR primers (*ACTB* = 1.90, *KBTBD6* = 1.98, *KBTBD7* = 1.92, and *POP5* = 1.80). Because the range of efficiency between measured genes was greater than 10%, we analyzed our qPCR results with the Pfaffl method (Pfaffl 2001). A one-sided *t*-test was used to compare experimental to control samples.

### Data and code availability

All cell lines and plasmids are available upon request. Summary statistics, plasmid maps, and code are available at Zenodo (doi: 10.5281/zenodo.8136834) and are freely available. Additional code is available here: https://github.com/BennyStrobes/leafcutter_sqtl_viz, https://github.com/bulik/ldsc, https://github.com/stephenslab/susieR. Any additional information required to reanalyze the data reported here is available upon request. TOPMed genomic data and telomere length estimates are available by study in the database of Genotypes and Phenotypes (dbGaP) (https://www.ncbi.nlm.nih.gov/gap/?term=TOPMed). GTEx_v8 eQTL, sQTL, and LeafCutter exon-exon junction quantifications are available for download through the GTEx portal (https://gtexportal.org/home/). eQTLGen cis-eQTL data are available for download (https://www.eqtlgen.org/). DICE cis-eQTL data are available for download (https://dice-database.org/landing).

## Supplemental Note

This supplemental note conveys the rationale for the assigned putative causal gene for each signal. For 17 signals no colocalization results were available and there were no known genes involved in telomere length in the region (Supplemental Note Table 1, see rows with “Proximal gene (no other supporting information)”). In these cases, the proximal gene was assigned. For 33 signals there was colocalization data in at least one QTL dataset (Figure 2B). However, colocalization within and between datasets supported different genes for 14 meta-analysis signals (Discussed below). For each signal we show the Manhattan plot for the meta-analysis signal and the best colocalization result for each gene in each dataset. For datasets with multiple cellular contexts and in cases where the meta-analysis signal colocalized with a QTL for the same gene across cellular contexts, we show the QTL that had the highest PPH4. We considered PPH4 > 0.7 for GTEx and eQTLGen, PPH4 > 0.5 for DICE to be colocalized, all colocalization results are reported in Supplementary Tables 3-6. The meta-analysis Manhattan plots were centered on the lead SNP and include the region ± 1 Mb the lead SNP (hg38) and the x-axis is matched for each plot. In all plots the meta-analysis lead SNP was shown as a black diamond and r^2^ was calculated with respect to the lead SNP using all TOPMed individuals included in the meta-analysis.

**Supplemental Note Table 1:**

| Lead SNP | Novel Signal | Attributed gene | Supporting evidence for attributed gene |
| --- | --- | --- | --- |
| rs542948485 |  | DCLRE1B | Known biology, colocalization |
| rs12044242 |  | PSMB4 | Colocalization, proximal gene |
| rs146042055 |  | PARP1 | Known biology |
| rs62139251 |  | TSPYL6 | Colocalization |
| rs11894326 |  | CPS1 | Colocalization |
| rs35510081 |  | TERC | Known biology |
| rs3775946 |  | SLC2A9 | Colocalization, proximal gene |
| rs4691895 |  | NAF1 | Known biology, colocalization |
| rs33961405 |  | TERT | Known biology |
| rs56099285 |  | UBE2D2 | Proximal gene (no other supporting information) |
| rs3131064 | * | POU5F1 | Colocalization, proximal gene |
| rs1150748 |  | BAG6 | Colocalization |
| rs6968500 |  | POT1 | Known biology |
| rs10954213 |  | IRF5 | Colocalization, proximal gene |
| rs3008267 | * | ZNF596 | Proximal gene (no other supporting information) |
| rs10958468 |  | TMEM68 | Proximal gene (no other supporting information) |
| rs73687065 |  | TERF1 | Known biology, proximal gene |
| rs10111287 |  | VIRMA | Colocalization, proximal |
| rs62560860 |  | IRL11A | Colocalization |
| rs958919990 | * | GRHPR | Proximal gene (no other supporting information) |
| rs3736462 |  | TASOR2 | Proximal gene (no other supporting information) |
| rs3758526 |  | NOC3L | Proximal gene (no other supporting information) |
| rs7923385 | * | RRP12 | Proximal gene (no other supporting information) |
| rs11190126 |  | NKX2-3 | Proximal gene (no other supporting information) |
| rs112519582 | * | BTRC | Proximal gene (no other supporting information) |
| rs2475215 |  | OBFC1 | Known biology |
| rs12241155 | * | SORCS1 | Proximal gene (no other supporting information) |
| rs582297 |  | ATM | Known biology, Colocalization, Proximal gene |
| rs74892322 |  | POP5 | Known biology |
| rs28755851 |  | ZCCHC8 | Known biology |
| rs1411041 |  | KBTBD6/KBTBD7 | Colocalization, Proximal gene |
| rs532687339 |  | TINF2 | Known biology |
| rs4902358 |  | MAX | Colocalization, proximal |
| rs2572 |  | DCAF4 | Proximal gene (no other supporting information) |
| rs11623533 |  | PPP4R3A | Proximal gene (no other supporting information) |
| rs2887399 |  | TCL1A | Colocalization, proximal gene |
| rs113119217 |  | ATP8B4 | Colocalization, proximal gene |
| rs12934863 |  | DUS2 | Colocalization, proximal gene |
| rs9939870 |  | TERF2 | Known biology, colocalization, proximal gene |
| rs12149396 |  | CLEC18C | Colocalization |
| rs7193541 |  | RFWD3 | Colocalization, proximal gene |
| rs6564996 |  | MPHOSPH6 | Colocalization, proximal gene |
| rs11117354 |  | BANP | Proximal gene (no other supporting information) |
| rs59922886 |  | CTC1 | Known biology, colocalization, proximal gene |
| rs208011 | * | SKAP1 | Proximal gene (no other supporting information) |
| rs144204502 |  | TK1 | Colocalization, proximal gene |
| rs2124616 |  | TYMSOS | Colocalization |
| rs28782011 |  | SETBP1 | Proximal gene (no other supporting information) |
| rs139955496 |  | POLI | Proximal gene (no other supporting information) |
| rs8105767 |  | ZNF257 | Colocalization, proximal gene |
| rs79476302 |  | SAMHD1 | Proximal gene (no other supporting information) |
| rs114703330 |  | RTEL1 | Known biology, colocalization, proximal gene |
| rs28663120 |  | GAB4 | Colocalization, proximal gene |
| rs131784 |  | TYMP | Colocalization |
| rs12394264 |  | MIR223HG | Colocalization, proximal gene |
| rs5945232 |  | DKC1 | Known biology |

**rs542948485 (chr1:113917053:G:T)**

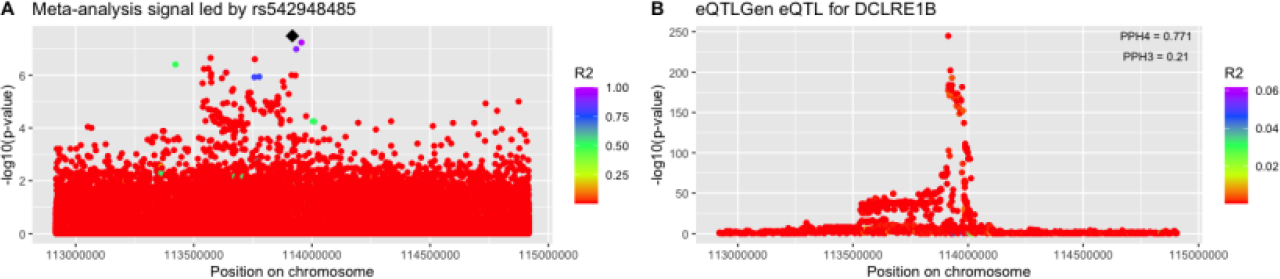

This meta-analysis signal colocalized with a *DCLRE1B* eQTL in eQTLGen. *DCLRE1B* is also the proximal gene. In addition, *DCLRE1B* is known to contribute to telomere length regulation. Therefore, we concluded that *DCLRE1B* was the best supported putative casual gene.

**rs12044242 (chr1:151398465:C:T)**

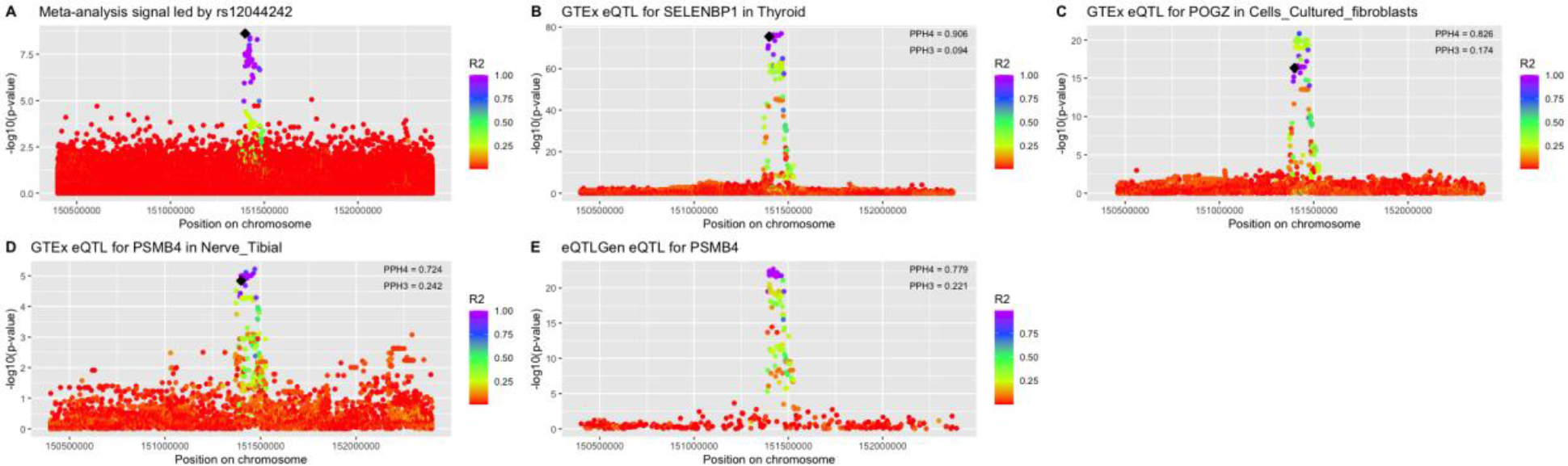

This meta-analysis signal colocalized with QTLs for *PSMB4*, *POGZ*, and *SELENBP1*. We observed that the *PSMB4* eQTL colocalization was replicated in eQTLGen. *PSMB4* is also the proximal gene for this signal, therefore we concluded that *PSMB4* was the best supported putative causal gene.

**rs62139251 (chr2:54251468:G:T)**

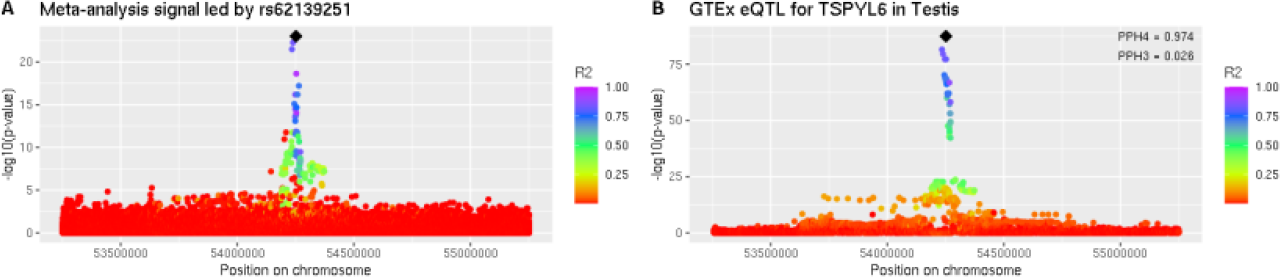

This meta-analysis signal only colocalized with a *TSPYL6* eQTL. The proximal gene was *ACYP2.* We concluded that *TSPYL6* was the best supported putative causal gene.

**rs11894326 (chr2:209808365:C:T)**

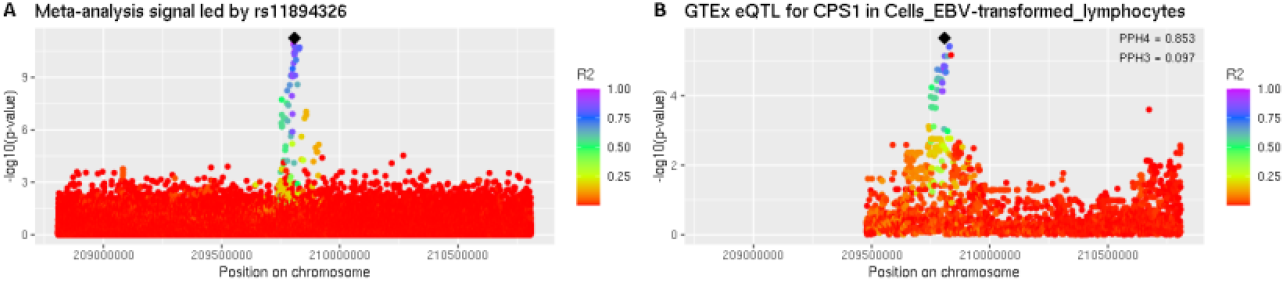

This meta-analysis signal only colocalized with a *CPS1* QTL. The proximal gene was *UNC80*. We concluded that *CPS1* was the best supported putative causal gene.

**rs3775946 (chr4:9993632:A:G)**

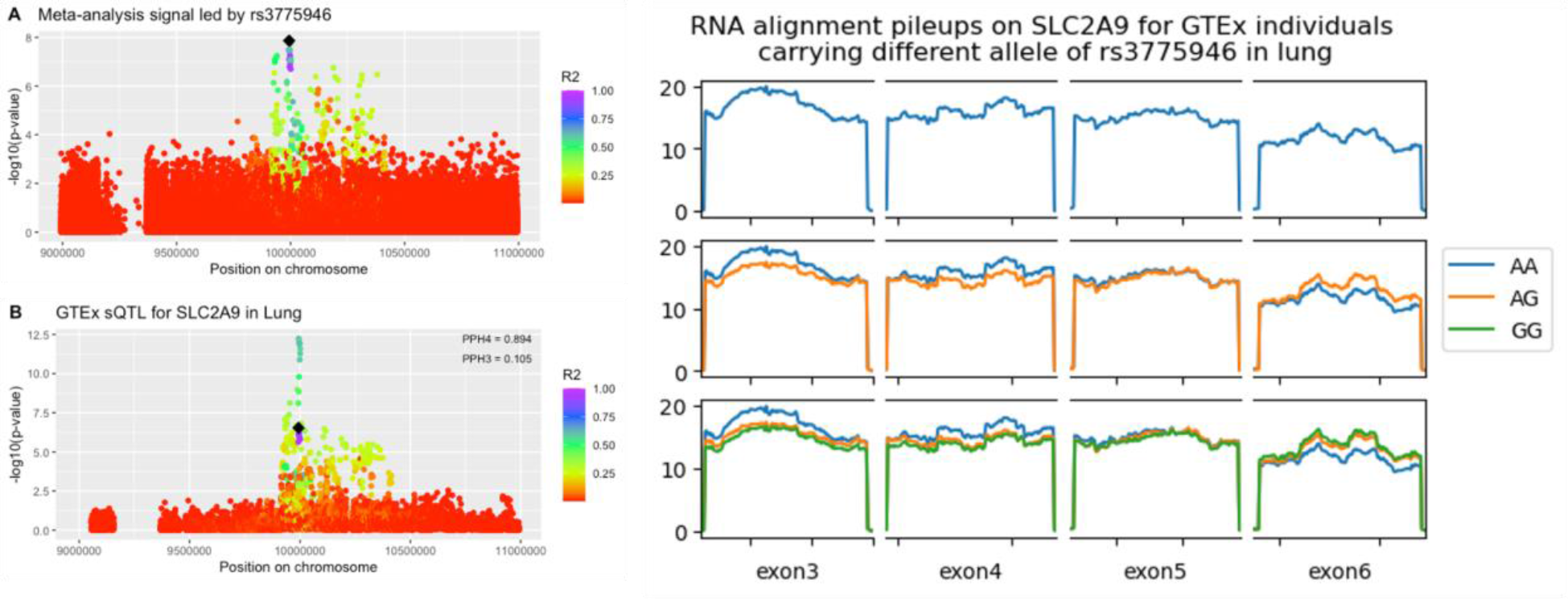

The meta-analysis signal colocalized with a *SLC2A9* sQTL. The RNA pileup plot shows the aligned reads in the indicated GTEx tissue for the indicated exons that were included in the LeafCutter splicing cluster. Unlike an eQTL, a subset of exons show differences in the amount of reads aligned when stratified by the indicated genotype, supporting that this is a sQTL. *SLC2A9* was also the proximal gene, therefore we concluded that *SLC2A9* was the best supported putative causal gene.

**rs4691895 (chr4:163127047:G:C)**

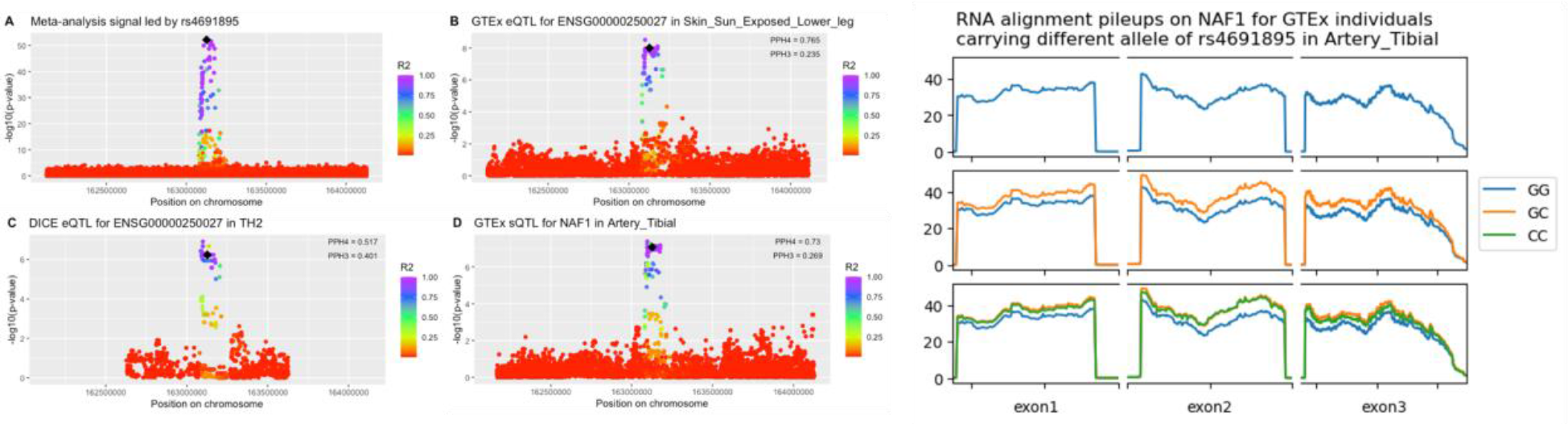

The only characterized gene QTL this meta-analysis signal colocalized with was a *NAF1* sQTL. *NAF1* is also a known telomere regulation gene and the gene proximal to the signal. The RNA pileup plot shows the aligned reads in the indicated GTEx tissue for the indicated exons that were included in the LeafCutter splicing cluster. Unlike an eQTL, a subset of exons show differences in the amount of reads aligned when stratified by the indicated genotype, supporting that this is a sQTL.Therefore, we concluded that *NAF1* was the best supported putative causal gene.

**rs3131064 (chr6:30796116:T:C)**

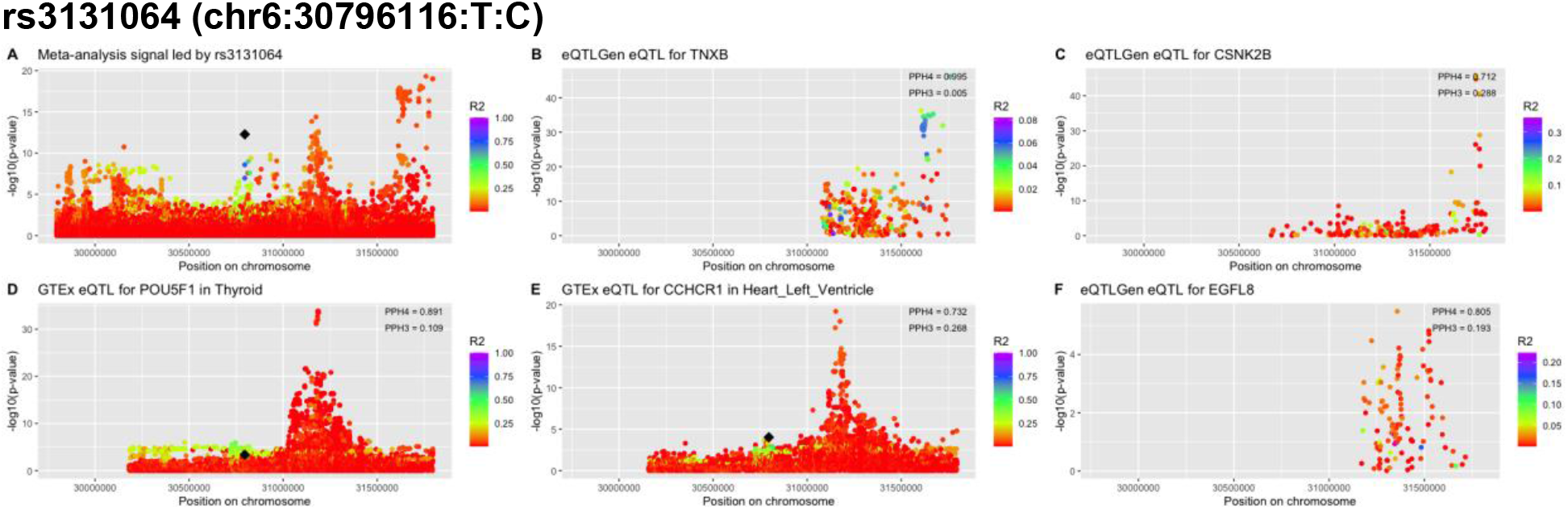

This meta-analysis signal is near the *HLA* locus and there may be several independent signals within this region. We defined signals based on position alone, therefore we are treating this region as a single signal. rs3131064 was considered the lead SNP because lead SNPs were chosen by ordering the genome-wide significant SNPs by p-value, selecting the top SNP within a 1Mb region, and then removing any other SNPs within 1Mb of that top SNP. The peaks adjacent to rs3131064 within panel A are within 1Mb of rs1150748 and were therefore excluded from being considered the lead SNP for this signal. One of these adjacent signals, led by rs1265156, colocalized with with QTLs for *POU5F1* and *CCHCR1*. The proximal gene was *HCG20*. We concluded that *POU5F1* was the best supported putative causal gene for this region.

**rs1150748 (chr6:31804139:G:C)**

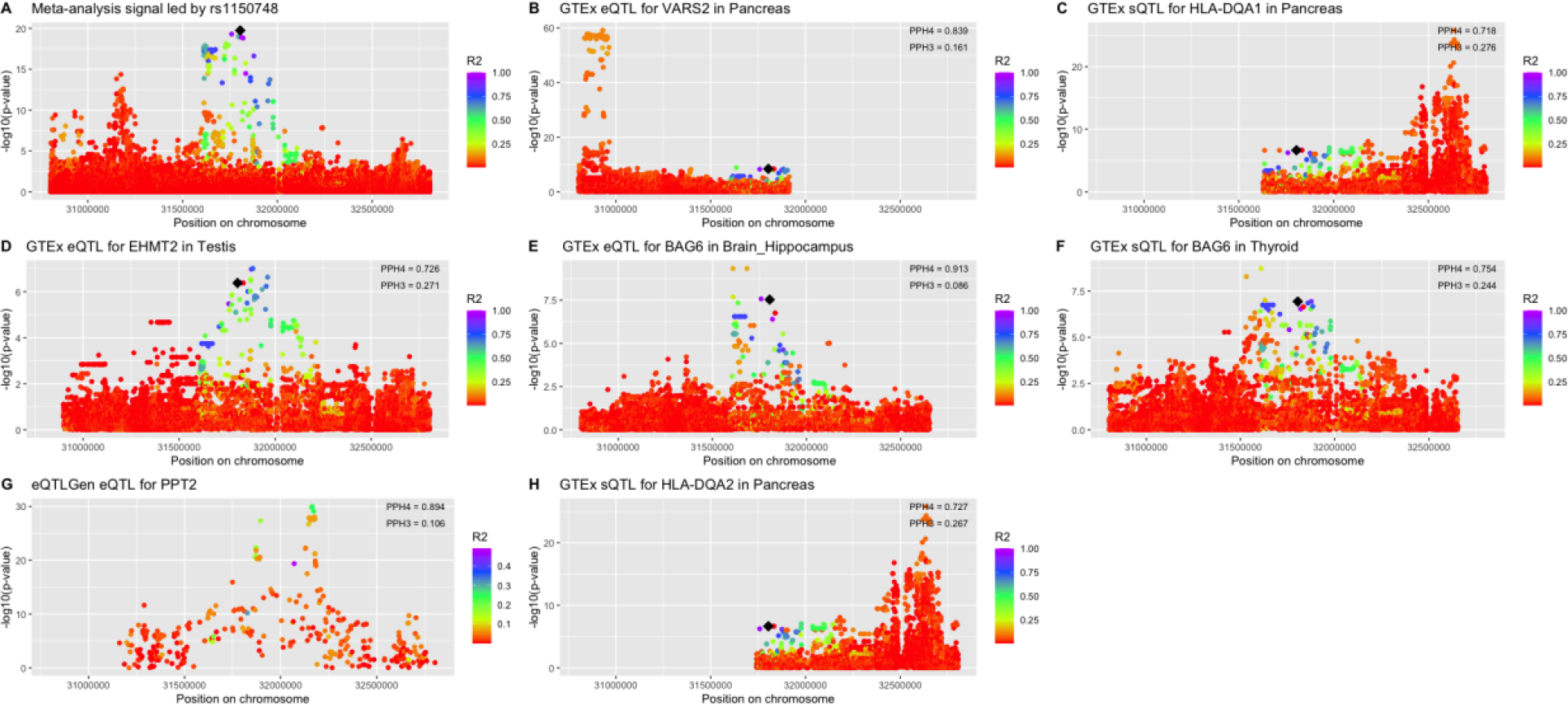

This meta-analysis signal is near the *HLA* locus and there may be several independent signals within this region. We defined signals based on position alone, therefore we are treating this region as a single signal. rs1150748 was considered the lead SNP because lead SNPs were chosen by ordering the genome-wide significant SNPs by p-value, selecting the top SNP within a 1Mb region, and then removing any other SNPs within 1Mb of that top SNP. This signal colocalized well with several gene QTLs but the association signal structure was best captured by the *BAG6* QTL. *LSM2* was the proximal gene. We concluded that *BAG6* was the best supported putative causal gene.

**rs6968500 (chr7:124791668:G:C)**

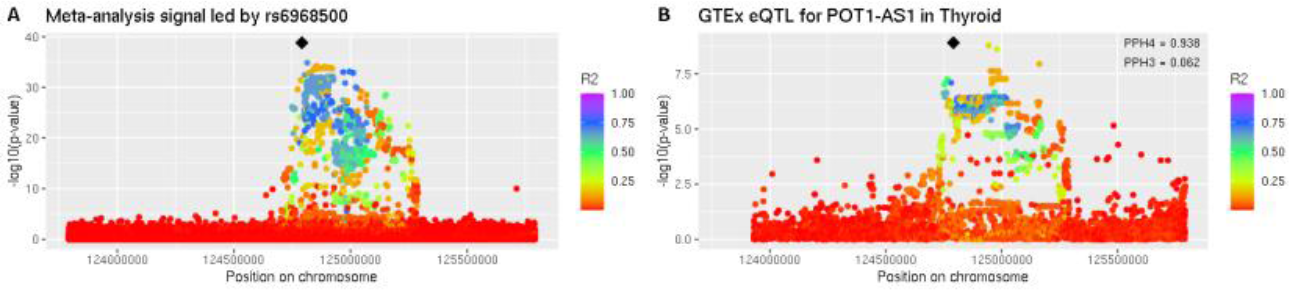

This meta-analysis signal colocalized with a QTL for *POT1-AS1*. The proximal gene was *C7orf77*. *POT1*, 30 kb away from the lead SNP, has known roles in telomere length regulation. Therefore, we concluded that *POT1* was the most likely putative causal gene.

**rs10954213 (chr7:128949373:A:G)**

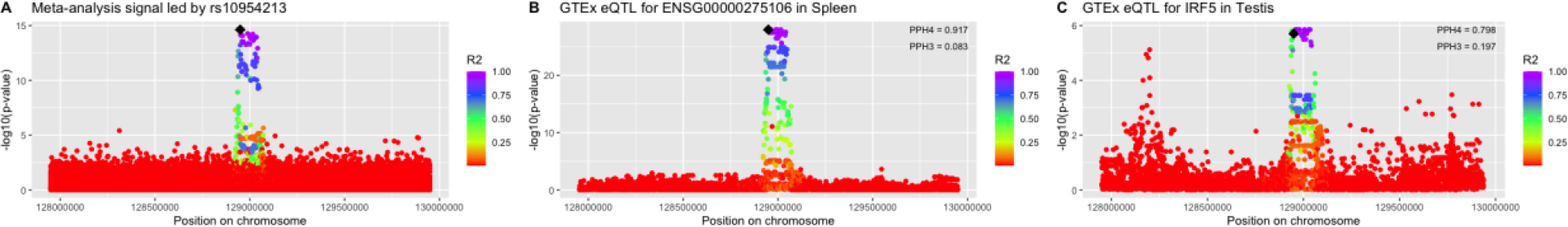

The only characterized gene QTL this meta-analysis signal colocalized with was *IRF5*, which is also the proximal gene. Therefore, we concluded that *IRF5* was the best supported putative causal gene.

**rs10111287 (chr8:94566198:C:T)**

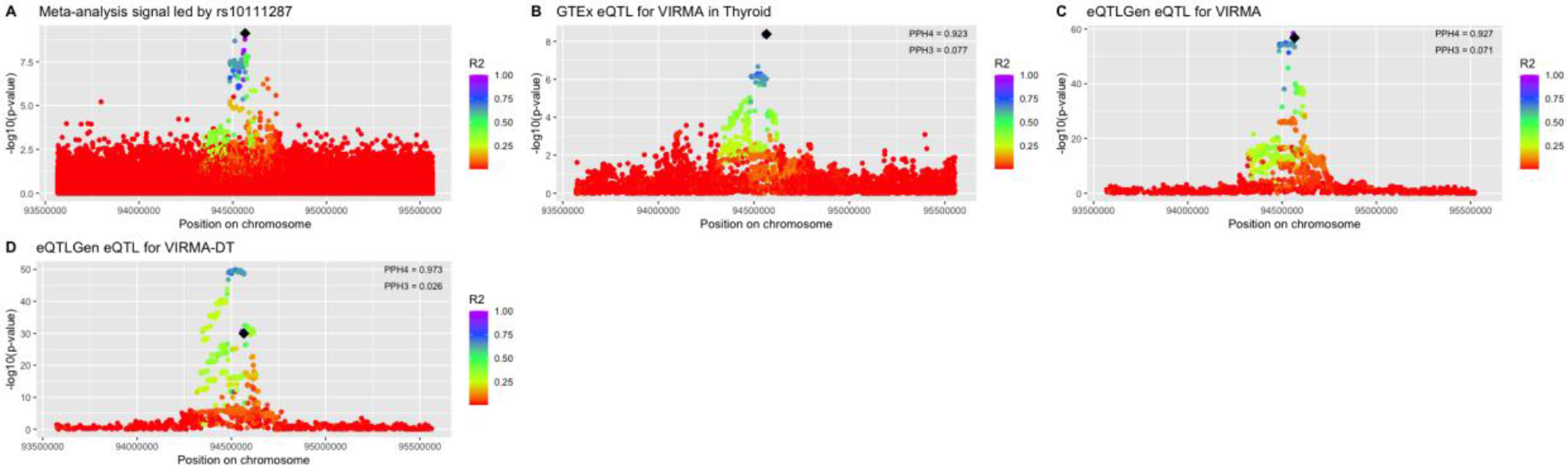

This meta-analysis signal best colocalized with *VIRMA* QTLs. *VIRMA* is also the proximal gene. Therefore, we concluded that *VIRMA* is the best supported putative causal gene.

**rs62560860 (chr9:34077464:G:A)**

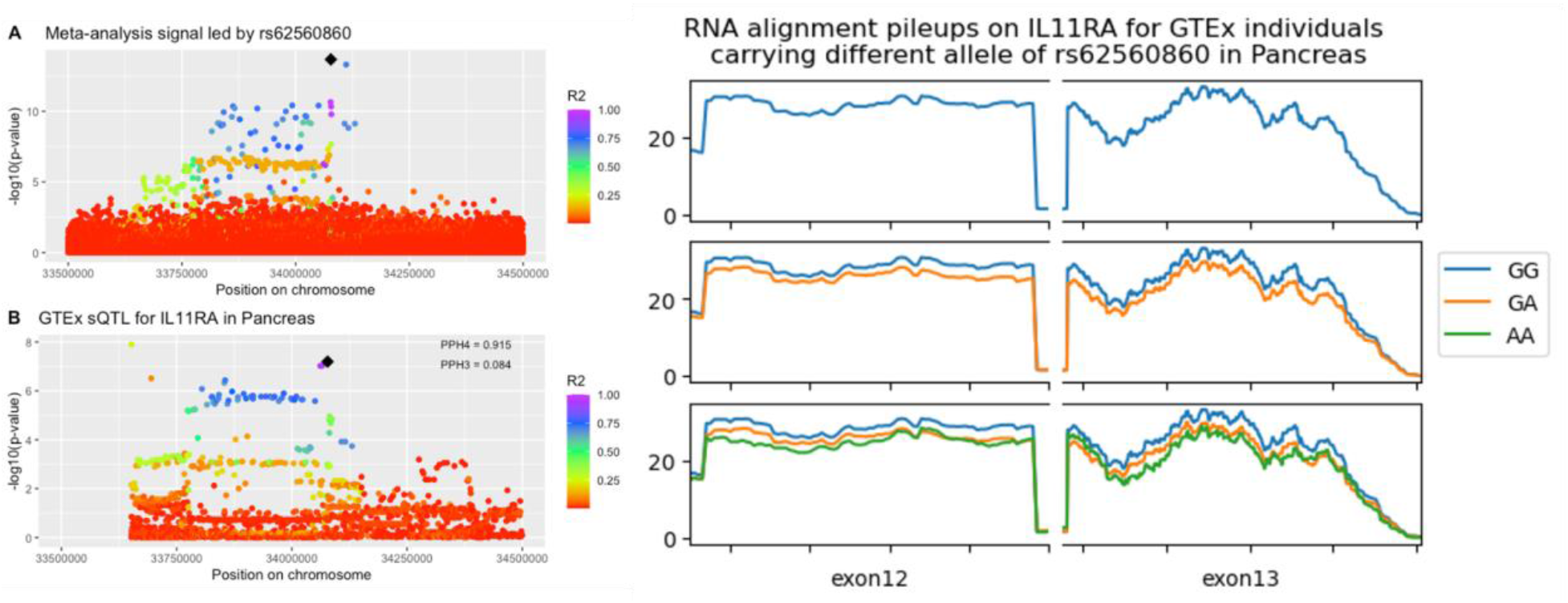

This meta-analysis signal only colocalized with an *IL11RA* sQTL. The proximal gene was *DCAF12*. Note that plot is 1 Mb wide instead of 2 Mb wide to improve visualization of the sQTL because there is a nearby SNP, rs11575580, that has a strong association (p=3.04×10^-119^) but r^2^ with meta-analysis lead SNP = 0.0169 and did not contribute to the colocalization signal. To improve clarity, we reduced the plot region to 1 Mb centered on the meta-analysis lead SNP. The RNA pileup plot shows the aligned reads in the indicated GTEx tissue for the indicated exons that were included in the LeafCutter splicing cluster. Unlike an eQTL, a subset of exons show differences in the amount of reads aligned when stratified by the indicated genotype, supporting that this is a sQTL.Given the colocalization analysis results we concluded that *ILR11A* was the most supported putative causal gene.

**rs2475215 (chr10:103900944:T:C)**

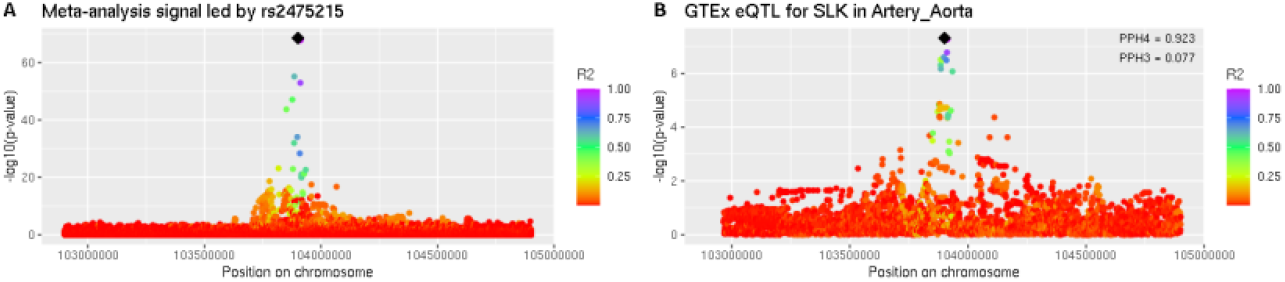

This meta-analysis signal best colocalized with an *SLK* QTL. However, *OBFC1* is a known telomere length regulation gene located 17 kB away. Given the known biology, we concluded that *OBFC1* was the most likely putative causal signal.

**rs582297 (chr11:108294680:C:G)**

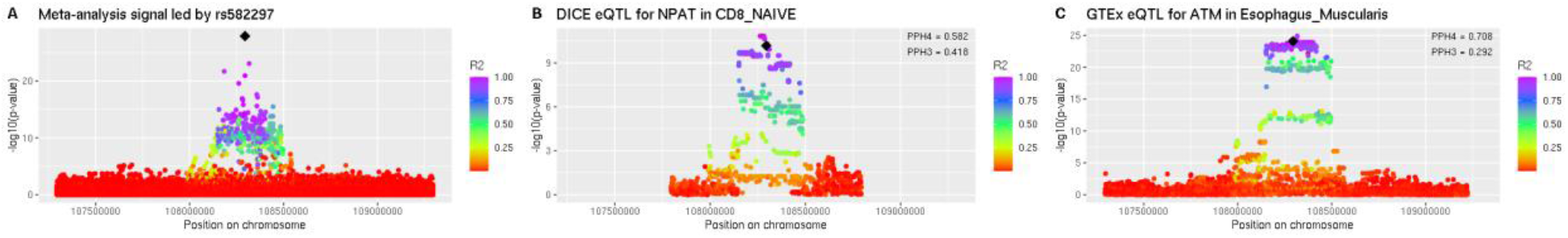

This meta-analysis signal colocalized with QTLs for *NPAT* and *ATM*. *ATM* was the proximal gene and has known roles in telomere length regulation. Therefore, we concluded that *ATM* was the best supported putative causal gene.

**rs74892322 (chr12:120533371:A:T)**

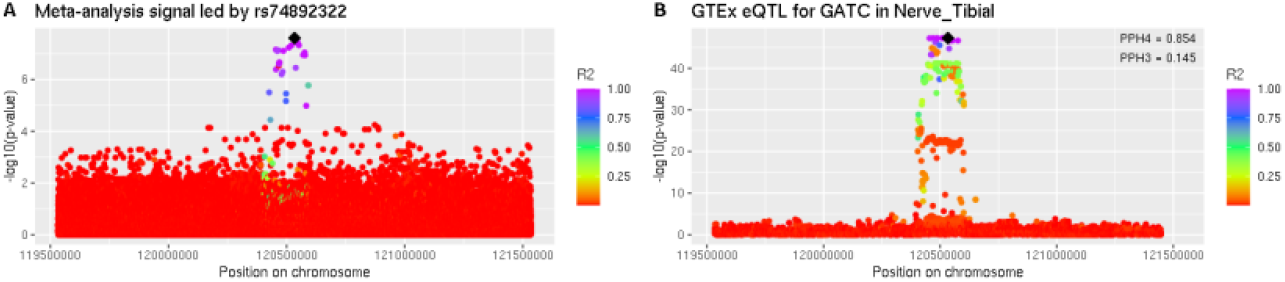

This meta-analysis signal colocalized with QTLs for *GATC* and *POP5*. The proximal gene was *RNF10*. This signal colocalized with a *POP5* eQTL in GTEx nucleus accumbens basal ganglia, but was below the threshold for being included in these plots (Supplementary Table 3). Given our results from Figure 6, we concluded that the best supported putative causal gene was *POP5*.

**rs28755851 (chr12:123001735:A:T)**

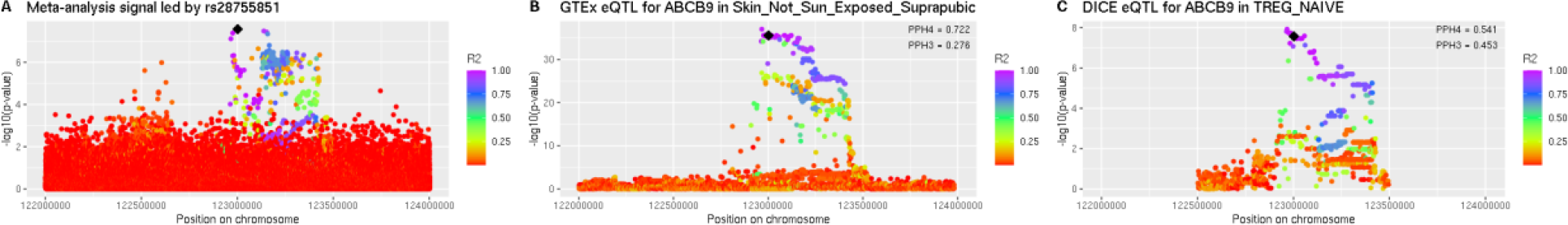

This meta-analysis signal colocalized with an *ABCB9* eQTL in GTEx and this was replicated in DICE. The proximal gene was *PITPNM2*. However, *ZCCHC8* has known roles in telomere length regulation and is 530 kB away. We concluded that *ZCCHC8* was the most likely putative causal gene despite lack of colocalization.

**rs1411041 (chr13:41150640:A:T)**

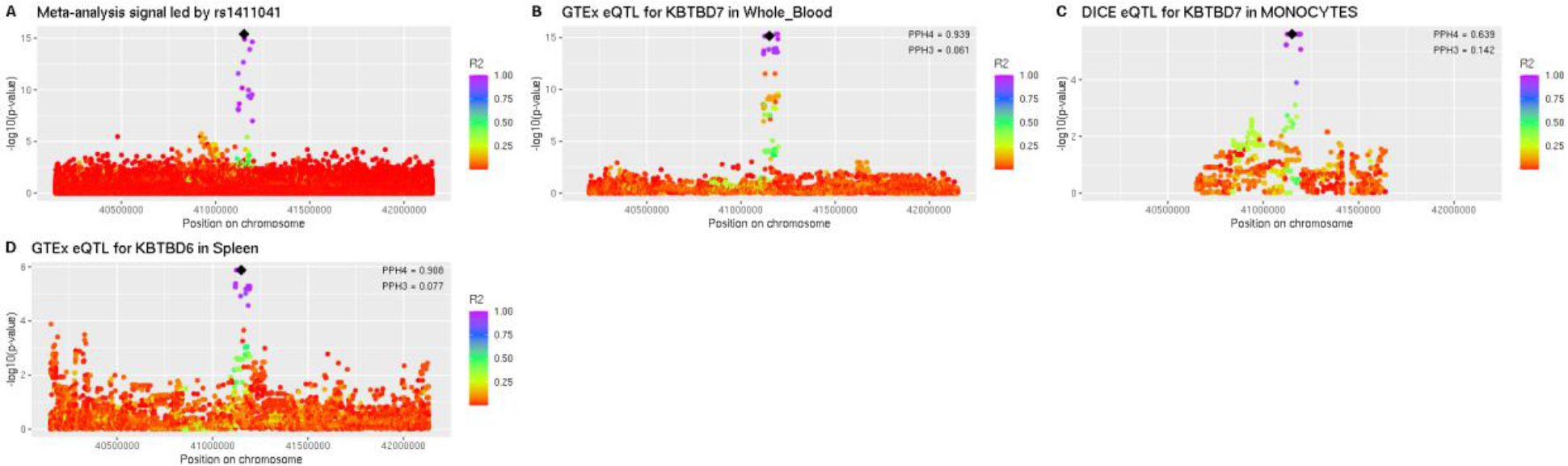

This meta-analysis signal colocalized well with *KBTBD6* and *KBTBD7* QTLs. *KBTBD6* is the proximal gene (23 kB from the lead SNP whereas *KBTBD7* is 39 kB). Particularly considering our validation experiments, we are unable to choose a single putative causal gene for this signal. We label the Manhattan plot in Figure 1 *KBTBD6* for clarity.

**rs4902358 (chr14:65075759:A:G)**

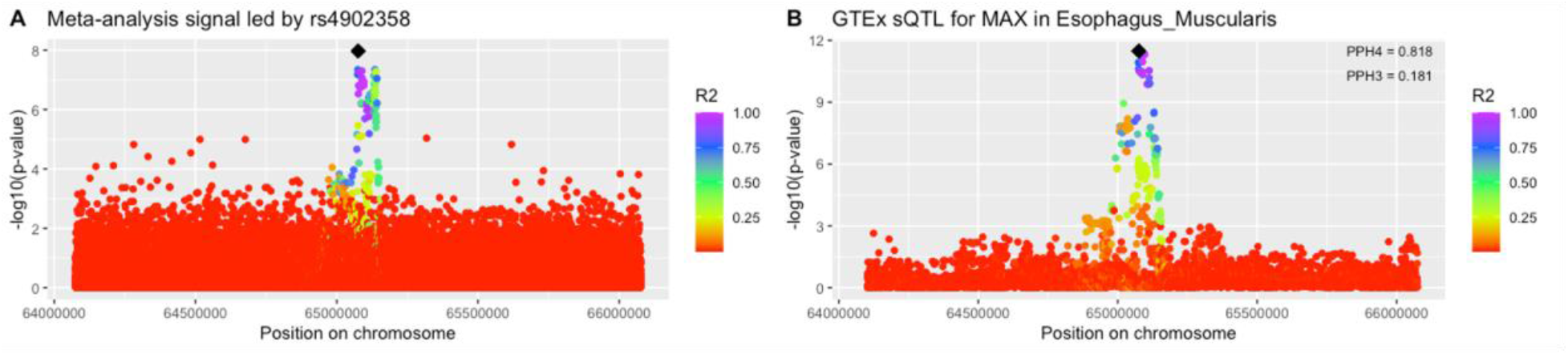
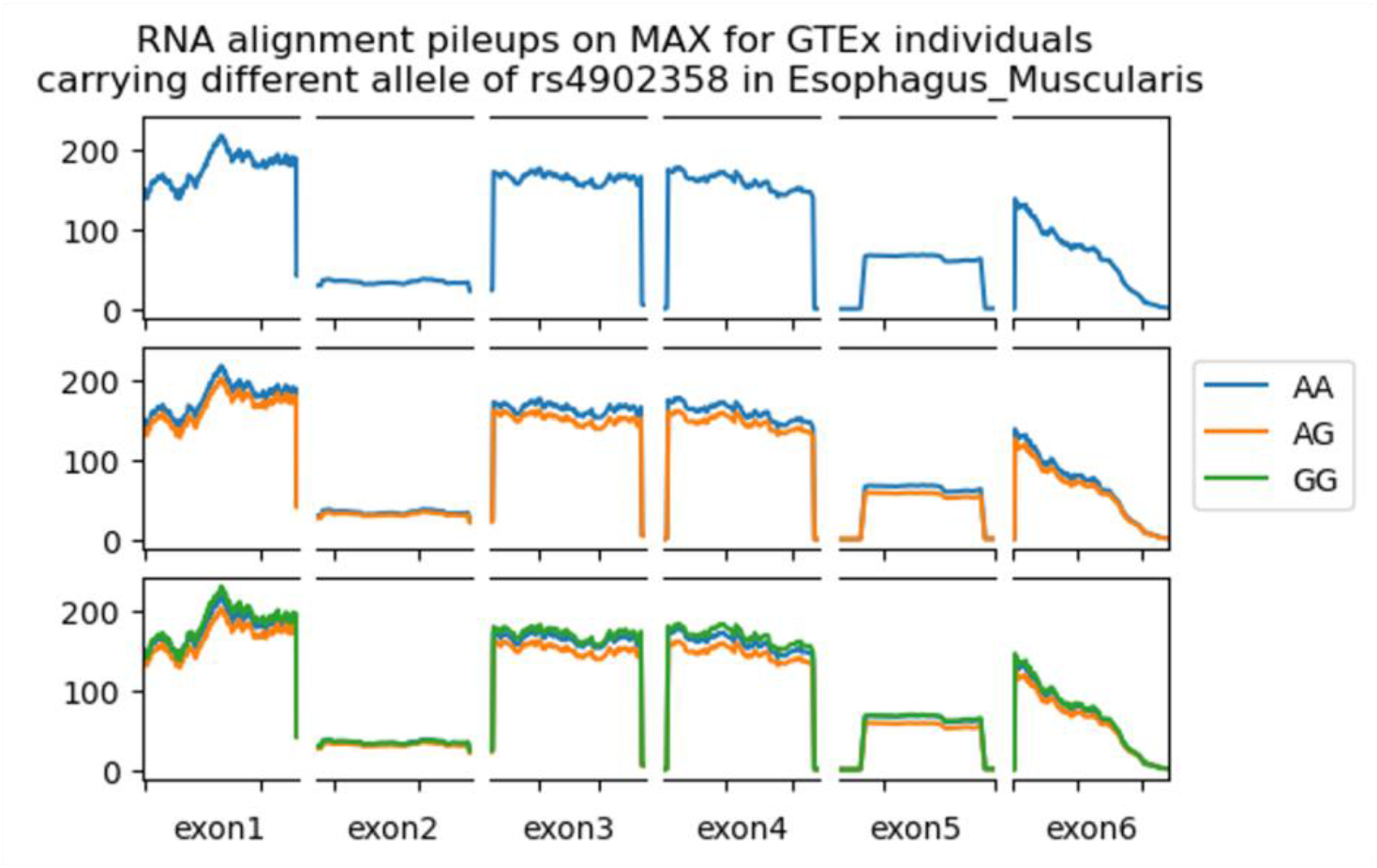

This meta-analysis signal best colocalized with *MAX* QTLs. It is also the proximal gene. The RNA pileup plot shows the aligned reads in the indicated GTEx tissue for the indicated exons that were included in the LeafCutter splicing cluster. Unlike an eQTL, a subset of exons show differences in the amount of reads aligned when stratified by the indicated genotype, supporting that this is a sQTL.Therefore, we concluded that *MAX* was the best supported putative causal gene.

**rs2887399 (chr14:95714358:G:T)**

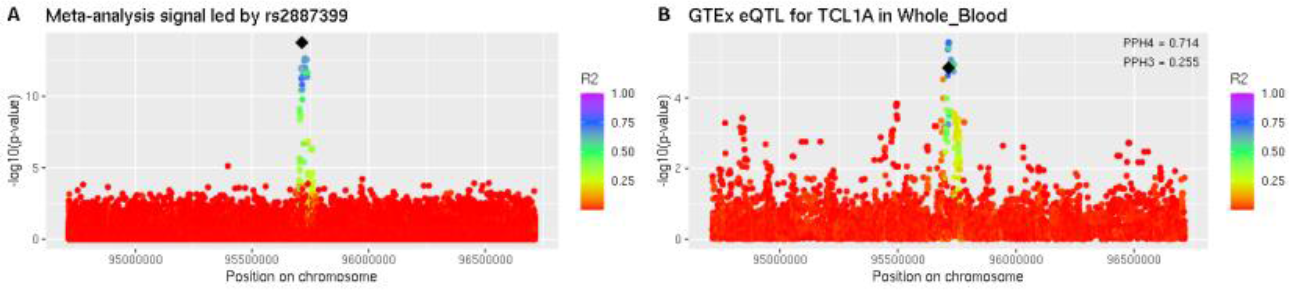

This meta-analysis signal best colocalized with *TCL1A* QTLs. *TCL1A* was also the proximal gene. Therefore, we concluded that *TCL1A* was the best supported putative casual gene.

**rs113119217 (chr15:50073451:T:A)**

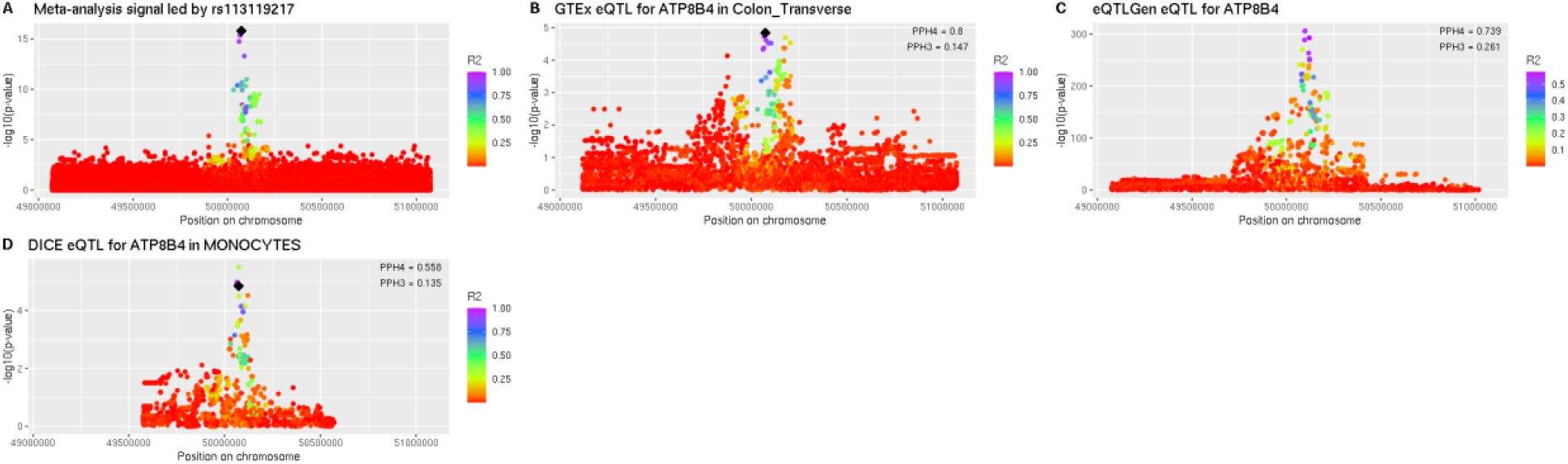

This meta-analysis signal best colocalized with QTLs for *ATP8B4* in multiple QTL datasets. *ATP8B4* is also the proximal gene. Therefore, we concluded that *ATP8B4* was the best supported putative causal gene.

**rs12934863 (chr16:68043168:A:G)**

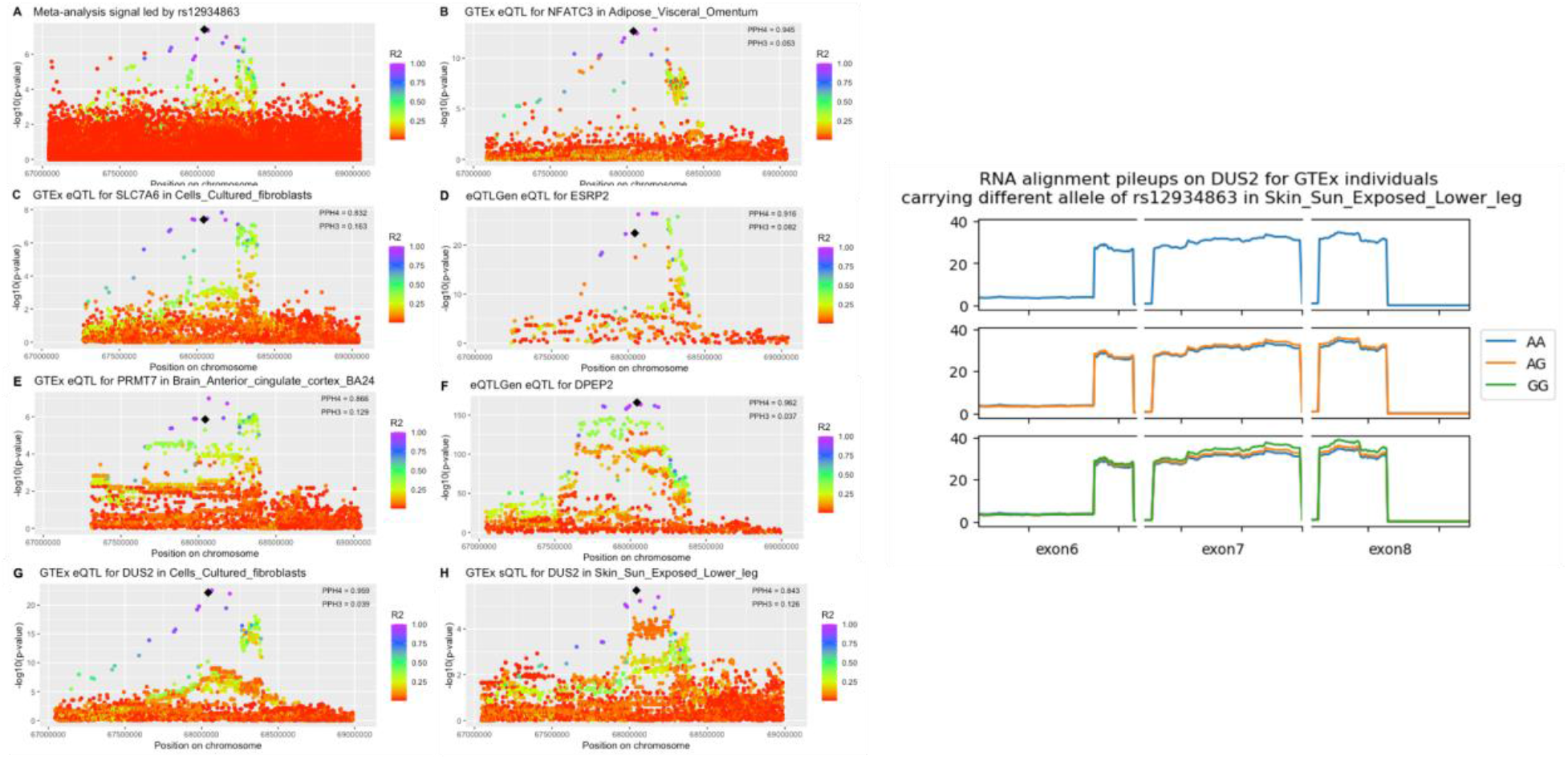

This meta-analysis signal colocalized strongly with QTLs for *DUS2*, *NFATC3*, and *DPEP2*. The proximal gene was *DUS2*. The RNA pileup plot shows the aligned reads in the indicated GTEx tissue for the indicated exons that were included in the LeafCutter splicing cluster. Unlike an eQTL, a subset of exons show differences in the amount of reads aligned when stratified by the indicated genotype, supporting that this is a sQTL. We concluded that the best supported putative causal gene was *DUS2*.

**rs9939870 (chr16:69362682:C:T)**

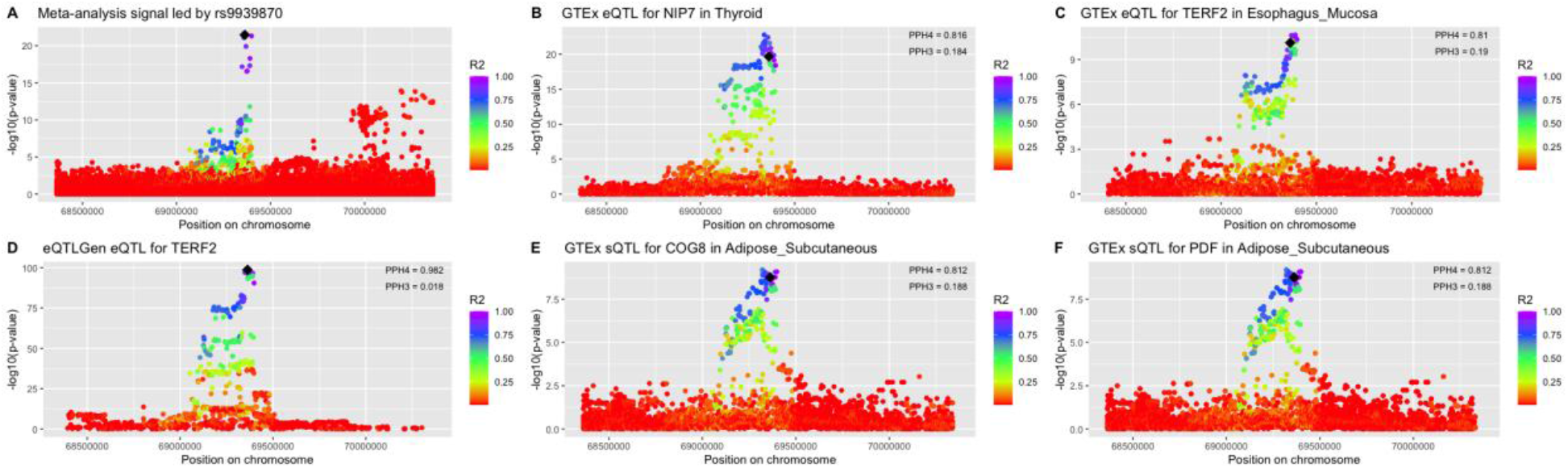

This meta-analysis signal colocalized strongly with QTLs for *TERF2*, *NIP7*, *COG8*, and *PDF*. The proximal gene was *TERF2* and *TERF2* has known roles in telomere length regulation. Therefore, we concluded that *TERF2* is the best supported putative causal gene.

**rs12149396 (chr16:70392835:A:C)**

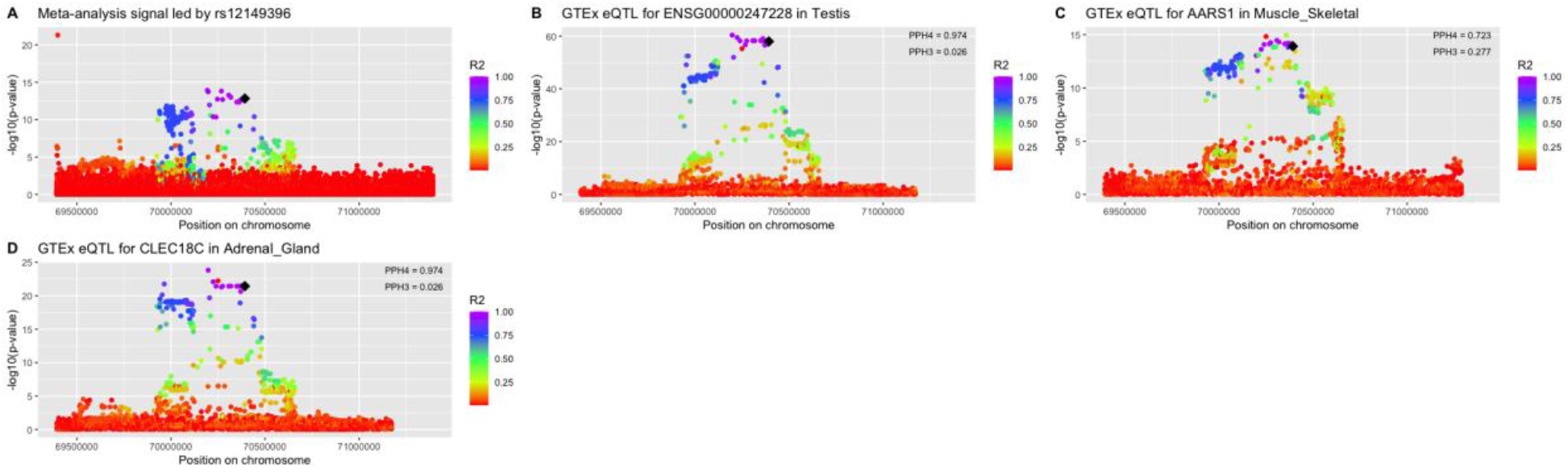

This meta-analysis signal colocalized with *AARS1* and *CLEC18C* QTLs. The proximal gene was *ST3GAL2*. Based on the strength of colocalization, we concluded that *CLEC18C* was the best supported putative causal gene.

**rs7193541 (chr16:74630845:T:C)**

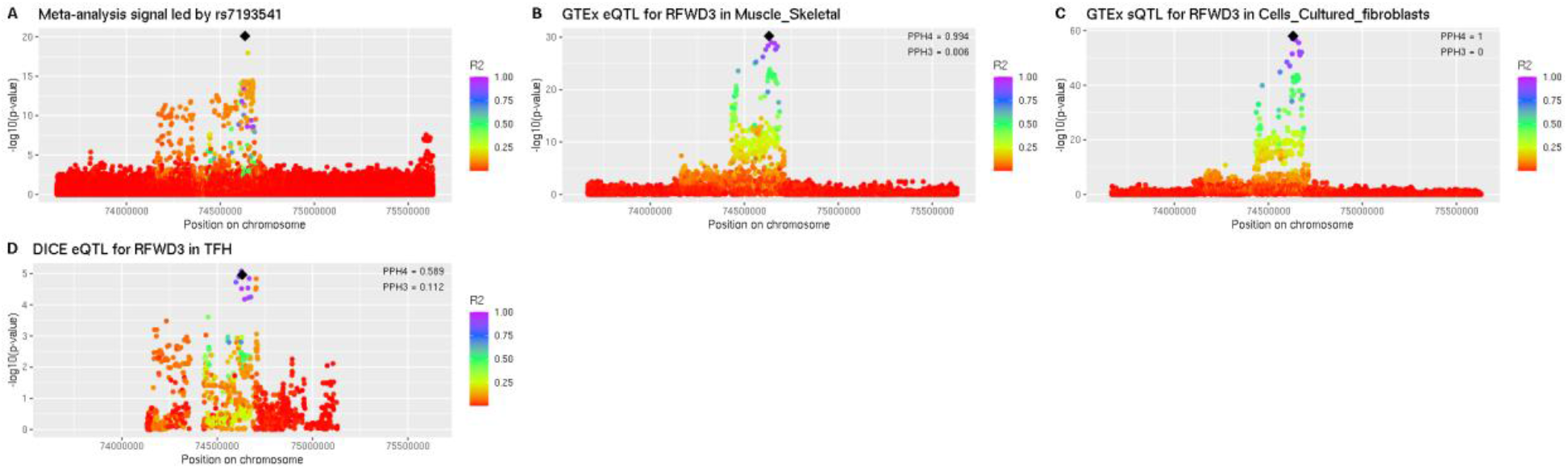

This meta-analysis signal best colocalized with *RFWD3* QTLs. *RFWD3* was also the proximal gene. We note that LeafCutter visualization of the splicing pattern supported an effect of the lead SNP at the association signal over different *RFWD3* splicing patterns as discussed in greater detail in the main text. Based on these results, we concluded that *RFWD3* was the best supported putative causal gene.

**rs6564996 (chr16:82173937:T:C)**

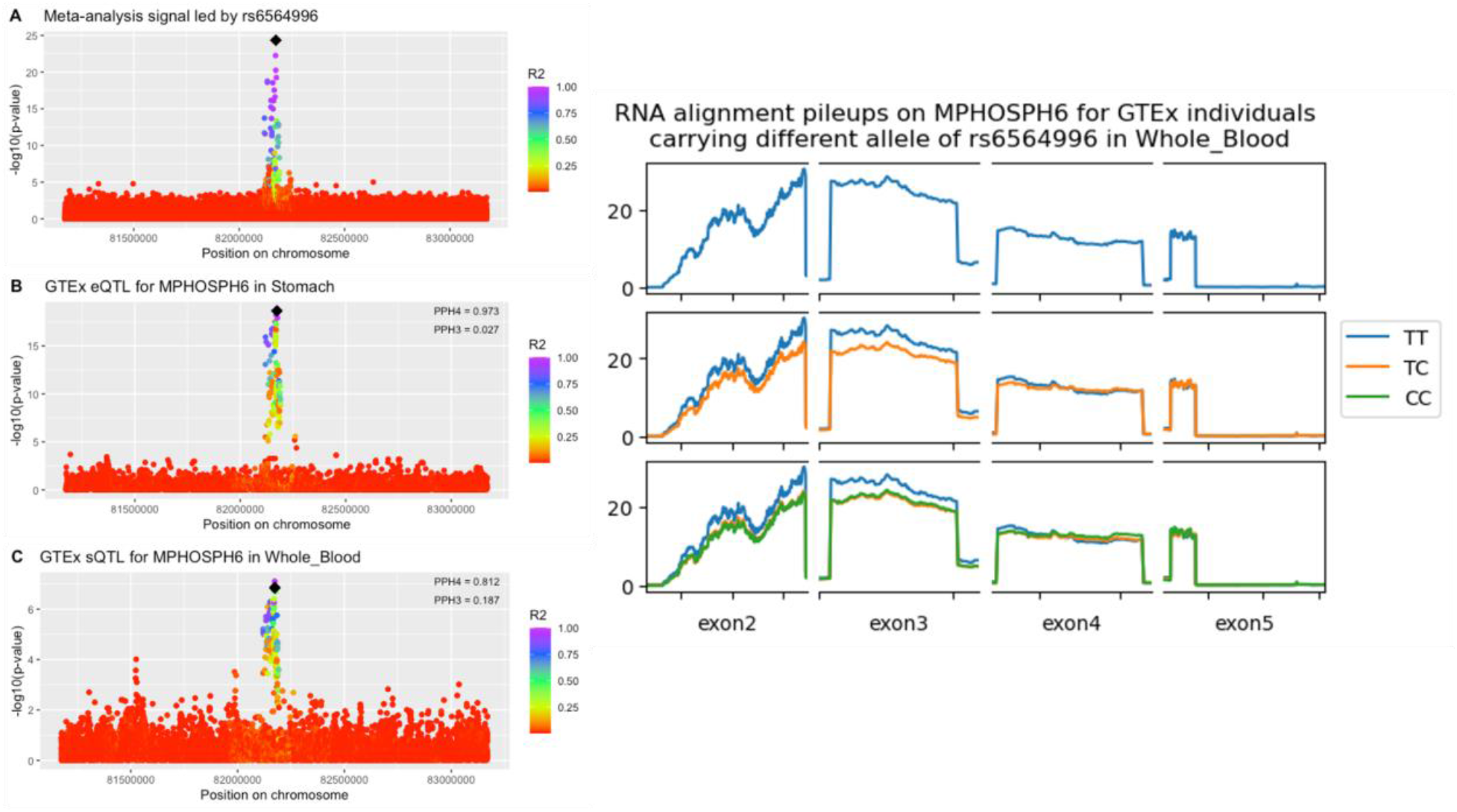

This meta-analysis signal best colocalized with *MPHOSPH6* QTLs. *MPHOSPH6* was also the proximal gene. The RNA pileup plot shows the aligned reads in the indicated GTEx tissue for the indicated exons that were included in the LeafCutter splicing cluster. Unlike an eQTL, a subset of exons show differences in the amount of reads aligned when stratified by the indicated genotype, supporting that this is a sQTL. Therefore, we concluded that *MPHOSPH6* was the best supported putative causal gene.

**rs59922886 (chr17:8236454:A:T)**

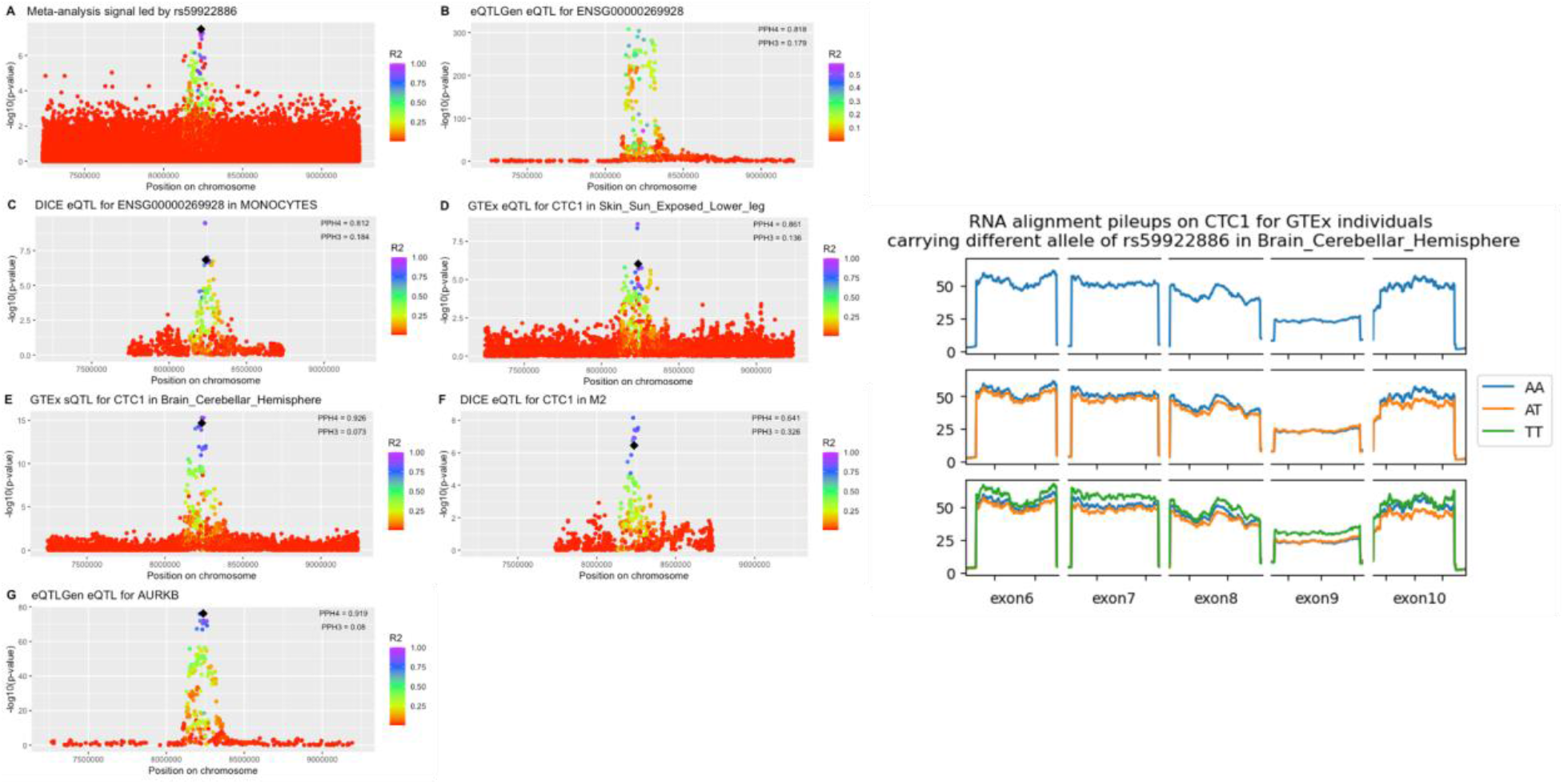

This meta-analysis signal strongly colocalized with QTLs for *CTC1* and *AURKB*. *CTC1* is the proximal gene and has known roles in telomere length regulation. The RNA pileup plot shows the aligned reads in the indicated GTEx tissue for the indicated exons that were included in the LeafCutter splicing cluster. Unlike an eQTL, a subset of exons show differences in the amount of reads aligned when stratified by the indicated genotype, supporting that this is a sQTL. Therefore, we concluded that *CTC1* was the best supported putative causal gene.

**rs144204502 (chr17:78187152:C:T)**

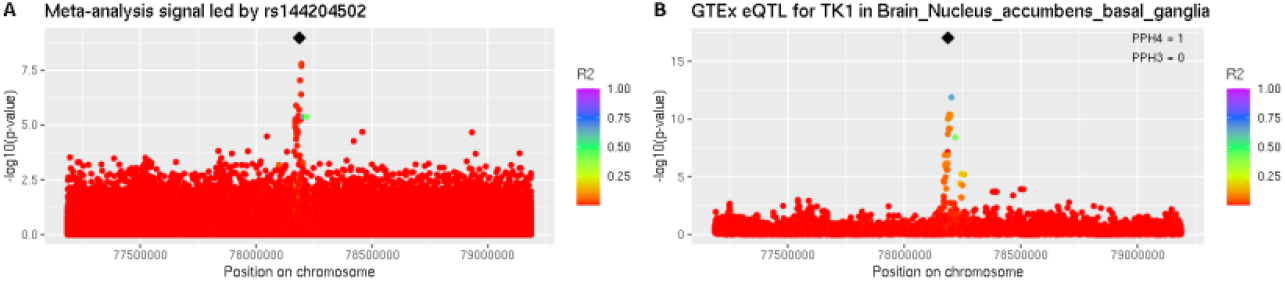

This meta-analysis signal best colocalized with *TK1* QTLs. *TK1* was also the proximal gene. Therefore, we concluded that *TK1* is the best supported putative casual gene.

**rs2124616 (chr18:661917:G:A)**

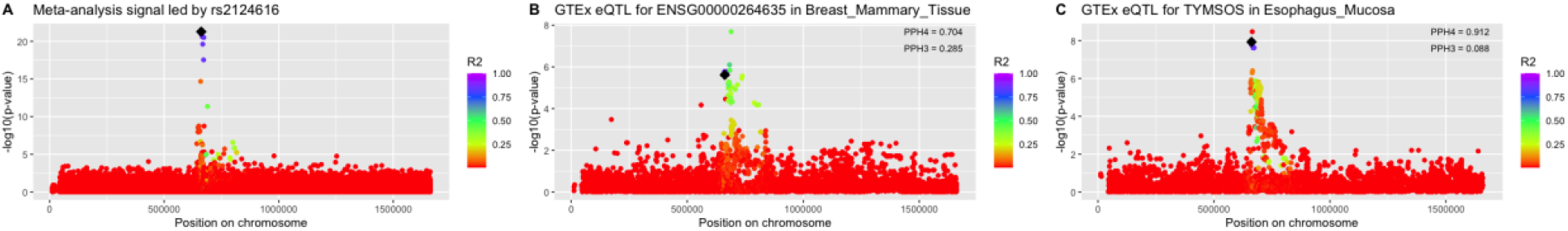

This meta-analysis signal best colocalized with a *TYMSOS* QTL. *TYMP* was the proximal gene. We concluded that *TYMSOS* was the best supported putative causal gene.

**rs8105767 (chr19:22032639:A:G)**

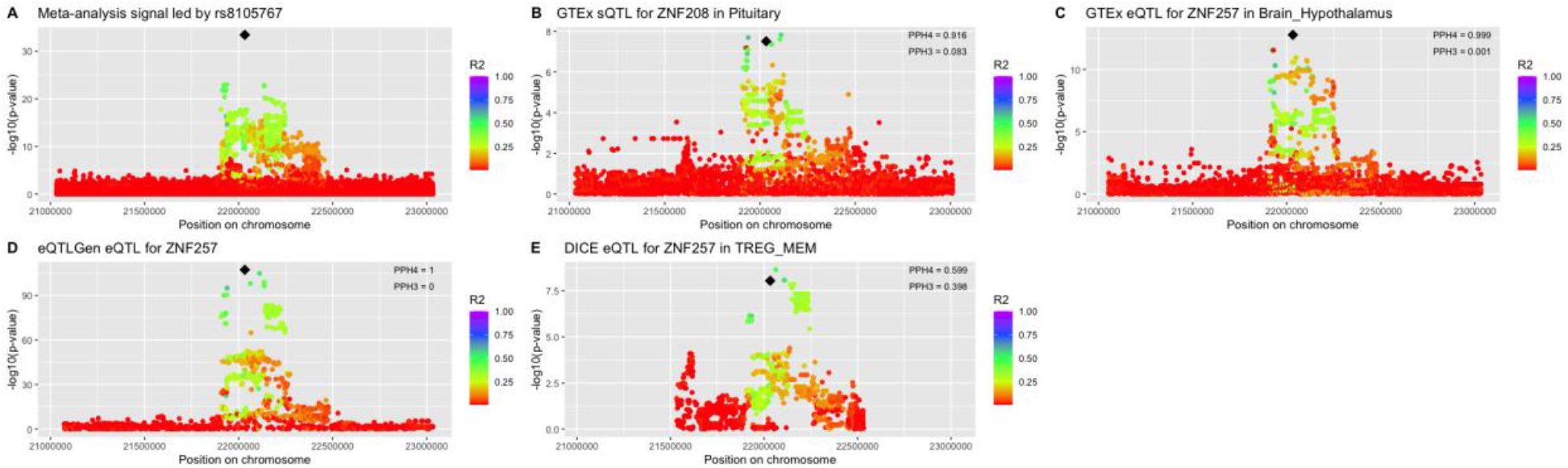

This meta-analysis signal colocalized with *ZNF257* and *ZNF208* QTLs. The colocalization with *ZNF257* QTLs was replicated in multiple datasets. *ZNF257* was also the proximal gene. Therefore, we concluded that *ZNF257* was the best supported putative causal gene.

**rs114703330 (chr20:63678039:T:C)**

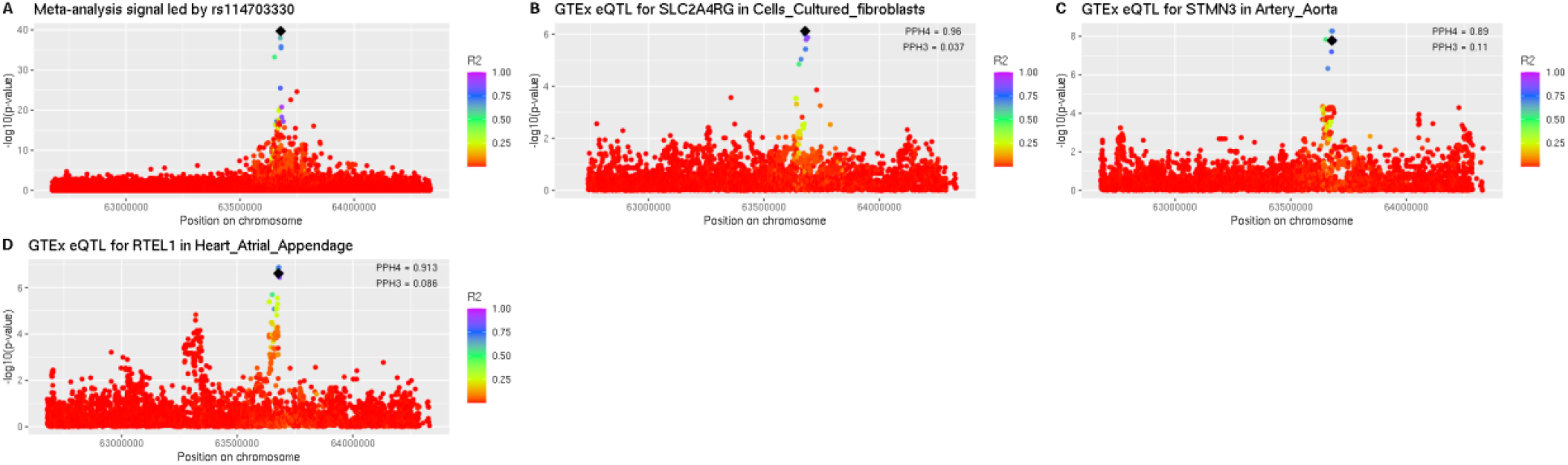

This meta-analysis signal best colocalized with QTLs for *RTEL1*, *SLC2A4RG*, and *STMN3*. The proximal gene was *RTEL1* and *RTEL1* has known roles in telomere length regulation. Therefore, we concluded that *RTEL1* was the best supported putative causal gene.

**rs28663120 (chr22:16973188:T:C)**

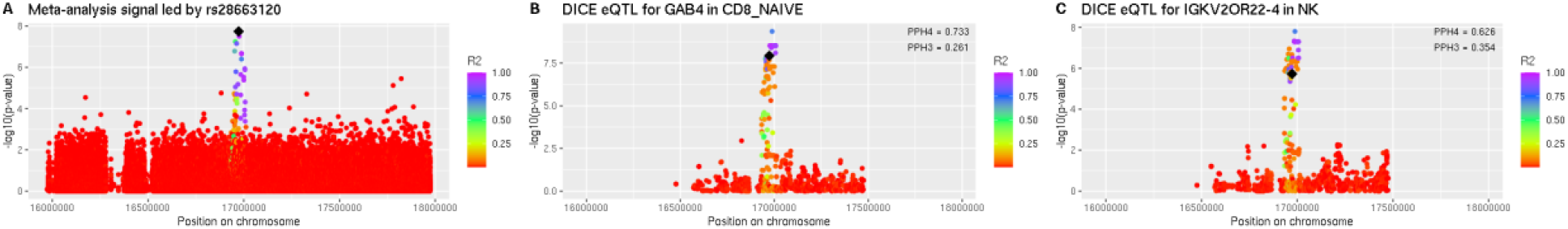

This meta-analysis signal colocalized with QTLs for *GAB4* and *IGKV2OR22-4*. *GAB4* was the proximal gene. Therefore, we concluded that *GAB4* is the best supported putative causal gene.

**rs131784 (chr22:50543007:G:A)**

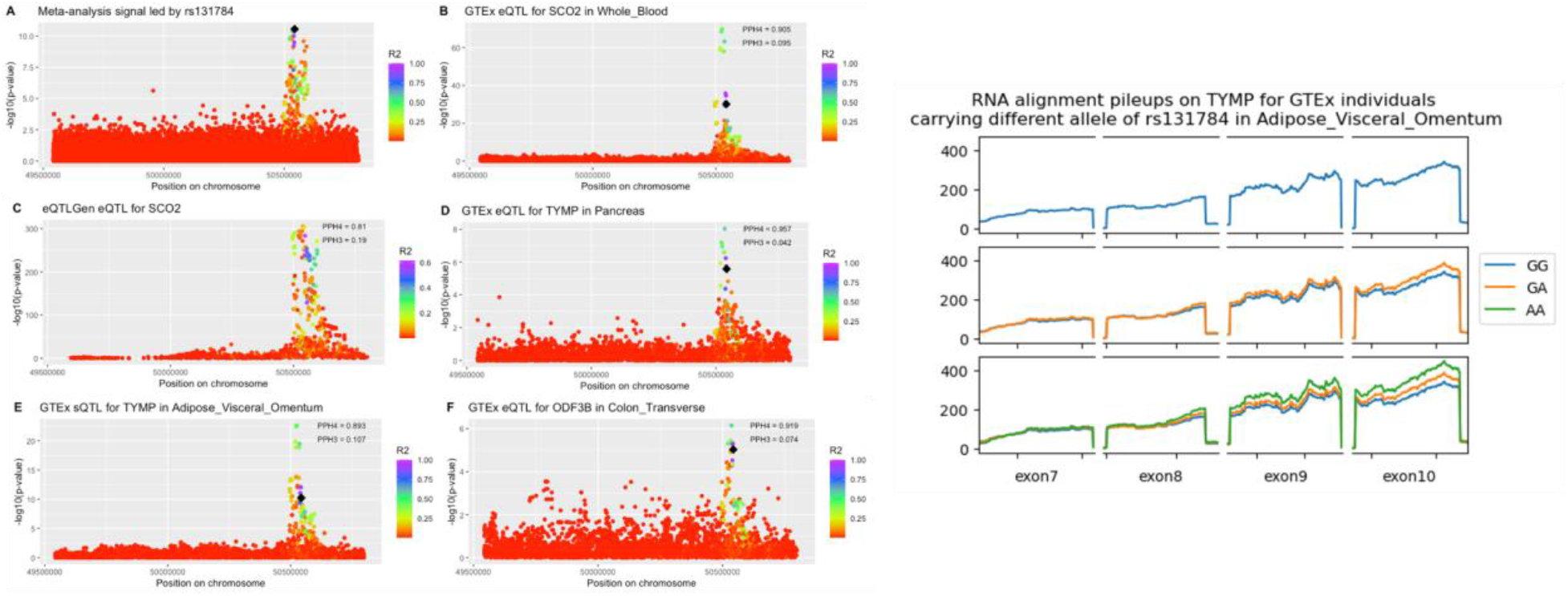

This meta-analysis signal colocalized best with QTLs for *TYMP*, *SCO2*, and *ODF3B*. The proximal gene was *KLHDC7B*. The RNA pileup plot shows the aligned reads in the indicated GTEx tissue for the indicated exons that were included in the LeafCutter splicing cluster. Unlike an eQTL, a subset of exons show differences in the amount of reads aligned when stratified by the indicated genotype, supporting that this is a sQTL. As colocalization was strongest with *TYMP* QTLs, we concluded that *TYMP* was the best supported putative causal gene.

**rs12394264 (chrX:66015290:G:A)**

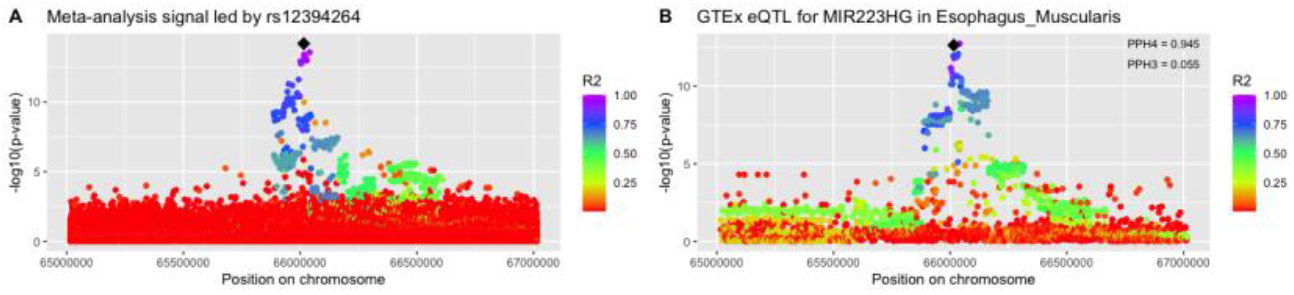

This meta-analysis signal best colocalized with QTLs for *MIR223HG*. *MIR223HG* was also the proximal gene. Therefore, we concluded that the best supported putative casual gene was *MIR223HG*.

## Supplemental Acknowledgements

### Generation of TOPMed whole genome sequencing data by study

Whole genome sequencing (WGS) for the Trans-Omics in Precision Medicine (TOPMed) program was supported by the National Heart, Lung and Blood Institute (NHLBI). WGS for NHLBI TOPMed: AFLMU (phs001543) was performed at Broad Genomics (3UM1HG008895- 01S2; HHSN268201500014C); WGS for NHLBI TOPMed: Amish (phs000956) was performed at Broad Genomics (3R01HL121007-01S1); WGS for NHLBI TOPMed: ARIC (phs001211) was performed at Baylor (3U54HG003273-12S2 / HHSN268201500015C,3R01HL092577-06S1), Broad Genomics (3U54HG003273-12S2 / HHSN268201500015C,3R01HL092577-06S1); WGS for NHLBI TOPMed: BioMe (phs001644) was performed at MGI (HHSN268201600037I,HHSN268201600033I,3UM1HG008853-01S2), Baylor (HHSN268201600037I,HHSN268201600033I,3UM1HG008853-01S2); WGS for NHLBI TOPMed: CAMP (phs001726) was performed at NWGC (HHSN268201600032I); WGS for NHLBI TOPMed: CARDIA (phs001612) was performed at Baylor (HHSN268201600033I); WGS for NHLBI TOPMed: CARE_BADGER (phs001728) was performed at NWGC (HHSN268201600032I); WGS for NHLBI TOPMed: CARE_CLIC (phs001729) was performed at NWGC (HHSN268201600032I); WGS for NHLBI TOPMed: CARE_PACT (phs001730) was performed at NWGC (HHSN268201600032I); WGS for NHLBI TOPMed: CARE_TREXA (phs001732) was performed at NWGC (HHSN268201600032I); WGS for NHLBI TOPMed: CFS (phs000954) was performed at NWGC (HHSN268201600032I,3R01HL098433-05S1); WGS for NHLBI TOPMed: ChildrensHS_GAP (phs001602) was performed at NWGC (HHSN268201600032I); WGS for NHLBI TOPMed: ChildrensHS_IGERA (phs001603) was performed at NWGC (HHSN268201600032I); WGS for NHLBI TOPMed: ChildrensHS_MetaAir (phs001604) was performed at NWGC (HHSN268201600032I); WGS for NHLBI TOPMed: CHIRAH (phs001605) was performed at NWGC (HHSN268201600032I); WGS for NHLBI TOPMed: CHS (phs001368) was performed at Baylor (HHSN268201600033I,3U54HG003273- 12S2 / HHSN268201500015C); WGS for NHLBI TOPMed: COPDGene (phs000951) was performed at NWGC (3R01HL089856-08S1,HHSN268201500014C), Broad Genomics (3R01HL089856-08S1,HHSN268201500014C); WGS for NHLBI TOPMed: CRA (phs000988) was performed at NWGC (3R37HL066289-13S1,HHSN268201600032I); WGS for NHLBI TOPMed: DHS (phs001412) was performed at Broad Genomics (HHSN268201500014C); WGS for NHLBI TOPMed: ECLIPSE (phs001472) was performed at MGI (HHSN268201600037I); WGS for NHLBI TOPMed: EOCOPD (phs000946) was performed at NWGC (3R01HL089856- 08S1); WGS for NHLBI TOPMed: FHS (phs000974) was performed at Broad Genomics (3U54HG003067- 12S2,3R01HL092577-06S1); WGS for NHLBI TOPMed: GALAI (phs001542) was performed at NWGC (HHSN268201600032I); WGS for NHLBI TOPMed: GALAII (phs000920) was performed at NYGC (3R01HL117004-02S3,HHSN268201600032I), NWGC (3R01HL117004-02S3,HHSN268201600032I), NYGC (UM1 HG008901); WGS for NHLBI TOPMed: GeneSTAR (phs001218) was performed at Psomagen (3R01HL112064- 04S1,R01HL112064,HHSN268201500014C), Illumina (3R01HL112064-04S1,R01HL112064,HHSN268201500014C), Broad Genomics (3R01HL112064-04S1,R01HL112064,HHSN268201500014C); WGS for NHLBI TOPMed: GENOA (phs001345) was performed at NWGC (3R01HL055673-18S1,HHSN268201500014C), Broad Genomics (3R01HL055673-18S1,HHSN268201500014C); WGS for NHLBI TOPMed: GenSalt (phs001217) was performed at Baylor (HHSN268201500015C); WGS for NHLBI TOPMed: GOLDN (phs001359) was performed at NWGC (3R01HL104135-04S1); WGS for NHLBI TOPMed: HCHS/SOL (phs001395) was performed at Baylor College of Medicine Human Genome Sequencing Center (HHSN268201600033I); WGS for NHLBI TOPMed: HVH (phs000993) was performed at Broad Genomics (3R01HL092577-06S1,3U54HG003273-12S2 / HHSN268201500015C), Baylor (3R01HL092577-06S1,3U54HG003273-12S2 / HHSN268201500015C); WGS for NHLBI TOPMed: HyperGEN (phs001293) was performed at NWGC (3R01HL055673-18S1); WGS for NHLBI TOPMed: IPF (phs001607) was performed at MGI (HHSN268201600037I); WGS for NHLBI TOPMed: JHS (phs000964) was performed at NWGC (HHSN268201100037C); WGS for NHLBI TOPMed: LTRC (phs001662) was performed at Broad Genomics (HHSN268201600034I); WGS for NHLBI TOPMed: Mayo_VTE (phs001402) was performed at Baylor (3U54HG003273-12S2 / HHSN268201500015C); WGS for NHLBI TOPMed: MESA (phs001416) was performed at Broad Genomics (3U54HG003067- 13S1,HHSN268201500014C); WGS for NHLBI TOPMed: MLOF (phs001515) was performed at Baylor (HHSN268201600033I,HHSN268201500016C), NYGC (HHSN268201600033I,HHSN268201500016C); WGS for NHLBI TOPMed: OMG_SCD (phs001608) was performed at Baylor (HHSN268201500015C); WGS for NHLBI TOPMed: PCGC_CHD (phs001735) was performed at Broad Genomics (HHSN268201600034I); WGS for NHLBI TOPMed: PharmHU (phs001466) was performed at Baylor (HHSN268201500015C); WGS for NHLBI TOPMed: PIMA (phs001727) was performed at NWGC (HHSN268201600032I); WGS for NHLBI TOPMed: PUSH_SCD (phs001682) was performed at Baylor (HHSN268201500015C); WGS for NHLBI TOPMed: REDS-III_Brazil (phs001468) was performed at Baylor (HHSN268201500015C); WGS for NHLBI TOPMed: SAFS (phs001215) was performed at Illumina (R01HL113322,3R01HL113323-03S1); WGS for NHLBI TOPMed: SAGE (phs000921) was performed at NYGC (3R01HL117004- 02S3,HHSN268201600032I), NWGC (3R01HL117004-02S3,HHSN268201600032I); WGS for NHLBI TOPMed: SAPPHIRE_asthma (phs001467) was performed at NWGC (HHSN268201600032I); WGS for NHLBI TOPMed: SARP (phs001446) was performed at NYGC (HHSN268201500016C); NHLBI TOPMed: SAS (phs000972) was performed at NWGC (HHSN268201100037C,HHSN268201500016C), NYGC (HHSN268201100037C,HHSN268201500016C); WGS for NHLBI TOPMed: THRV (phs001387) was performed at Baylor (3R01HL111249-04S1 / HHSN26820150015C); WGS for NHLBI TOPMed: VAFAR (phs000997) was performed at Broad Genomics (3U54HG003067-12S2 / 3U54HG003067- 13S1; 3UM1HG008895-01S2; 3UM1HG008895-01S2,3R01HL092577-06S1); WGS for NHLBI TOPMed: VU_AF (phs001032) was performed at Broad Genomics (3R01HL092577-06S1); WGS for NHLBI TOPMed: walk_PHaSST (phs001514) was performed at Baylor (HHSN268201500015C); WGS for NHLBI TOPMed: WGHS (phs001040) was performed at Broad Genomics (3R01HL092577-06S1); WGS for NHLBI TOPMed: WHI (phs001237) was performed at Broad Genomics (HHSN268201500014C). Core support including centralized genomic read mapping and genotype calling, along with variant quality metrics and filtering were provided by the TOPMed Informatics Research Center (3R01HL-117626-02S1; contract HHSN268201800002I). Core support including phenotype harmonization, data management, sample-identity QC, and general program coordination were provided by the TOPMed Data Coordinating Center (R01HL-120393; U01HL-120393; contract HHSN268201800001I). We gratefully acknowledge the studies and participants who provided biological samples and data for TOPMed. NYGC = New York Genome Center; Broad Genomics = Broad Institute Genomics Platform; NWGC = University of Washington Northwest Genomics Center; Illumina = Illumina Genomic Services; Psomagen = Psomagen Corp.; Baylor = Baylor Human Genome Sequencing Center; MGI = McDonnell Genome Institute

### Study-specific acknowledgements

#### NHLBI TOPMed: Atrial Fibrillation Biobank LMU (AFLMU) in the context of the ArrhythmiaBiobank-LMU

AFLMU is a repository of AF patients recruited in the context of the German Competence Network for Atrial Fibrillation (AFNET) and at the Department of Medicine I of the University Hospital Munich. In this context, DNA samples were preferentially sampled if the patient developed AF before the age of 60 years. Cases were selected if the diagnosis of atrial fibrillation was made on an electrocardiogram analyzed by a trained physician. Patients with signs of moderate to severe heart failure, moderate to severe valve disease or with hyperthyroidism were excluded from the study. All participants provided written informed consent. AFLMU was approved by the Ethics Committee at the Ludwig-Maximilian’s University.

#### NHLBI TOPMed: Genetics of Cardiometabolic Health in the Amish (Amish)

The Amish studies upon which these data are based were supported by NIH grants R01 AG18728, U01 HL072515, R01 HL088119, R01 HL121007, and P30 DK072488. See publication: PMID: 18440328

#### NHLBI TOPMed: Atherosclerosis Risk in Communities (ARIC)

The Atherosclerosis Risk in Communities study has been funded in whole or in part with Federal funds from the National Heart, Lung, and Blood Institute, National Institutes of Health, Department of Health and Human Services (contract numbers HHSN268201700001I, HHSN268201700002I, HHSN268201700003I, HHSN268201700004I and HHSN268201700005I). The authors thank the staff and participants of the ARIC study for their important contributions.

#### NHLBI TOPMed: The Genetics and Epidemiology of Asthma in Barbados (BAGS)

We gratefully acknowledge the contributions of Pissamai and Trevor Maul, Paul Levett, Anselm Hennis, P. Michele Lashley, Raana Naidu, Malcolm Howitt and Timothy Roach, and the numerous health care providers, and community clinics and co-investigators who assisted in the phenotyping and collection of DNA samples, and the families and patients for generously donating DNA samples to the Barbados Asthma Genetics Study (BAGS). The Genetics and Epidemiology of Asthma in Barbados is supported by National Institutes of Health (NIH) National Heart, Lung, Blood Institute TOPMed (R01 HL104608-S1) and: R01 AI20059, K23 HL076322, R01HL087699, and RC2 HL101651. For the specific cohort descriptions and descriptions regarding the collection of phenotype data can be found at: https://www.nhlbiwgs.org/group/bags-asthma. The authors wish to give special recognition to the individual study participants who provided biological samples and or data, without their support in research none of this would be possible.

#### NHLBI TOPMed: BioMe Biobank at Mount Sinai (BioMe)

The Mount Sinai BioMe Biobank has been supported by The Andrea and Charles Bronfman Philanthropies and in part by Federal funds from the NHLBI and NHGRI (U01HG00638001; U01HG007417; X01HL134588). We thank all participants in the Mount Sinai Biobank. We also thank all our recruiters who have assisted and continue to assist in data collection and management and are grateful for the computational resources and staff expertise provided by Scientific Computing at the Icahn School of Medicine at Mount Sinai.

#### NHLBI TOPMed: CAMP

We thank the clinical centers and the Data Coordinating Center of the Childhood Asthma Management Program (CAMP) as well as all of the study participants at the 8 clinical sites. The CAMP study was supported by NHLBI P01 HL132825.

#### NHLBI TOPMed: Coronary Artery Risk Development in Young Adults Study (CARDIA)

The Coronary Artery Risk Development in Young Adults Study (CARDIA) is conducted and supported by the National Heart, Lung, and Blood Institute (NHLBI) in collaboration with the University of Alabama at Birmingham (HHSN268201800005I & HHSN268201800007I), Northwestern University (HHSN268201800003I), University of Minnesota (HHSN268201800006I), and Kaiser Foundation Research Institute (HHSN268201800004I). CARDIA was also partially supported by the Intramural Research Program of the National Institute on Aging (NIA) and an intra-agency agreement between NIA and NHLBI (AG0005).

#### NHLBI TOPMed: CARE_BADGER

This research was supported by grants from the National Heart, Lung, and Blood Institute (NHLBI), ((5U10HL064287, 5U10HL064288, 5U10HL064295, 5U10HL064307, 5U10HL064305, 5U10HL064313, and HL080083)

#### NHLBI TOPMed: CARE_CLIC

This research was supported by grants from the National Heart, Lung, and Blood Institute (NHLBI), ((5U10HL064287, 5U10HL064288, 5U10HL064295, 5U10HL064307, 5U10HL064305, 5U10HL064313, and HL080083)

#### NHLBI TOPMed: CARE_PACT

This research was supported by grants from the National Heart, Lung, and Blood Institute (NHLBI), ((5U10HL064287, 5U10HL064288, 5U10HL064295, 5U10HL064307, 5U10HL064305, 5U10HL064313, and HL080083)

#### NHLBI TOPMed: CARE_TREXA

This research was supported by grants from the National Heart, Lung, and Blood Institute (NHLBI), ((5U10HL064287, 5U10HL064288, 5U10HL064295, 5U10HL064307, 5U10HL064305, 5U10HL064313, and HL080083)

#### NHLBI TOPMed: The Cleveland Family Study (CFS)

The Cleveland Family Study has been supported in part by National Institutes of Health grants [R01- HL046380, KL2-RR024990, R35-HL135818, and R01-HL113338].

#### NHLBI TOPMed: Children’s Health Study: Integrative Genetic Approaches to Gene-Air Pollution Interactions in Asthma (ChildrensHS_GAP)

The Integrative Genetic Approaches to Gene-Air Pollution Interactions in Asthma (GAP) study was supported by the National Institute of Environmental Health Sciences (NIEHS) grant # R01ES021801. The Children’s Health Study (CHS) was supported by the Southern California Environmental Health Sciences Center (grant P30ES007048); National Institute of Environmental Health Sciences (grants 5P01ES011627, ES021801, ES023262, P01ES009581, P01ES011627, P01ES022845, R01 ES016535, R03ES014046, P50 CA180905, R01HL061768, R01HL076647, R01HL087680 and RC2HL101651), the Environmental Protection Agency (grants RD83544101, R826708, RD831861, and R831845), and the Hastings Foundation.

#### NHLBI TOPMed: Children’s Health Study: Integrative Genomics and Environmental Research of Asthma (ChildrensHS_IGERA)

The Integrative Genomics and Environmental Research of Asthma (IGERA) Study was supported by the National Heart, Lung and Blood Institute (grant # RC2HL101543-The Asthma BioRepository for Integrative Genomics Research, PI Gilliland/Raby). The Children’s Health Study (CHS) was supported by the Southern California Environmental Health Sciences Center (grant P30ES007048); National Institute of Environmental Health Sciences (grants 5P01ES011627, ES021801, ES023262, P01ES009581, P01ES011627, P01ES022845, R01 ES016535, R03ES014046, P50 CA180905, R01HL061768, R01HL076647, R01HL087680 and RC2HL101651), the Environmental Protection Agency (grants RD83544101, R826708, RD831861, and R831845), and the Hastings Foundation.

#### NHLBI TOPMed: Children’s Health Study: Effects of Air Pollution on the Development of Obesity in Children (ChildrensHS_MetaAir)

The Effects of Air Pollution on the Development of Obesity in Children (Meta-AIR) study was supported by the Southern California Children’s Environmental Health Center funded by the National Institute of Environmental Health Sciences (NIEHS) (P01ES022845) and the Environmental Protection Agency (EPA) (RD-83544101–0). The Children’s Health Study (CHS) was supported by the Southern California Environmental Health Sciences Center (grant P30ES007048); National Institute of Environmental Health Sciences (grants 5P01ES011627, ES021801, ES023262, P01ES009581, P01ES011627, P01ES022845, R01 ES016535, R03ES014046, P50 CA180905, R01HL061768, R01HL076647, R01HL087680 and RC2HL101651), the Environmental Protection Agency (grants RD83544101, R826708, RD831861, and R831845), and the Hastings Foundation.

#### NHLBI TOPMed: Genetics Sub-Study of Chicago Initiative to Raise Asthma Health Equity (CHIRAH)

Support for the Genetics Sub-Study of Chicago Initiative to Raise Asthma Health Equity was provided by NHLBI grant number UO1 HL072496.

#### NHLBI TOPMed: Cardiovascular Health Study (CHS)

This research was supported by contracts HHSN268201200036C, HHSN268200800007C, HHSN268201800001C, N01HC55222, N01HC85079, N01HC85080, N01HC85081, N01HC85082, N01HC85083, N01HC85086, and 75N92021D00006, and grants U01HL080295 and U01HL130114 from the National Heart, Lung, and Blood Institute (NHLBI), with additional contribution from the National Institute of Neurological Disorders and Stroke (NINDS). Additional support was provided by R01AG023629 from the National Institute on Aging (NIA). A full list of principal CHS investigators and institutions can be found at CHS-NHLBI.org. The content is solely the responsibility of the authors and does not necessarily represent the official views of the National Institutes of Health.

#### NHLBI TOPMed: Genetic Epidemiology of COPD (COPDGene) in the TOPMed Program

The COPDGene project described was supported by Award Number U01 HL089897 and Award Number U01 HL089856 from the National Heart, Lung, and Blood Institute. The content is solely the responsibility of the authors and does not necessarily represent the official views of the National Heart, Lung, and Blood Institute or the National Institutes of Health. The COPDGene project is also supported by the COPD Foundation through contributions made to an Industry Advisory Board comprised of AstraZeneca, Boehringer Ingelheim, GlaxoSmithKline, Novartis, Pfizer, Siemens and Sunovion. A full listing of COPDGene investigators can be found at: http://www.copdgene.org/directory

#### NHLBI TOPMed: The Genetic Epidemiology of Asthma in Costa Rica (CRA)

This study was supported by NHLBI grants R37 HL066289 and P01 HL132825. We wish to acknowledge the investigators at the Channing Division of Network Medicine at Brigham and Women’s Hospital, the investigators at the Hospital Nacional de Niños in San José, Costa Rica and the study subjects and their extended family members who contributed samples and genotypes to the study, and the NIH/NHLBI for its support in making this project possible.

#### NHLBI TOPMed: Diabetes Heart Study (DHS)

This work was supported by R01 HL92301, R01 HL67348, R01 NS058700, R01 AR48797, R01 DK071891, R01 AG058921, the General Clinical Research Center of the Wake Forest University School of Medicine (M01 RR07122, F32 HL085989), the American Diabetes Association, and a pilot grant from the Claude Pepper Older Americans Independence Center of Wake Forest University Health Sciences (P60 AG10484).

#### NHLBI TOPMed: ECLIPSE

The ECLIPSE study (NCT00292552) was sponsored by GlaxoSmithKline. The ECLIPSE investigators included: ECLIPSE Investigators — Bulgaria: Y. Ivanov, Pleven; K. Kostov, Sofia. Canada: J. Bourbeau, Montreal; M. Fitzgerald, Vancouver, BC; P. Hernandez, Halifax, NS; K. Killian, Hamilton, ON; R. Levy, Vancouver, BC; F. Maltais, Montreal; D. O’Donnell, Kingston, ON. Czech Republic: J. Krepelka, Prague. Denmark: J. Vestbo, Hvidovre. The Netherlands: E. Wouters, Horn-Maastricht. New Zealand: D. Quinn, Wellington. Norway: P. Bakke, Bergen. Slovenia: M. Kosnik, Golnik. Spain: A. Agusti, J. Sauleda, P. de Mallorca. Ukraine: Y. Feschenko, V. Gavrisyuk, L. Yashina, Kiev; N. Monogarova, Donetsk. United Kingdom: P. Calverley, Liverpool; D. Lomas, Cambridge; W. MacNee, Edinburgh; D. Singh, Manchester; J. Wedzicha, London. United States: A. Anzueto, San Antonio, TX; S. Braman, Providence, RI; R. Casaburi, Torrance CA; B. Celli, Boston; G. Giessel, Richmond, VA; M. Gotfried, Phoenix, AZ; G. Greenwald, Rancho Mirage, CA; N. Hanania, Houston; D. Mahler, Lebanon, NH; B. Make, Denver; S. Rennard, Omaha, NE; C. Rochester, New Haven, CT; P. Scanlon, Rochester, MN; D. Schuller, Omaha, NE; F. Sciurba, Pittsburgh; A. Sharafkhaneh, Houston; T. Siler, St. Charles, MO; E. Silverman, Boston; A. Wanner, Miami; R. Wise, Baltimore; R. ZuWallack, Hartford, CT. ECLIPSE Steering Committee: H. Coxson (Canada), C. Crim (GlaxoSmithKline, USA), L. Edwards (GlaxoSmithKline, USA), D. Lomas (UK), W. MacNee (UK), E. Silverman (USA), R. Tal-Singer (Co-chair, GlaxoSmithKline, USA), J. Vestbo (Co-chair, Denmark), J. Yates (GlaxoSmithKline, USA). ECLIPSE Scientific Committee: A. Agusti (Spain), P. Calverley (UK), B. Celli (USA), C. Crim (GlaxoSmithKline, USA), B. Miller (GlaxoSmithKline, USA), W. MacNee (Chair, UK), S. Rennard (USA), R. Tal-Singer (GlaxoSmithKline, USA), E. Wouters (The Netherlands), J. Yates (GlaxoSmithKline, USA).

#### NHLBI TOPMed: Boston Early-Onset COPD Study in the TOPMed Program (EOCOPD)

The Boston Early-Onset COPD Study was supported by R01 HL113264 and U01 HL089856 from the National Heart, Lung, and Blood Institute.

#### NHLBI TOPMed: Whole Genome Sequencing and Related Phenotypes in the Framingham Heart Study (FHS)

The Framingham Heart Study (FHS) acknowledges the support of contracts NO1-HC-25195, HHSN268201500001I, and 75N92019D00031 from the National Heart, Lung and Blood Institute and grant supplement R01 HL092577-06S1 for this research. We also acknowledge the dedication of the FHS study participants without whom this research would not be possible.

#### NHLBI TOPMed: Genes-environments and Admixture in Latino Asthmatics (GALA I) Study

The Genes-environments and Admixture in Latino Americans (GALA I) Study was supported by the National Heart, Lung, and Blood Institute of the National Institute of Health (NIH) grants R01HL117004 and X01HL134589; study enrollment supported by Sandler Center for Basic Research in Asthma and the Sandler Family Foundation, the American Asthma Foundation, the American Lung Association, the NIH grants K23HL04464 and HL07185, the Resource Centers for Minority Aging Research from the National Institute on Aging, RCMAR P30-AG15272, the National Institute of Nursing Research and the National Center on Minority Health and Health Disparities.

#### NHLBI TOPMed: Genes-environments and Admixture in Latino Asthmatics (GALA II) Study

The Genes-environments and Admixture in Latino Americans (GALA II) Study was supported by the National Heart, Lung, and Blood Institute of the National Institute of Health (NIH) grants R01HL117004 and X01HL134589; study enrollment supported by the Sandler Family Foundation, the American Asthma Foundation, the RWJF Amos Medical Faculty Development Program, Harry Wm. and Diana V. Hind Distinguished Professor in Pharmaceutical Sciences II and the National Institute of Environmental Health Sciences grant R01ES015794. WGS of part of GALA II was performed by New York Genome Center under The Centers for Common Disease Genomics of the Genome Sequencing Program (GSP) Grant (UM1 HG008901). The GSP Coordinating Center (U24 HG008956) contributed to cross-program scientific initiatives and provided logistical and general study coordination. GSP is funded by the National Human Genome Research Institute, the National Heart, Lung, and Blood Institute, and the National Eye Institute. The GALA II study collaborators include Shannon Thyne, UCSF; Harold J. Farber, Texas Children’s Hospital; Denise Serebrisky, Jacobi Medical Center; Rajesh Kumar, Lurie Children’s Hospital of Chicago; Emerita Brigino-Buenaventura, Kaiser Permanente; Michael A. LeNoir, Bay Area Pediatrics; Kelley Meade, UCSF Benioff Children’s Hospital, Oakland; William Rodriguez-Cintron, VA Hospital, Puerto Rico; Pedro C. Avila, Northwestern University; Jose R. Rodriguez-Santana, Centro de Neumologia Pediatrica; Luisa N. Borrell, City University of New York; Adam Davis, UCSF Benioff Children’s Hospital, Oakland; Saunak Sen, University of Tennessee and Fred Lurmann, Sonoma Technologies, Inc. The authors acknowledge the families and patients for their participation and thank the numerous health care providers and community clinics for their support and participation in GALA II. In particular, the authors thank study coordinator Sandra Salazar; the recruiters who obtained the data: Duanny Alva, MD, Gaby Ayala-Rodriguez, Lisa Caine, Elizabeth Castellanos, Jaime Colon, Denise DeJesus, Blanca Lopez, Brenda Lopez, MD, Louis Martos, Vivian Medina, Juana Olivo, Mario Peralta, Esther Pomares, MD, Jihan Quraishi, Johanna Rodriguez, Shahdad Saeedi, Dean Soto, Ana Taveras; and the lab researcher Celeste Eng who processed the biospecimens.

#### NHLBI TOPMed: GeneSTAR (Genetic Study of Atherosclerosis Risk)

The Johns Hopkins Genetic Study of Atherosclerosis Risk (GeneSTAR) was supported by grants from the National Institutes of Health through the National Heart, Lung, and Blood Institute (U01HL72518, HL087698, HL112064) and by a grant from the National Center for Research Resources (M01- RR000052) to the Johns Hopkins General Clinical Research Center. We would like to thank the participants and families of GeneSTAR and our dedicated staff for all their sacrifices.

#### NHLBI TOPMed: Genetic Epidemiology Network of Arteriopathy (GENOA)

Support for GENOA was provided by the National Heart, Lung and Blood Institute (HL054457, HL054464, HL054481, HL119443, and HL087660) of the National Institutes of Health.

#### NHLBI TOPMed: Genetic Epidemiology Network of Salt Sensitivity (GenSalt)

The Genetic Epidemiology Network of Salt-Sensitivity (GenSalt) was supported by research grants (U01HL072507, R01HL087263, and R01HL090682) from the National Heart, Lung and Blood Institute, National Institutes of Health, Bethesda, MD.

#### NHLBI TOPMed: Genetics of Lipid Lowering Drugs and Diet Network (GOLDN)

GOLDN biospecimens, baseline phenotype data, and intervention phenotype data were collected with funding from National Heart, Lung and Blood Institute (NHLBI) grant U01 HL072524. Whole-genome sequencing in GOLDN was funded by NHLBI grant R01 HL104135 and supplement R01 HL104135- 04S1.

#### NHLBI TOPMed: Hispanic Community Health Study/Study of Latinos (HCHS_SOL)

The Hispanic Community Health Study/Study of Latinos is a collaborative study supported by contracts from the National Heart, Lung, and Blood Institute (NHLBI) to the University of North Carolina (HHSN268201300001I / N01-HC-65233), University of Miami (HHSN268201300004I / N01-HC65234), Albert Einstein College of Medicine (HHSN268201300002I / N01-HC-65235), University of Illinois at Chicago – HHSN268201300003I / N01-HC-65236 Northwestern Univ), and San Diego State University (HHSN268201300005I / N01-HC-65237). The following Institutes/Centers/Offices have contributed to the HCHS/SOL through a transfer of funds to the NHLBI: National Institute on Minority Health and Health Disparities, National Institute on Deafness and Other Communication Disorders, National Institute of Dental and Craniofacial Research, National Institute of Diabetes and Digestive and Kidney Diseases, National Institute of Neurological Disorders and Stroke, NIH Institution-Office of Dietary Supplements.

#### NHLBI TOPMed: Heart and Vascular Health Study (HVH)

The Heart and Vascular Health Study was supported by grants HL068986, HL085251, HL095080, and HL073410 from the National Heart, Lung, and Blood Institute.

#### NHLBI TOPMed: Hypertension Genetic Epidemiology Network (HyperGEN)

The HyperGEN Study is part of the National Heart, Lung, and Blood Institute (NHLBI) Family Blood Pressure Program; collection of the data represented here was supported by grants U01 HL054472 (MN Lab), U01 HL054473 (DCC), U01 HL054495 (AL FC), and U01 HL054509 (NC FC). The HyperGEN: Genetics of Left Ventricular Hypertrophy Study was supported by NHLBI grant R01 HL055673 with whole-genome sequencing made possible by supplement-18S1.

#### NHLBI TOPMed: IPF

This research was supported by the National Heart, Lung and Blood Institute (R01-HL097163, P01- HL092870, and UH3-HL123442) and the Department of Defense (W81XWH-17-1-0597).

#### NHLBI TOPMed: The Jackson Heart Study (JHS)

The Jackson Heart Study (JHS) is supported and conducted in collaboration with Jackson State University (HHSN268201800013I), Tougaloo College (HHSN268201800014I), the Mississippi State Department of Health (HHSN268201800015I) and the University of Mississippi Medical Center (HHSN268201800010I, HHSN268201800011I and HHSN268201800012I) contracts from the National Heart, Lung, and Blood Institute (NHLBI) and the National Institute on Minority Health and Health Disparities (NIMHD). The authors also wish to thank the staffs and participants of the JHS.

#### NHLBI TOPMed: LTRC

This study utilized biological specimens and data provided by the Lung Tissue Research Consortium (LTRC) supported by the National Heart, Lung, and Blood Institute (NHLBI). The LTRC was sponsored by a contract from the

#### NHLBI: HHSN2682016000021 NHLBI TOPMed: Mayo Clinic Venous Thromboembolism Study (Mayo_VTE)

Funded, in part, by grants from the National Institutes of Health, National Heart, Lung and Blood Institute (HL66216 and HL83141). the National Human Genome Research Institute (HG04735, HG06379), and research support provided by Mayo Foundation.

#### NHLBI TOPMed: Multi-Ethnic Study of Atherosclerosis (MESA)

Whole genome sequencing (WGS) for the Trans-Omics in Precision Medicine (TOPMed) program was supported by the National Heart, Lung and Blood Institute (NHLBI). WGS for “NHLBI TOPMed: Multi-Ethnic Study of Atherosclerosis (MESA)” (phs001416.v3.p1) was performed at the Broad Institute of MIT and Harvard (3U54HG003067-13S1). Centralized read mapping and genotype calling, along with variant quality metrics and filtering were provided by the TOPMed Informatics Research Center (3R01HL-117626-02S1). Phenotype harmonization, data management, sample-identity QC, and general study coordination, were provided by the TOPMed Data Coordinating Center (3R01HL-120393-02S1), and TOPMed MESA Multi-Omics (HHSN2682015000031/HSN26800004). The MESA projects are conducted and supported by the National Heart, Lung, and Blood Institute (NHLBI) in collaboration with MESA investigators. Support for the Multi-Ethnic Study of Atherosclerosis (MESA) projects are conducted and supported by the National Heart, Lung, and Blood Institute (NHLBI) in collaboration with MESA investigators. Support for MESA is provided by contracts 75N92020D00001, HHSN268201500003I, N01-HC-95159, 75N92020D00005, N01-HC-95160, 75N92020D00002, N01-HC-95161, 75N92020D00003, N01-HC-95162, 75N92020D00006, N01-HC-95163, 75N92020D00004, N01-HC-95164, 75N92020D00007, N01-HC-95165, N01-HC-95166, N01- HC-95167, N01-HC-95168, N01-HC-95169, UL1-TR-000040, UL1-TR-001079, UL1-TR-001420, UL1TR001881, DK063491, and R01HL105756. The authors thank the other investigators, the staff, and the participants of the MESA study for their valuable contributions. A full list of participating MESA investigators and institutes can be found at http://www.mesa-nhlbi.org.

#### NHLBI TOPMed: My Life, Our Future (MLOF)

The My Life, Our Future samples and data are made possible through the partnership of Bloodworks Northwest, the American Thrombosis and Hemostasis Network, the National Hemophilia Foundation, and Bioverativ. We gratefully acknowledge the hemophilia treatment centers and their patients who provided biological samples and phenotypic data.

#### NHLBI TOPMed: Outcome Modifying Genes in Sickle Cell Disease (OMG-SCD)

The OMG-SCD study was administered by Marilyn J. Telen, M.D. and Allison E. Ashley-Koch, Ph.D. from Duke University Medical Center, and collection of the data set was supported by grants HL068959 and HL079915 from the National Heart, Lung, and Blood Institute (NHLBI) of the National Institute of Health (NIH).

#### NHLBI TOPMed: Pediatric Cardiac Genomics Consortium’s Congenital Heart Disease Biobank (PCGC-CHD)

The Pediatric Cardiac Genomics Consortium (PCGC) program is funded by the National Heart, Lung, and Blood Institute, National Institutes of Health, U.S. Department of Health and Human Services through grants UM1HL128711, UM1HL098162, UM1HL098147, UM1HL098123, UM1HL128761, and U01HL131003.

#### NHLBI TOPMed: The Pharmacogenomics of Hydroxyurea in Sickle Cell Disease (PharmHU)

Collection of the PharmHU samples and data were supported in part by the Department of Pediatrics, Baylor College of Medicine funds, National Institutes of Health (NIH) National Institute of Diabetes and Digestive and Kidney Diseases (NIDDK) grant 1K08 DK110448-01, NIH NHLBI R01 HL069234, and U01-HL117721 funded by NHLBI. We are very grateful to the patients with sickle cell disease for their participation in PharmHU.

#### NHLBI TOPMed: PIMA

This research was supported by grants from the National Heart, Lung, and Blood Institute (NHLBI), ((5U10HL064287, 5U10HL064288, 5U10HL064295, 5U10HL064307, 5U10HL064305, 5U10HL064313, and HL080083)

#### NHLBI TOPMed: PUSH_SCD

We thank Dr. Victor R Gordeuk and the investigators of the PUSH study and the patients who participated in the study. We also thank the PUSH clinical site team: Howard University: Victor R Gordeuk, Sergei Nekhai, Oswaldo Castro, Sohail Rana, Mehdi Nouraie, James G Taylor 6th, Children National Medical Center: Caterina Minniti, Deepika Darbari, Lori Lutchman-Jones, Nitti Dham, Craig Sable, NHLBI: Mark Gladwin, Greg Kato. University of Michigan: Andrew Campbell, Gregory Ensing, Manuel Arteta, Special thanks to the volunteers who participated in the PUSH study. This project was funded with federal funds from the NHLBI, NIH. Detail description of the study was published in Haematologica. 2009 Mar;94(3):340-7, MInniti C, et al. “Elevated tricuspid regurgitant jet velocity in children and adolescents with sickle cell disease: association with hemolysis and hemoglobin oxygen desaturation.”

#### NHLBI TOPMed: Recipient Epidemiology and Donor Evaluation Study-III (REDS-III_Brazil)

The Recipient Epidemiology and Donor Evaluation Study (REDS)-III was funded by NIH NHLBI contract HHSN268201100007I and conducted under the leadership of Simone Glynn (NHLBI), and principle investigators Brian Custer and Ester Sabino. We are grateful to the Brazilian sickle cell disease patients who participated in the REDS-III study and provided blood samples for whole genome sequencing as well as the REDS-III staff: Vitalant Research Institute (Shannon Kelly), University of Sao Paulo (Miriam V Flor Park, Ligia Capuani), Hemominas Belo Horizonte (Anna Barbara Proietti), Hemominas Montes Claros (Rosimere Alfonso), Hemominas Juiz de Fora (Daniela de O. Werneck Rodrigues), Hemope (Paula Loureiro), Hemorio (Claudia Maximo).

#### NHLBI TOPMed: San Antonio Family Heart Study (SAFS)

Collection of the San Antonio Family Study data was supported in part by National Institutes of Health (NIH) grants P01 HL045522, R01 MH078143, R01 MH078111 and R01 MH083824; and whole genome sequencing of SAFS subjects was supported by U01 DK085524 and R01 HL113323. We are very grateful to the participants of the San Antonio Family Study for their continued involvement in our research programs.

#### NHLBI TOPMed: Study of African Americans, Asthma, Genes and Environment (SAGE)

The Study of African Americans, Asthma, Genes and Environments (SAGE) was supported by by the National Heart, Lung, and Blood Institute of the National Institute of Health (NIH) grants R01HL117004 and X01HL134589; study enrollment supported by the Sandler Family Foundation, the American Asthma Foundation, the RWJF Amos Medical Faculty Development Program, Harry Wm. and Diana V. Hind Distinguished Professor in Pharmaceutical Sciences II. The SAGE study collaborators include Harold J. Farber, Texas Children’s Hospital; Emerita Brigino-Buenaventura, Kaiser Permanente; Michael A. LeNoir, Bay Area Pediatrics; Kelley Meade, UCSF Benioff Children’s Hospital, Oakland; Luisa N. Borrell, City University of New York; Adam Davis, UCSF Benioff Children’s Hospital, Oakland and Fred Lurmann, Sonoma Technologies, Inc. The authors acknowledge the families and patients for their participation and thank the numerous health care providers and community clinics for their support and participation in SAGE. In particular, the authors thank study coordinator Sandra Salazar; the recruiters who obtained the data: Lisa Caine, Elizabeth Castellanos, Brenda Lopez, MD, Shahdad Saeedi; and the lab researcher Celeste Eng who processed the biospecimens.

#### NHLBI TOPMed: Study of Asthma Phenotypes & Pharmacogenomic Interactions by RaceEthnicity (SAPPHIRE_asthma)

The SAPPHIRE cohort was supported by grant funding from the Fund for Henry Ford Hospital, the American Asthma Foundation, and the following institutes of the National Institutes of Health: the National Heart Lung and Blood Institute (R01HL141845, R01HL118267, X01HL134589, R01HL079055), the National Institute of Allergy and Infectious Diseases (R01AI079139, R01AI061774), and the National Institute of Diabetes and Digestive and Kidney Diseases (R01DK113003, R01DK064695).

#### NHLBI TOPMed: Genetics of Sarcoidosis in African Americans (Sarcoidosis)

Supported by the National Institutes of Health under Grant R01HL113326-05, P30 GM110766- 01, and U54GM104938-06.

#### NHLBI TOPMed: Severe Asthma Research Program (SARP)

The authors acknowledge the contributions of the study coordinators and staff at each of the clinical centers and the Data Coordinating Center as well as all the study participants that have been integral to the success of the NHLBI Severe Asthma Research Program (funded by U10 HL109164, U10 HL109257, U10 HL109146, U10 HL109172, U10 HL109250, U10 HL109250, U10 HL109250, U10 HL109168, U10 HL109152, U10 HL109086).

#### NHLBI TOPMed: Genome-wide Association Study of Adiposity in Samoans (SAS)

Financial support from the U.S. National Institutes of Health Grants R01-HL093093 and R01HL133040. We acknowledge the assistance of the Samoa Ministry of Health and the Samoa Bureau of Statistics for their guidance and support in the conduct of this study. We thank the local village officials for their help and the participants for their generosity. The following publication describes the origin of the dataset: Hawley NL, Minster RL, Weeks DE, Viali S, Reupena MS, Sun G, Cheng H, Deka R, McGarvey ST. Prevalence of Adiposity and Associated Cardiometabolic Risk Factors in the Samoan Genome-Wide Association Study. Am J Human Biol 2014. 26: 491-501. DOI: 10.1002/jhb.22553. PMID: 24799123.

#### NHLBI TOPMed: Rare Variants for Hypertension in Taiwan Chinese (THRV)

The Rare Variants for Hypertension in Taiwan Chinese (THRV) is supported by the National Heart, Lung, and Blood Institute (NHLBI) grant (R01HL111249) and its participation in TOPMed is supported by an NHLBI supplement (R01HL111249-04S1). THRV is a collaborative study between Washington University in St. Louis, LA BioMed at Harbor UCLA, University of Texas in Houston, Taichung Veterans General Hospital, Taipei Veterans General Hospital, Tri-Service General Hospital, National Health Research Institutes, National Taiwan University, and Baylor University. THRV is based (substantially) on the parent SAPPHIRe study, along with additional population-based and hospital-based cohorts. SAPPHIRe was supported by NHLBI grants (U01HL54527, U01HL54498) and Taiwan funds, and the other cohorts were supported by Taiwan funds.

#### NHLBI TOPMed: The Vanderbilt AF Ablation Registry (VAFAR)

The research reported in this article was supported by grants from the American Heart Association to Dr. Shoemaker (11CRP742009), Dr. Darbar (EIA 0940116N), and grants from the National Institutes of Health (NIH) to Dr. Darbar (R01 HL092217), and Dr. Roden (U19 HL65962, and UL1 RR024975). The project was also supported by a CTSA award (UL1 TR00045) from the National Center for Advancing Translational Sciences. Its contents are solely the responsibility of the authors and do not necessarily represent the official views of the National Center for Advancing Translational Sciences or the NIH.

#### NHLBI TOPMed: The Vanderbilt Atrial Fibrillation Registry (VU_AF)

The research reported in this article was supported by grants from the American Heart Association to Dr. Darbar (EIA 0940116N), and grants from the National Institutes of Health (NIH) to Dr. Darbar (HL092217), and Dr. Roden (U19 HL65962, and UL1 RR024975). This project was also supported by CTSA award (UL1TR000445) from the National Center for Advancing Translational Sciences. Its contents are solely the responsibility of the authors and do not necessarily represent the official views of the National Center for Advancing Translational Sciences of the NIH.

#### NHLBI TOPMed: Treatment of Pulmonary Hypertension and Sickle Cell Disease With Sildenafil Therapy (Walk-PHaSST)

We thank Dr. Mark Gladwin and the investigators of the Walk-PHasst study and the patients who participated in the study. We also thanks the walk-PHaSST clinical site team: Albert Einstein College of Medicine: Jane Little and Verlene Davis; Columbia University: Robyn Barst, Erika Rosenzweig, Margaret Lee and Daniela Brady; UCSF Benioff Children’s Hospital Oakland: Claudia Morris, Ward Hagar, Lisa Lavrisha, Howard Rosenfeld, and Elliott Vichinsky; Children’s Hospital of Pittsburgh of UPMC: Regina McCollum; Hammersmith Hospital, London: Sally Davies, Gaia Mahalingam, Sharon Meehan, Ofelia Lebanto, and Ines Cabrita; Howard University: Victor Gordeuk, Oswaldo Castro, Onyinye Onyekwere,, Alvin Thomas, Gladys Onojobi, Sharmin Diaz, Margaret Fadojutimi-Akinsiku, and Randa Aladdin; Johns Hopkins University: Reda Girgis, Sophie Lanzkron and Durrant Barasa; NHLBI: Mark Gladwin, Greg Kato, James Taylor, Vandana Sachdev, Wynona Coles, Catherine Seamon, Mary Hall, Amy Chi, Cynthia Brenneman, Wen Li, and Erin Smith; University of Colorado: Kathryn Hassell, David Badesch, Deb McCollister and Julie McAfee; University of Illinois at Chicago: Dean Schraufnagel, Robert Molokie, George Kondos, Patricia Cole-Saffold, and Lani Krauz; National Heart & Lung Institute, Imperial College London: Simon Gibbs. Thanks also to the data coordination center team from Rho, Inc.: Nancy Yovetich, Rob Woolson, Jamie Spencer, Christopher Woods, Karen Kesler, Vickie Coble, and Ronald W. Helms. We also thank Dr. Yingze Zhang for directing the Walk-PHasst repository and Dr. Mehdi Nouraie for maintaining the Walk-PHasst database and Dr. Jonathan Goldsmith as a NIH program director for this study. Special thanks to the volunteers who participated in the Walk-PHaSST study. This project was funded with federal funds from the NHLBI, NIH, Department of Health and Human Services, under contract HHSN268200617182C. This study is registered at www.clinicaltrials.gov as NCT00492531. Detail description of the study was published in Blood, 2011 118:855-864, Machado et al “Hospitalization for pain in patients with sickle cell disease treated with sildenafil for elevated TRV and low exercise capacity”.

#### NHLBI TOPMed: Novel Risk Factors for the Development of Atrial Fibrillation in Women (WGHS)

The WGHS is supported by the National Heart, Lung, and Blood Institute (HL043851 and HL080467) and the National Cancer Institute (CA047988 and UM1CA182913). The most recent cardiovascular endpoints were supported by ARRA funding HL099355.

#### NHLBI TOPMed: Women’s Health Initiative (WHI)

The WHI program is funded by the National Heart, Lung, and Blood Institute, National Institutes of Health, U.S. Department of Health and Human Services through contracts HHSN268201600018C, HHSN268201600001C, HHSN268201600002C, HHSN268201600003C, and HHSN268201600004C. This manuscript was prepared in collaboration with investigators of the WHI, and has been reviewed and/or approved by the Women’s Health Initiative (WHI). The short list of WHI investigators can be found at https://www.whi.org/researchers/Documents%20%20Write%20a%20Paper/WHI%20Investigator%20Short%20List.pdf.

### Other funding acknowledgements

Rebecca Keener was supported in part by NIH/NIGMS grant 5K12GM123914. This work was carried out at the Advanced Research Computing at Hopkins (ARCH) core facility (rockfish.jhu.edu), which is supported by the National Science Foundation (NSF) grant number OAC1920103. Matthew P. Conomos was supported by R01HL-120393; contract HHSN268201800001I; U01 HL137162. Brian E. Cade was supported by R01HL153805. Barry I. Freedman was supported by R01 DK071891. Lifang Hou was supported by R01AG081244, R01AG069120. Marilyn J. Telen was supported by NHLBI (R01HL68959, R01HL87681 and R01HL079915) for the collection of the samples. Allison E. Ashley-Koch was supported by NHLBI (R01HL68959, R01HL87681 and R01HL079915) for the collection of the samples. Joshua C. Bis was supported by R01HL105756. Zhanghua Chen was supported by P30ES007048, 5P01ES011627, ES021801, ES023262, P01ES009581, P01ES011627, P01ES022845, R01 ES016535, R03ES014046, P50 CA180905, R01HL061768, R01HL076647, R01HL087680, RC2HL101651, RD83544101, R826708, RD831861, R831845, R00ES027870, and the Hastings Foundation. Dr. Patrick T. Ellinor was supported by grants from the National Institutes of Health (1RO1HL092577, 1R01HL157635, 5R01HL139731), from the American Heart Association Strategically Focused Research Networks (18SFRN34110082), and from the European Union (MAESTRIA 965286). Myriam Fornage was supported by U01AG052409, U01AG058589. Bruce D. Gelb was supported by U01HL153009. Frank D. Gilliland was supported by P30ES007048, 5P01ES011627, ES021801, ES023262, P01ES009581, P01ES011627, P01ES022845, R01 ES016535, R03ES014046, P50 CA180905, R01HL061768, R01HL076647, R01HL087680, RC2HL101651, RD83544101, R826708, RD831861, R831845, R00ES027870, and the Hastings Foundation. Talat Islam was supported by P30ES007048, 5P01ES011627, ES021801, ES023262, P01ES009581, P01ES011627, P01ES022845, R01 ES016535, R03ES014046, P50 CA180905, R01HL061768, R01HL076647, R01HL087680, RC2HL101651, RD83544101, R826708, RD831861, R831845, R00ES027870, and the Hastings Foundation. Rajesh Kumar was supported by UM1AI160040, U01AI160018-01, UG1HL139125, R01AI153239. Ruth J.F. Looks was supported by R01DK107786; R01HG010297. Nicholette D. Palmer was supported by R01 AG058921. Susan Redline was supported by R35HL135818. David Schwartz was supported by P01-HL162607, R01- HL158668, R01-HL149836, IO1BX005295, UG3/UH3-HL151865, and X01-HL134585. The Johns Hopkins Genetic Study of Atherosclerosis Risk (GeneSTAR) was supported by grants from the National Institutes of Health through the National Heart, Lung, and Blood Institute (U01HL72518, HL087698, HL112064) and by a grant from the National Center for Research Resources (M01-RR000052) to the Johns Hopkins General Clinical Research Center. Dr. Rasika Mathias receives support as the Sarah Miller Coulson Scholar in the Johns Hopkins Center for Innovative Medicine. Carol W. Greider was supported by NIH R35 CA209974. Alexis Battle was supported by NIH/NIGMS R35GM139580.

### TOPMed Consortium members

Namiko Abe, Gonçalo Abecasis, Francois Aguet, Christine Albert, Laura Almasy, Alvaro Alonso, Seth Ament, Peter Anderson, Pramod Anugu, Deborah Applebaum-Bowden, Kristin Ardlie, Dan Arking, Donna K Arnett, Allison Ashley-Koch, Stella Aslibekyan, Tim Assimes, Paul Auer, Dimitrios Avramopoulos, Najib Ayas, Adithya Balasubramanian, John Barnard, Kathleen Barnes, R. Graham Barr, Emily Barron-Casella, Lucas Barwick, Terri Beaty, Gerald Beck, Diane Becker, Lewis Becker, Rebecca Beer, Amber Beitelshees, Emelia Benjamin, Takis Benos, Marcos Bezerra, Larry Bielak, Joshua Bis, Thomas Blackwell, John Blangero, Nathan Blue, Eric Boerwinkle, Donald W. Bowden, Russell Bowler, Jennifer Brody, Ulrich Broeckel, Jai Broome, Deborah Brown, Karen Bunting, Esteban Burchard, Carlos Bustamante, Erin Buth, Brian Cade, Jonathan Cardwell, Vincent Carey, Julie Carrier, April P. Carson, Cara Carty, Richard Casaburi, Juan P Casas Romero, James Casella, Peter Castaldi, Mark Chaffin, Christy Chang, Yi-Cheng Chang, Daniel Chasman, Sameer Chavan, Bo-Juen Chen, Wei-Min Chen, Yii-Der Ida Chen, Michael Cho, Seung Hoan Choi, Lee-Ming Chuang, Mina Chung, Ren-Hua Chung, Clary Clish, Suzy Comhair, Matthew Conomos, Elaine Cornell, Adolfo Correa, Carolyn Crandall, James Crapo, L. Adrienne Cupples, Joanne Curran, Jeffrey Curtis, Brian Custer, Coleen Damcott, Dawood Darbar, Sean David, Colleen Davis, Michelle Daya, Mariza de Andrade, Lisa de las Fuentes, Paul de Vries, Michael DeBaun, Ranjan Deka, Dawn DeMeo, Scott Devine, Huyen Dinh, Harsha Doddapaneni, Qing Duan, Shannon Dugan-Perez, Ravi Duggirala, Jon Peter Durda, Susan K. Dutcher, Charles Eaton, Lynette Ekunwe, Adel El Boueiz, Patrick Ellinor, Leslie Emery, Serpil Erzurum, Charles Farber, Jesse Farek, Tasha Fingerlin, Matthew Flickinger, Myriam Fornage, Nora Franceschini, Chris Frazar, Mao Fu, Stephanie M. Fullerton, Lucinda Fulton, Stacey Gabriel, Weiniu Gan, Shanshan Gao, Yan Gao, Margery Gass, Heather Geiger, Bruce Gelb, Mark Geraci, Soren Germer, Robert Gerszten, Auyon Ghosh, Richard Gibbs, Chris Gignoux, Mark Gladwin, David Glahn, Stephanie Gogarten, Da-Wei Gong, Harald Goring, Sharon Graw, Kathryn J. Gray, Daniel Grine, Colin Gross, C. Charles Gu, Yue Guan, Xiuqing Guo, Namrata Gupta, Jeff Haessler, Michael Hall, Yi Han, Patrick Hanly, Daniel Harris, Nicola L. Hawley, Jiang He, Ben Heavner, Susan Heckbert, Ryan Hernandez, David Herrington, Craig Hersh, Bertha Hidalgo, James Hixson, Brian Hobbs, John Hokanson, Elliott Hong, Karin Hoth, Chao (Agnes) Hsiung, Jianhong Hu, Yi-Jen Hung, Haley Huston, Chii Min Hwu, Marguerite Ryan Irvin, Rebecca Jackson, Deepti Jain, Cashell Jaquish, Jill Johnsen, Andrew Johnson, Craig Johnson, Rich Johnston, Kimberly Jones, Hyun Min Kang, Robert Kaplan, Sharon Kardia, Shannon Kelly, Eimear Kenny, Michael Kessler, Alyna Khan, Ziad Khan, Wonji Kim, John Kimoff, Greg Kinney, Barbara Konkle, Charles Kooperberg, Holly Kramer, Christoph Lange, Ethan Lange, Leslie Lange, Cathy Laurie, Cecelia Laurie, Meryl LeBoff, Jiwon Lee, Sandra Lee, Wen-Jane Lee, Jonathon LeFaive, David Levine, Daniel Levy, Joshua Lewis, Xiaohui Li, Yun Li, Henry Lin, Honghuang Lin, Xihong Lin, Simin Liu, Yongmei Liu, Yu Liu, Ruth J.F. Loos, Steven Lubitz, Kathryn Lunetta, James Luo, Ulysses Magalang, Michael Mahaney, Barry Make, Ani Manichaikul, Alisa Manning, JoAnn Manson, Lisa Martin, Melissa Marton, Susan Mathai, Rasika Mathias, Susanne May, Patrick McArdle, Merry-Lynn McDonald, Sean McFarland, Stephen McGarvey, Daniel McGoldrick, Caitlin McHugh, Becky McNeil, Hao Mei, James Meigs, Vipin Menon, Luisa Mestroni, Ginger Metcalf, Deborah A Meyers, Emmanuel Mignot, Julie Mikulla, Nancy Min, Mollie Minear, Ryan L Minster, Braxton D. Mitchell, Matt Moll, Zeineen Momin, May E. Montasser, Courtney Montgomery, Donna Muzny, Josyf C Mychaleckyj, Girish Nadkarni, Rakhi Naik, Take Naseri, Pradeep Natarajan, Sergei Nekhai, Sarah C. Nelson, Bonnie Neltner, Caitlin Nessner, Deborah Nickerson, Osuji Nkechinyere, Kari North, Jeff O’Connell, Tim O’Connor, Heather Ochs-Balcom, Geoffrey Okwuonu, Allan Pack, David T. Paik, Nicholette Palmer, James Pankow, George Papanicolaou, Cora Parker, Gina Peloso, Juan Manuel Peralta, Marco Perez, James Perry, Ulrike Peters, Patricia Peyser, Lawrence S Phillips, Jacob Pleiness, Toni Pollin, Wendy Post, Julia Powers Becker, Meher Preethi Boorgula, Michael Preuss, Bruce Psaty, Pankaj Qasba, Dandi Qiao, Zhaohui Qin, Nicholas Rafaels, Laura Raffield, Mahitha Rajendran, Vasan S. Ramachandran, D.C. Rao, Laura Rasmussen-Torvik, Aakrosh Ratan, Susan Redline, Robert Reed, Catherine Reeves, Elizabeth Regan, Alex Reiner, Muagututi‘a Sefuiva Reupena, Ken Rice, Stephen Rich, Rebecca Robillard, Nicolas Robin, Dan Roden, Carolina Roselli, Jerome Rotter, Ingo Ruczinski, Alexi Runnels, Pamela Russell, Sarah Ruuska, Kathleen Ryan, Ester Cerdeira Sabino, Danish Saleheen, Shabnam Salimi, Sejal Salvi, Steven Salzberg, Kevin Sandow, Vijay G. Sankaran, Jireh Santibanez, Karen Schwander, David Schwartz, Frank Sciurba, Christine Seidman, Jonathan Seidman, Frédéric Sériès, Vivien Sheehan, Stephanie L. Sherman, Amol Shetty, Aniket Shetty, Wayne Hui-Heng Sheu, M. Benjamin Shoemaker, Brian Silver, Edwin Silverman, Robert Skomro, Albert Vernon Smith, Jennifer Smith, Josh Smith, Nicholas Smith, Tanja Smith, Sylvia Smoller, Beverly Snively, Michael Snyder, Tamar Sofer, Nona Sotoodehnia, Adrienne M. Stilp, Garrett Storm, Elizabeth Streeten, Jessica Lasky Su, Yun Ju Sung, Jody Sylvia, Adam Szpiro, Daniel Taliun, Hua Tang, Margaret Taub, Kent D. Taylor, Matthew Taylor, Simeon Taylor, Marilyn Telen, Timothy A. Thornton, Machiko Threlkeld, Lesley Tinker, David Tirschwell, Sarah Tishkoff, Hemant Tiwari, Catherine Tong, Russell Tracy, Michael Tsai, Dhananjay Vaidya, David Van Den Berg, Peter VandeHaar, Scott Vrieze, Tarik Walker, Robert Wallace, Avram Walts, Fei Fei Wang, Heming Wang, Jiongming Wang, Karol Watson, Jennifer Watt, Daniel E. Weeks, Joshua Weinstock, Bruce Weir, Scott T Weiss, Lu-Chen Weng, Jennifer Wessel, Cristen Willer, Kayleen Williams, L. Keoki Williams, Scott Williams, Carla Wilson, James Wilson, Lara Winterkorn, Quenna Wong, Baojun Wu, Joseph Wu, Huichun Xu, Lisa Yanek, Ivana Yang, Ketian Yu, Seyedeh Maryam Zekavat, Yingze Zhang, Snow Xueyan Zhao, Wei Zhao, Xiaofeng Zhu, Elad Ziv, Michael Zody, Sebastian Zoellner

### Hematology and Hemostasis working group members

Laura Almasy, Kurtis Anthony, Dan Arking, Allison Ashley-Koch, Paul Auer, Abraham Aviv, Andrea Baccarelli, Emily Barron-Casella, Lewis Becker, Romit Bhattacharya, Alexander Bick, Larry Bielak, Thomas Blackwell, John Blangero, Kelly Bolton, Jennifer Brody, Derek Brown, Deepika Burkardt, James Casella, Liam Cato, Christy Chang, Nilanjan Chatterjee, Han Chen, Ming-Huei Chen, Michael Cho, Zeynep Coban Akdemir, Jason Collins, Karen Conneely, Matthew Conomos, Paul de Vries, Dawn DeMeo, Pinkal Desai, Qing Duan, Connor Emdin, Nauder Faraday, Annette Fitzpatrick, Travis Fleming, James Floyd, Santhi Ganesh, Brady Gaynor, LaShaunta Glover, Jacob Graham, Edward Ha, Nadia Hansel, Manjit Hanspal, Ross Hardison, Ben Heavner, Julian Hecker, Scott Heemann, Craig Hersh, Chani Hodonsky, Michael Honigberg, Steve Horvath, Yao Hu, Jennifer Huffman, Carmen Isasi, Kruthika Raman Iyer, Sidd Jaiswal, Cashell Jaquish, Jin Jin, Jill Johnsen, Andrew Johnson, Brian Joyce, Joel Kaufman, Rebecca Keener, Shannon Kelly, Alyna Khan, Sumeet Khetarpal, Greg Kinney, Malgorzata Klauzinska, Barbara Konkle, Charles Kooperberg, Mohanraj Krishnan, Ethan Lange, Leslie Lange, Cathy Laurie, Brandon Lê, Grace Lee, Claire Leiser, Guillaume Lettre, Dan Levy, Joshua Lewis, Bingshan Li, Yun Li, L. A. Liggett, Amarise Little, Shelly-Ann Love, Megan Lynch, Mitchell Machiela, Rasika Mathias, Ravi Mathur, Karen Miga, Anna Mikhaylova, Julie Mikulla, Braxton D. Mitchell, Alanna C Morrison, Rakhi Naik, Drew Nannini, Vivek Naranbhai, Pradeep Natarajan, Jeff O’Connell, Christopher O’Donnell, Nels Olson, Helena Palma Gudiel, Nathan Pankratz, Benedict Paten, James Perry, James Pirruccello, Linda Polfus, Diddier Prada, Bruce Psaty, Laura Raffield, Elizabeth Regan, Alex Reiner, Stephen Rich, Shabnam Salimi, Vijay G. Sankaran, Noah Simon, Nicholas Smith, James Stewart, Adrienne M. Stilp, Shakira Suglia, Weihong Tang, Hua Tang, Margaret Taub, Kent D. Taylor, Marilyn Telen, Florian Thibord, Timothy A. Thornton, Russell Tracy, Md Mesbah Uddin, Heming Wang, Lachelle Weeks, Joshua Weinstock, Ellen Werner, Marsha Wheeler, Eric Whitsel, Kerri L. Wiggins, Lisa Yanek, Yu-Chung Yang, Kimberley Youkhana, Michael Young, Anthony Zannas, Seyedeh Maryam Zekavat, Wei Zhao, Yinan Zheng, Ying Zhou

### Structural Variation working group members

Paul Auer, Kathleen Barnes, Thomas Blackwell, Harrison Brand, Ulrich Broeckel, Deepika Burkardt, Mark Chaisson, Kei Hang Katie Chan, Seung Hoan Choi, Zechen Chong, Bradley Coe, John Cole, Ryan Collins, Matthew Conomos, Michelle Daya, Scott Devine, Evan Eichler, Annette Fitzpatrick, C. Charles Gu, Amelia Weber Hall, Ira Hall, Bob Handsaker, Ben Heavner, Scott Heemann, James Hixson, Jicai Jiang, Jill Johnsen, Michelle Jones, Brian Joyce, Goo Jun, Hyun Min Kang, Spencer Kelley, Charles Kooperberg, John Lane, Cathy Laurie, Seung-been, Steven Lee, Dan Levy, Yang Li, Honghuang Lin, Simin Liu, Angel CY Mak, Alisa Manning, Rasika Mathias, Steve McCarroll, Julie Mikulla, Jean Monlong, Drew Nannini, Giuseppe Narzisi, Jeff O’Connell, Wanda O’Neal, Grier Page, Nathan Pankratz, Benedict Paten, Alexandre Pereira, Patricia Peyser, Nathan Pezant, Gloria Quach, Aakrosh Ratan, Alex Reiner, Stephen Rich, Ingo Ruczinski, Aniko Sabo, Steven Salzberg, Jonathan Seidman, Minseok Seo, Yichen Si, Nasa Sinnott Armstrong, Albert Vernon Smith, Vinodh Srinivasasainagendra, Arvis Sulovari, Margaret Taub, Joshua Weinstock, Marsha Wheeler, James Wilson, Huichun Xu, Wei Zhao, Xuefang Zhao, Yinan Zheng, Degui Zhi, Sebastian Zoellner

---

**Supplementary Figure 1:**
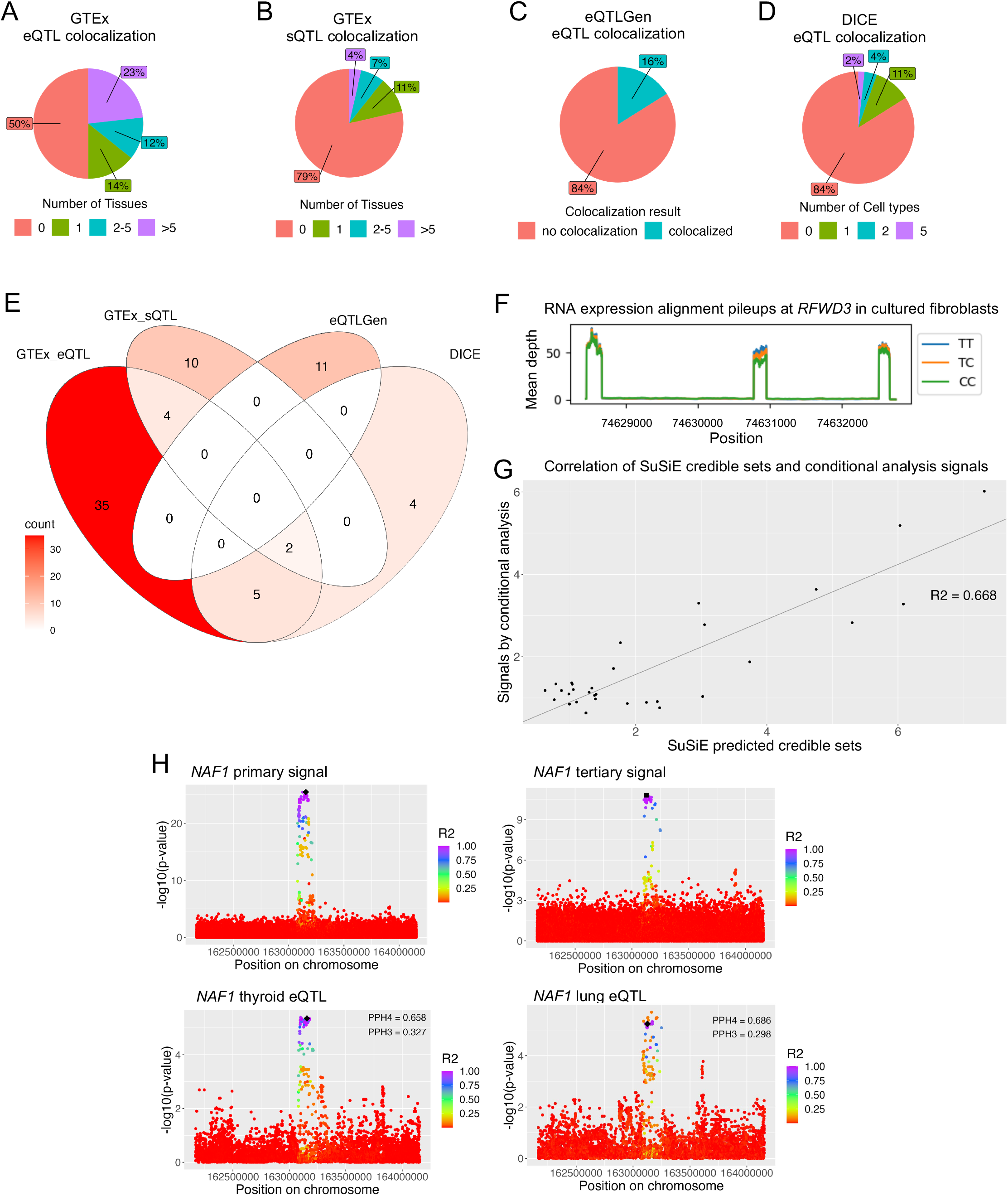
Fine-mapping analyses nominate putative causal variants and genes affecting telomere length. A-B. Percent of meta-analysis signals that colocalize (PPH4 > 0.7) with a GTEx cis-eQTL or cis-sQTL for any gene across differing numbers of tissues. C. Percent of meta-analysis signals that colocalize (PPH4 > 0.5) with a DICE eQTL for any gene in any cell type. The threshold for PPH4 was reduced because the DICE dataset has lower power to detect eQTLs since the dataset is derived from 91 individuals. A-C. In some instances one signal may colocalize with one gene in tissue/cell type X while colocalizing with a second gene in tissue/cell type Y; this case would be reported as number of tissue/cell type = 2. D. Percent of meta-analysis signals that colocalize (PPH4 > 0.7) with an eQTLGen cis-eQTL. eQTLGen cis-eQTLs are derived from whole blood only. E. Venn diagram showing in which datasets meta-analysis signals colocalized with the same gene quantitative trait locus (QTL) in any cell type across datasets. F. RNA expression pileup plots from GTEx v8 for *RFWD3* in cultured fibroblasts. The plot is stratified by genotype for the sentinel SNP at the meta-analysis locus. G. Correlation of the number of SuSiE predicted credible sets and the number of signals by conditional analysis (Taub et al. 2022). H. *NAF1* primary signal from the TOPMed pooled GWAS analysis colored by r^2^ with the lead SNP (black diamond). The best colocalization result for this signal was the *NAF1* eQTL in thyroid. After two rounds of conditional analysis on the lead SNP and secondary signal lead SNP at the *NAF1* locus, a tertiary signal remained significant (Taub et al. 2022). The tertiary signal and *NAF1* lung eQTL are colored by r^2^ with the lead SNP at the tertiary signal (black square). The best colocalization result for the tertiary GWAS signal was with the *NAF1* eQTL in lung. The *NAF1* eQTL in thyroid did not colocalize with the *NAF1* eQTL in lung (PPH3 = 0.721, PPH4 = 0.217).

**Supplementary Figure 2:**
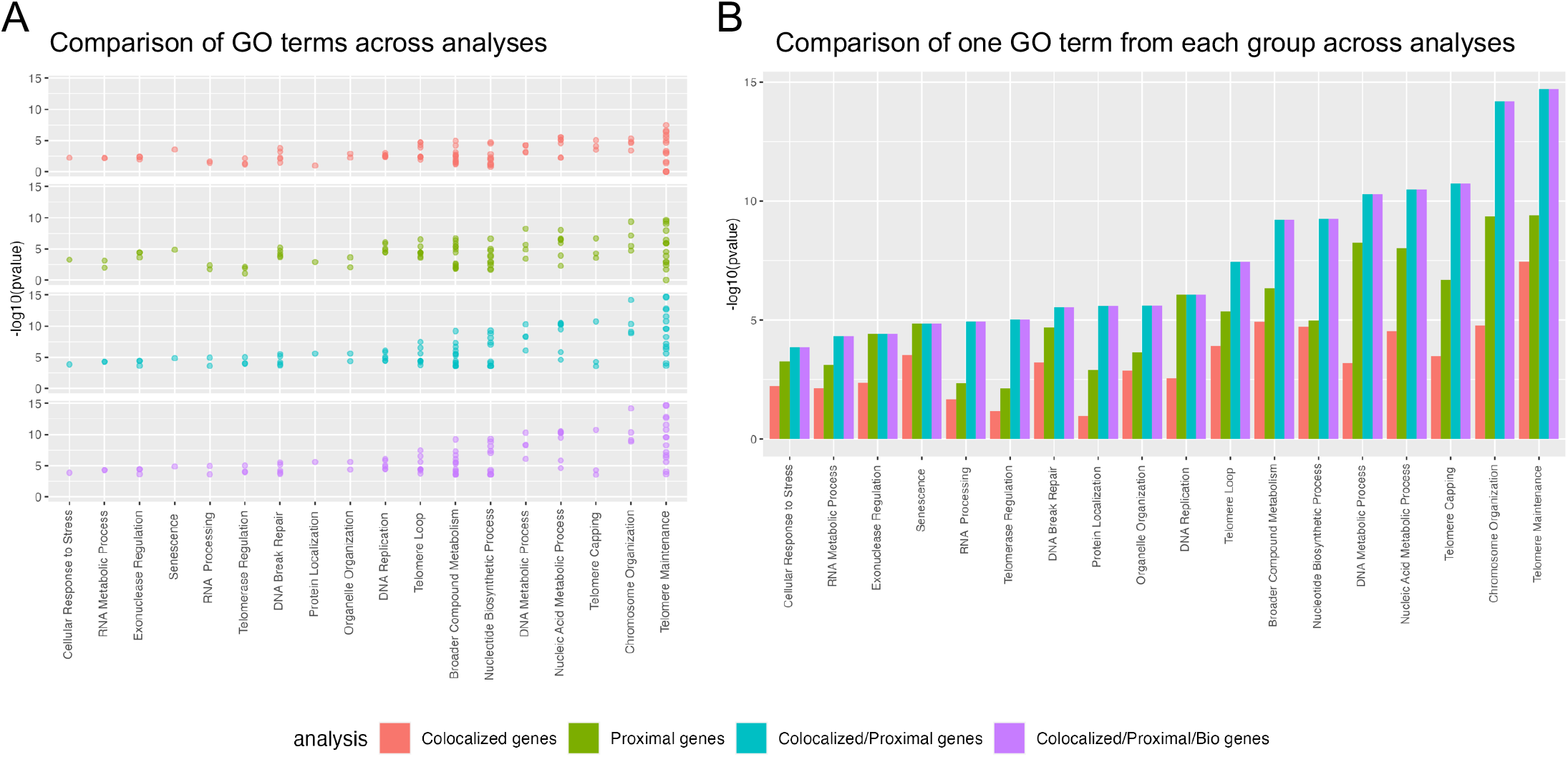
Comparison of GO enrichment analysis results with different gene input datasets. Each meta-analysis locus was assigned a gene based on genes indicated by colocalization analysis (red), the proximal gene (green), genes indicated by colocalization analysis where possible and proximal genes where not possible (blue), or genes indicated by proximity-plus-knowledge, colocalization analysis, or proximal genes where no other information was available (purple). In the fourth case (purple) there were five loci where a nearby gene has known roles in telomere length regulation but was neither the proximal gene nor the gene indicated by colocalization analysis (further explored in the Supplemental Note). Note that there are more genes included in the proximal gene list than the colocalized gene only list as every meta-analysis signal has a proximal gene but not all have colocalization results. GO terms were manually grouped based on related biology and the GO term with the smallest p-value in the Colocalized+Proximal+Bio analysis was chosen as a representative of the group in the plot. Group assignments and comparison of enrichment for all GO terms with FDR < 0.05 are reported in Supplementary Table 8. A. All GO terms that had FDR < 0.05 in at least one analysis are shown. B. The GO term with the smallest pvalue in the Colocalized+Proximal+Bio analysis was chosen for each group and the comparison of pvalues across analyses are shown.

**Supplementary Figure 3:**
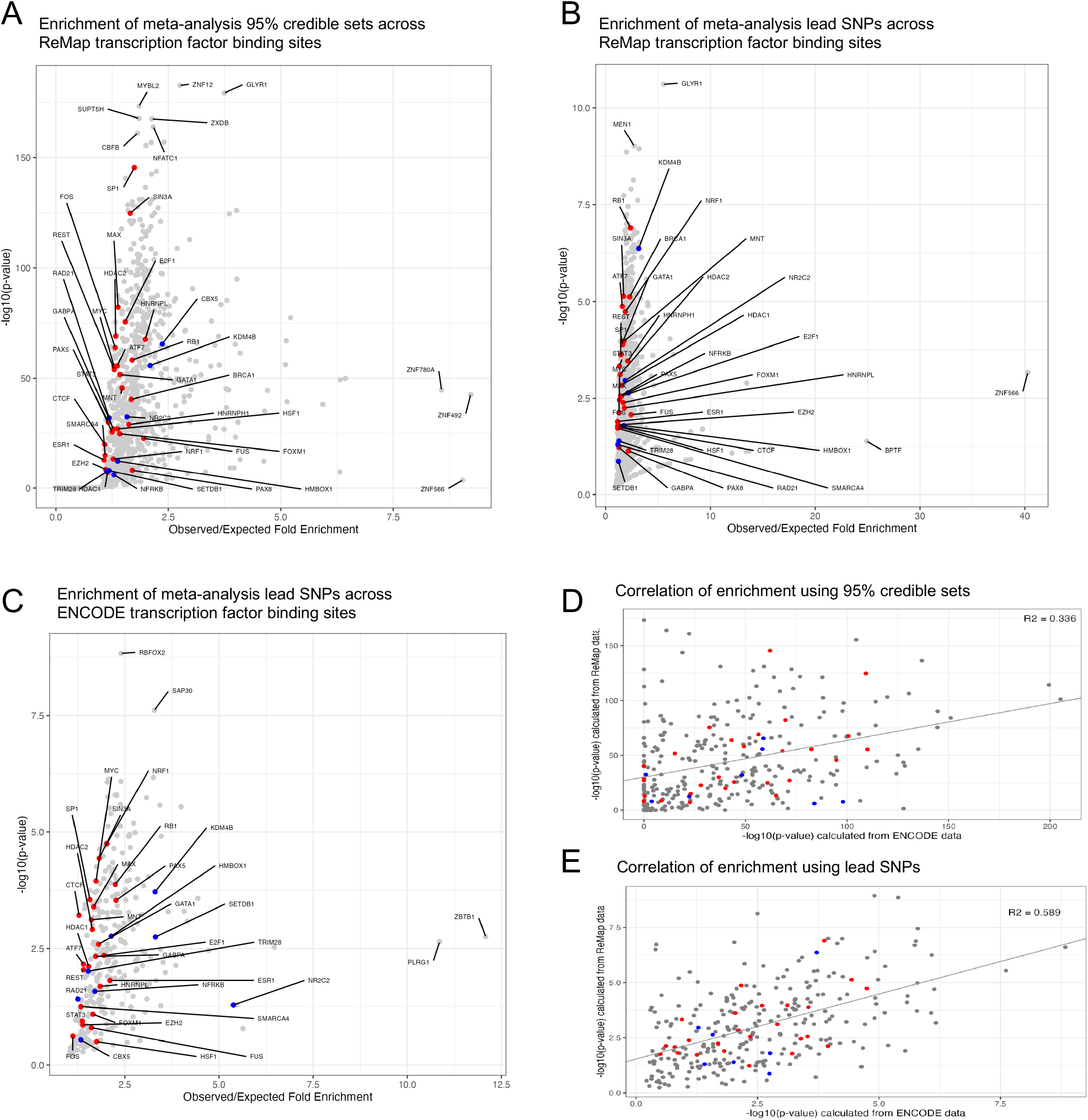
Meta-analysis signals are enriched for transcription factor binding sites of transcription factors with roles in telomere length regulation. The red points represent transcription factors with known roles in regulating telomere length regulation genes and the blue points represent transcription factors with known roles in the alternative telomere lengthening (ALT) pathway. A. The enrichment of 95% credible set SNPs across all transcription factors with data available from ReMap data (Methods). There were 176 transcription factors that fell at the (0,0) coordinate and are not shown for clarity; one (XRCC3) had known roles in ALT. B. The enrichment of only the lead SNP at each meta-analysis signal across all transcription factors with data available from ReMap data (Methods). There were 196 transcription factors that fell at the (0,0) coordinate and are not shown for clarity; one (XRCC3) had known roles in ALT. C. The enrichment of only the lead SNP at each meta-analysis signal across all transcription factors with data available from ENCODE data (Methods). There were 22 transcription factors that fell at the (0,0) coordinate and are not shown for clarity; one (XRCC3) had known roles in ALT. D-E. The enrichment of transcription factors included in both the ReMap and ENCODE datasets are shown. The grey line represents the regression between these two variables and the R2 is shown in the top right corner.

**Supplementary Figure 4:**
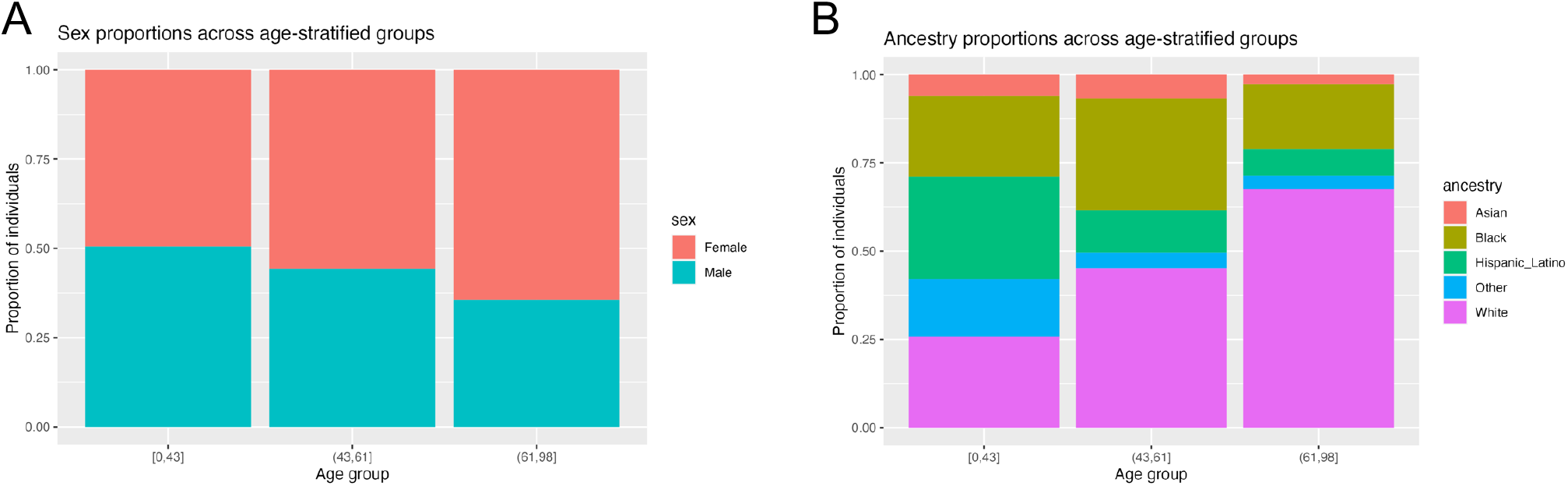
Demographics for age-stratified telomere length GWAS. The 109,122 TOPMed individuals with telomere length estimates (Taub, et al. 2022) were divided into three age groups such that there was a similar number of individuals per group. There were 36,980 individuals in the [0,43] group, 37,470 individuals in the (43,61] group, and 34,671 individuals in the (61,98] group. A. The proportion of individuals of each biological sex in each age group. B. The proportion of individuals of different ancestries in each age group. Ancestry was previously determined computationally (Taub et al. 2022).

**Supplementary Figure 5:**
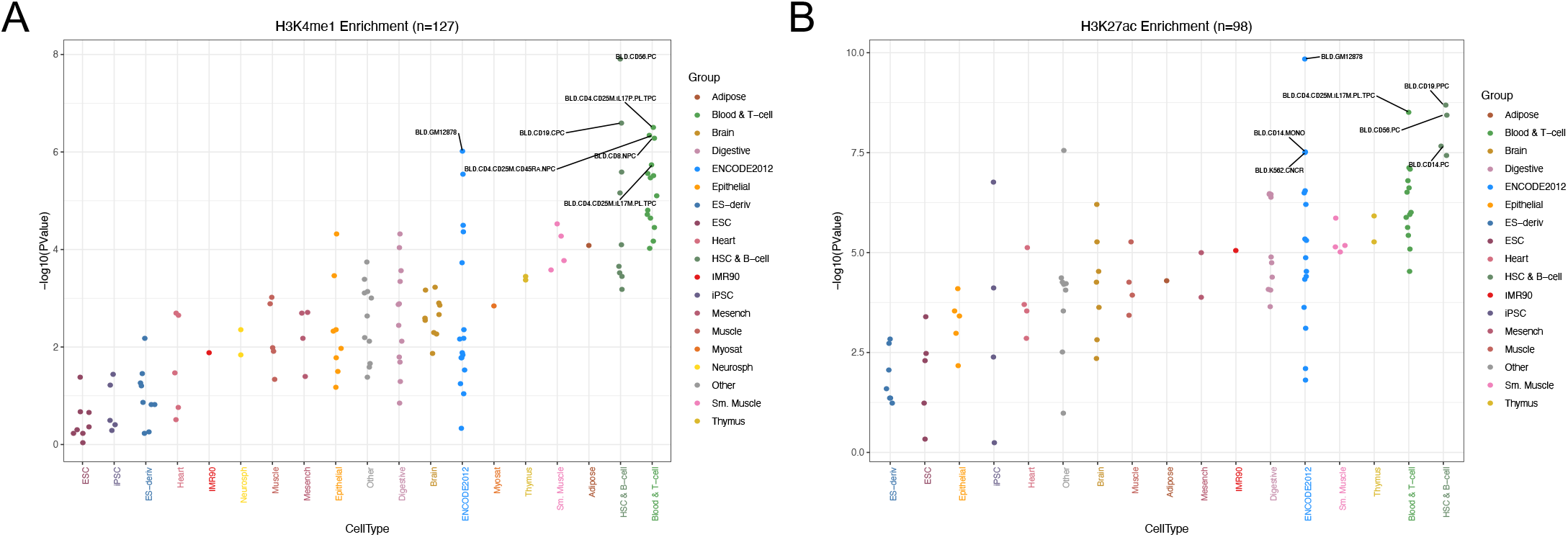
ChIP-seq signals for specific chromatin marks from Roadmap Epigenomics across cell types. Enrichment of Roadmap cell types for sentinel SNPs in H3K4me1 (A) or H3K27ac (B) peaks across 127 and 98 cell types, respectively. Included samples are listed in Supplementary Table 13.

**Supplementary Figure 6:**
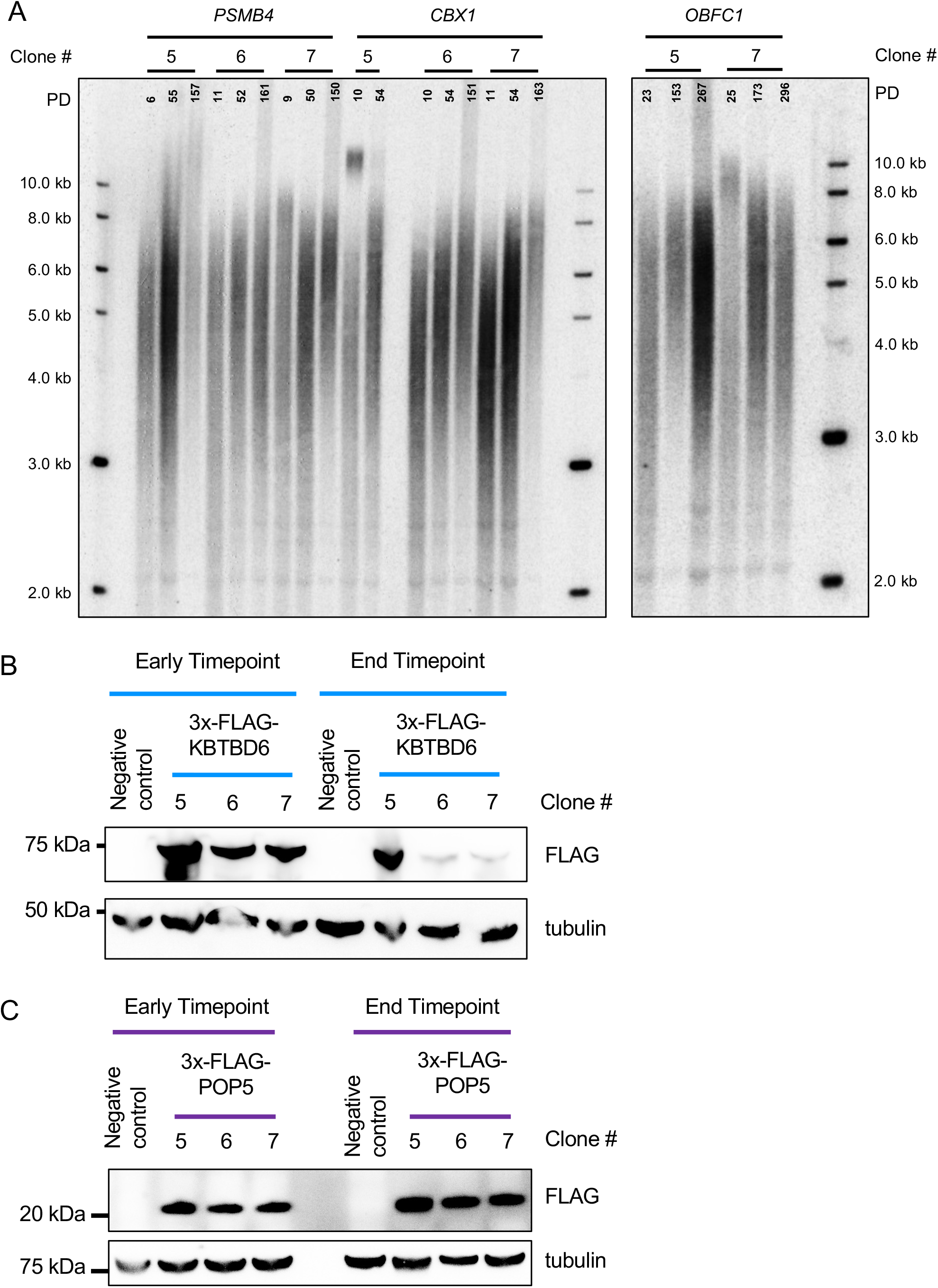
Control data for overexpression of *KBTBD6* and *POP5*. A. *PSMB4*, *CBX1*, or *OBFC1* was constitutively overexpressed from the CMV promoter in HeLa-FRT cells using the FLP-in system. Telomere Southern blots showing the bulk telomere length from a population of cells following an approximate normal distribution. Molecular weight standards were run alongside the samples and their size is indicated in kilobases (kb). Three time points are shown for each clone and the estimated number of population doublings (PD) for each timepoint are indicated. All transfection experiments began from the same population of HeLa-FRT cells. B. KBTBD6 overexpression was maintained in clone 5 over time but was lost in clones 6 and 7 as demonstrated by the end timepoint. The early timepoint was passage 8 of the experiment, approximate population doublings were: clone 5 = 51, clone 6 = 33, clone 7 = 45. The end timepoint was passage 31, approximate population doublings were: clone 5 = 273, clone 6 = 257, clone 7 = 274. C. POP5 overexpression was maintained across all three clones. The early timepoint was passage 8 of the experiment, approximate population doublings were: clone 5 = 65, clone 6 = 59, clone 7 = 67. The end timepoint was passage 31 of the experiment, approximate population doublings were: clone 5 = 296, clone 6 = 297, clone 7 = 318.

**Supplementary Figure 7:**
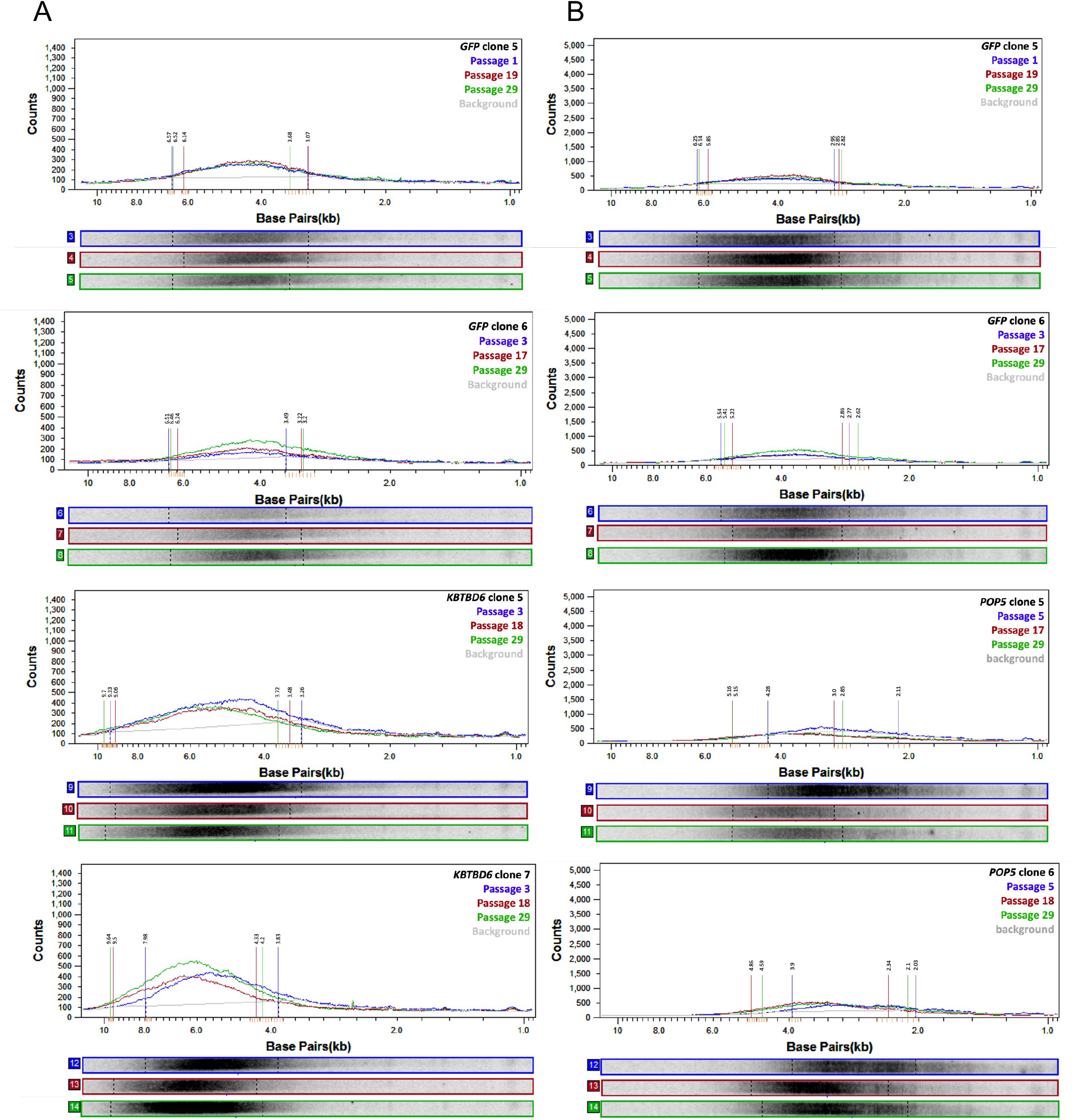
ImageQuant TL estimation of minimum, median, and maximum telomere length. Unprocessed scans of the telomere Southern blots were imported into ImageQuant TL and median telomere length was calculated taking molecular weight markers on either side of the Southern into account. The median telomere length was automatically estimated as the maximum value in these line plots for each line. Line plots were generated for the three time points (timepoints indicated by line color) for each clone. The grey lines indicate the background signal estimated by ImageQuant TL. The Southern blot lanes analyzed in each plot are shown below their respective line plots. The software indicates the range of the signal that it takes into account when estimating the median and these boundaries (dotted lines on the lanes) were used to represent the minimum and maximum telomere lengths. The vertical lines on the line plot were added manually and colored to match the sample they estimate, the values above them represent the estimated minimum or maximum. The software does not provide a quantitative estimate of these boundaries and so we inferred them from the units on the x-axis. Where the minimum or maximum did not fall close to an automated tick mark, we imputed additional tick marks (orange) by anchoring two lines on the available tick marks and adding another three lines in between, then distrubted evenly horizontally using Microsoft PowerPoint. A. Line plots from Figure 6A. B. Line plots from Figure 6B.

**Supplementary Figure 8:**
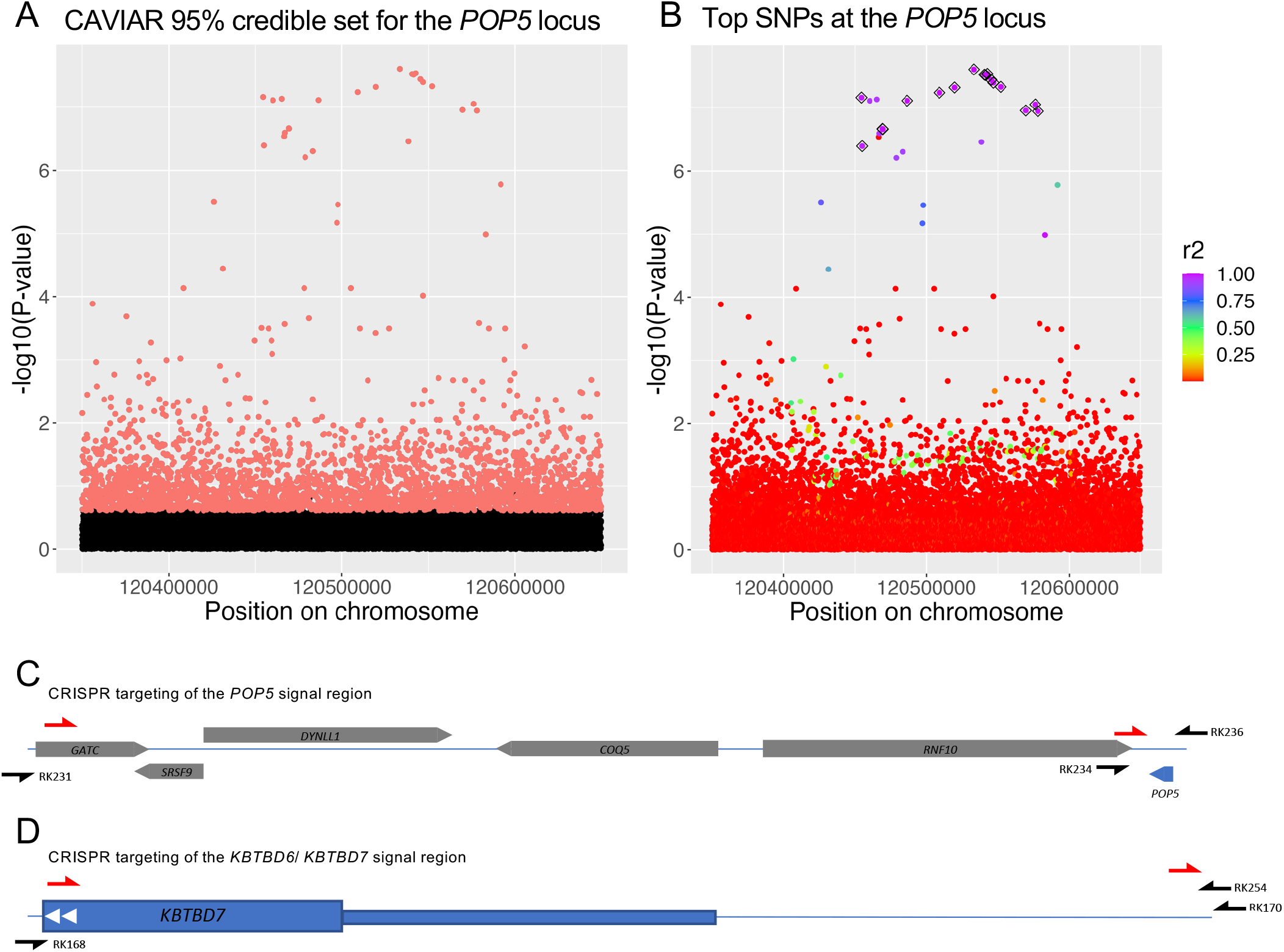
CRISPR/Cas9 targeted regions. A. Manhattan plot showing the association signal near *POP5*. Red SNPs were in the CAVIAR 95% credible set. CAVIAR was run assuming there was one causal SNP in the signal (c=1). B. Manhattan plot showing the association signal near *POP5*. Color indicates linkage disequilibrium (r^2^) calculated with respect to the lead SNP. C. 124 kb region targeted for CRISPR/Cas9 editing within the *POP5* association signal region. The red half arrows indicate the position of CRISPR/Cas9 gRNA sequences. The black half arrows indicate the position of primers used to genotype CRISPR/Cas9-edited cells (Methods). Primer and guide sequences are reported in Supplementary Table 14. The position and size of the indicated coding sequences were taken from the UCSC genome browser and are to scale. *POP5* is indicated in blue. D. 938 bp ATAC-seq peak region targeted for CRISPR/Cas9 editing within the *KBTBD6*/ *KBTBD7* association signal region. The red half arrows indicate the position of CRISPR/Cas9 gRNA sequences. Primer and guide sequences are reported in Supplementary Table 14. The position and structure of the *KBTBD7* coding sequence was taken from the UCSC genome browser and is to scale.

